# The genomes of medicinal skullcaps reveal the polyphyletic origins of clerodane diterpene biosynthesis in the family Lamiaceae

**DOI:** 10.1101/2022.09.25.509435

**Authors:** Haixiu Li, Song Wu, Ruoxi Lin, Yiren Xiao, Ana Luisa Malaco Morotti, Ya Wang, Tao Huang, Yong Zhao, Xun Zhou, Jun Yang, Qing Zhao, Angelos K. Kanellis, Cathie Martin, Evangelos C. Tatsis

## Abstract

The presence of anticancer clerodane diterpenoids is a chemotaxonomic marker for the traditional Chinese medicinal plant *Scutellaria barbata*, although the molecular mechanisms behind clerodane biosynthesis are unknown. Here, we report a high-quality assembly of the 414.98 Mb genome of *S. barbata* into thirteen pseudochromosomes. Using phylogenomic and biochemical data, we mapped the plastidial metabolism of kaurene (gibberellins), abietane and clerodane diterpenes in three species of the family Lamiaceae (*Scutellaria barbata, Scutellaria baicalensis* and *Salvia splendens*), facilitating the identification of genes involved in the biosynthesis of the clerodanes, kolavenol and isokolavenol. We show that clerodane biosynthesis evolved through recruitment and neofunctionalization of genes from gibberellin and abietane metabolism. Despite the assumed monophyletic origin of clerodane biosynthesis which is widespread in species of the Lamiaceae, our data show distinct evolutionary lineages and suggest polyphyletic origins of clerodane biosynthesis in the family Lamiaceae. Our study not only provides significant insights into the evolution of clerodane biosynthetic pathways in the mint family, Lamiaceae, but also will facilitate the elucidation of anticancer clerodanes biosynthesis and future metabolic engineering efforts to increase the production of these high-value chemicals.

## Introduction

Plants of the family Lamiaceae (mints) are used widely for their culinary, cosmetic and medicinal applications (Mint Evolutionary Genomics Consortium, 2018). Scutellaria is the second largest genus of the mint family and its members, widely known as skullcaps, have been used extensively for applications in Traditional Chinese Medicine (TCM) with fifty Scutellaria species recorded in TCM remedies (Shen et al., 2021; Zhao et al., 2016). *Scutellaria barbata* D. Don (barbed skullcap) is used in TCM treatments of cancers, particularly advanced metastatic cancers, where the efficacy of *S. barbata* extracts at reducing cancer progression as well as the absence of harmful side effects has resulted in renewed interest in using this TCM prescription (□□□ - bàn-zhī-lián) as a therapy complimentary to modern chemotherapies (Feng et al., 2021; Li et al., 2016; Marconett et al., 2010; Perez et al., 2010; Tomlinson et al., 2022). Many of the anticancer activities of *S. barbata* extracts have been attributed to the clerodane diterpenoids (Li *et al*., 2016; Wang et al., 2020a; Wang et al., 2020b; Wang et al., 2019; Yang et al., 2018; Yuan et al., 2017), and we have shown that the major clerodane diterpenoid, scutebarbatine A which is produced in specialized glandular trichomes on the leaves of *S. barbata* (Tomlinson *et al*., 2022), induces the apoptosis of Caco-2 cancer cells (human colon adenocarcinoma cell line). Scutebarbatine A promotes apoptosis specifically in cancer cells by reducing the abundance of the inhibitors of apoptosis proteins (Tomlinson *et al*., 2022). There are more than 150 different clerodanes isolated and structurally identified from the aerial parts of *S. barbata* (Table S1) (Chen et al., 2020; Li *et al*., 2016; Wang *et al*., 2020a) although the biosynthesis of clerodane diterpenes in *S. barbata* has not been described.

Clerodanes are a class of bicyclic diterpenes and their structural scaffold is a rearranged labdane carbon skeleton involving the 1,2 shift of two methyl groups (Fig. S1) (Li *et al*., 2016). Like other plant diterpenoids such as gibberellins, the first steps in biosynthesis of clerodane diterpenoids occur in plastids, where the common precursor for most of the plant diterpenoids, geranylgeranyl diphosphate (GGPP) is produced (Figure 1). GGPP is usually subject to paired catalytic activity by class II and class I diterpene synthases (diTPS) to form a polycyclic scaffold (Figure 1) (Andersen-Ranberg et al., 2016; Jia et al., 2019; Jia et al., 2016). Diterpene scaffolds are exported from plastids and they generally undergo oxidations catalysed by cytochrome P450 enzymes (Bathe and Tissier, 2019; Christianson, 2017; Zhou and Pichersky, 2020). Isotopologue analysis of the carbon skeleton of the clerodane, salvinorin A, after labelling with ^13^C CO_2_ and ^13^C glucose in *Salvia divinorum* plants (Fig. S1), demonstrated that the clerodane scaffold originates from the plastidial methylerythritol 4-phosphate (MEP) pathway (Kutrzeba et al., 2007; Ostrozhenkova, 2008). The first dedicated step in the biosynthesis of clerodanes has been identified in a few plants from the mint family such as *S. divinorum* (Chen et al., 2017; Pelot et al., 2017), *Vitex-angus castus* (Heskes et al., 2018), *Ajuga reptans* (Johnson et al., 2019), *Callicarpa americana* (Hamilton et al., 2020) and involves the class II diTPSs, kolavenyl diphosphate synthase (KPS) (Chen *et al*., 2017; Heskes *et al*., 2018; Pelot *et al*., 2017) or isokolavenyl diphosphate synthase (IKPS) (Johnson *et al*., 2019), which transform GGPP to kolavenyl diphosphate (KPP) and isokolavenyl diphosphate (IKPP), respectively (Chen *et al*., 2017; Hamilton *et al*., 2020; Johnson *et al*., 2019; Pelot *et al*., 2017) (Figure 1). The proposed mechanism for class II clerodane synthases begins with the initial protonation and cyclization of GGPP for the formation of the intermediate *ent*-labda-13*E*-en-8-yl diphosphate carbocation; a Wagner-Meerwein rearrangement takes place with two subsequent 1, 2 hydride and methyl shifts resulting to the generation of the clerodane scaffold with the formation of kolav-13E-en-4-yl diphosphate carbocation, and a final proton abstraction results in the formation of either KPP or IKPP (Figure S1) (Jia *et al*., 2019; Jia *et al*., 2016; Lemke et al., 2019; Potter et al., 2016). No class I diTPSs have yet been described which act on KPP or IKPP in a native pathway (Chen *et al*., 2017; Kwon et al., 2022; Pelot *et al*., 2017), limiting the elucidation of clerodane diterpenoid biosynthesis in the mint family.

**Figure 1.**
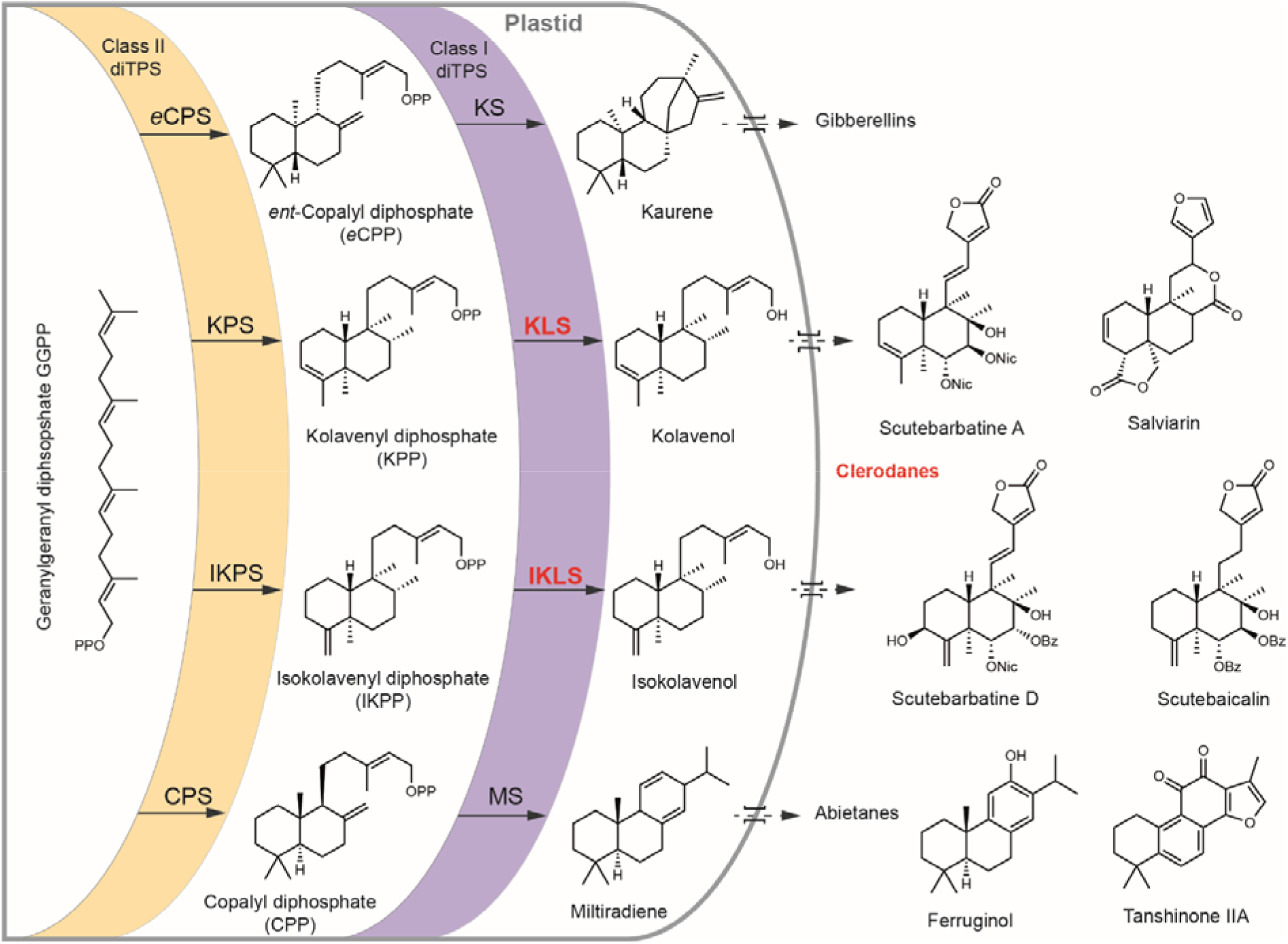
Schematic overview of the biosynthesis of the major diterpene scaffolds (kaurene, kolavenol, isokolavenol and miltiradiene) synthesized in *Scutellaria barbata, Scutellaria baicalensis* and *Salvia splendens*. Kaurene serves as a precursor of gibberellins, kolavenol and isokolavenol serve as precursors of clerodanes (such as scutebarbatine A, scutebarbatine D in *S. barbata*), and scutebaicalin (in *S. baicalensis*), salviarin (in *S. splendens*) and miltiradiene serves as the precursor of the abietane diterpene, ferruginol.

Like gibberellins that have been isolated from both plants (He et al., 2020) and fungi (Tudzynski, 2005), clerodanes have been reported from several hundred different plant species (Li *et al*., 2016; Merritt and Ley, 1992), as well as fungi (Andersen et al., 1983), bacteria (Nakano et al., 2015) and marine animals (Lhullier et al., 2019). Although clerodanes have been isolated from plants of the Asteraceae, Euphorbiaceae, Flacourtiaceae, and Menispermeaceae families, more than 50% of reported clerodanes have been isolated from members of the family Lamiaceae (Li *et al*., 2016; Merritt and Ley, 1992) and these have attracted considerable attention for their biological activities (Li *et al*., 2016). Some of the clerodanes produced by Scutellaria and Ajuga genera are considered potent insect antifeedants (Li *et al*., 2016); while the clerodanes from *Salvia divinorum*, like salvinorin A (Fig. S1), are used as opioid receptor probes for the development of new therapeutic tools for drug addiction (Hill et al., 2020; Li *et al*., 2016). Because clerodanes have been isolated and identified in genera from all Lamiaceae subfamilies (Table S2) (Li *et al*., 2016; Merritt and Ley, 1992), and there are no reports of clerodanes in other families of the order Lamiales (Li *et al*., 2016; Merritt and Ley, 1992), it has been widely assumed that clerodane biosynthesis had a monophyletic origin in the Lamiaceae.

To understand how clerodane diterpenes are synthesized in *S. barbata* and how the clerodane biosynthetic pathway evolved in the family Lamiaceae, we sequenced the genome of *S. barbata* and mapped diterpene metabolism by performing biochemical characterization of all the diterpene synthases from *S. barbata* and two other species from the family Lamiaceae, *Scutellaria baicalensis* (Zhao et al., 2019b) and *Salvia splendens* (Dong et al., 2018) combined with analysis of their phylogenetic and syntenic relationships. Our work offers new insight into the origins and evolution of diterpene synthases in clerodane biosynthesis in the family, Lamiaceae.

## Results

### *Scutellaria barbata* genome sequencing, assembly, and annotation

DNA was extracted from a single *Scutellaria barbata* plant growing in Shanghai Chenshan Botanical Garden. DNA was sequenced employing Illumina, Pacbio, Bionano and Hi-C sequencing techniques. Based on k-mer analysis of Illumina data the genome size was estimated as 414.98 Mb. We obtained 52.26 Gb total data by Illumina Novaseq achieving 157x sequence coverage, and respectively by PacBio 135.29 Gb (326x), by Bionano 265.2 Gb (640x) and by Hi-C 68.93 Gb (166x) (Table S3). We used Illumina paired-end short read sequencing to survey the genome. The Pacbio long-read sequencing was used for genome assembly while Illumina data were used for error correction of contigs. Bionano genome imaging was used to scaffold the contigs resulting in 376.96 Mb with a total number of 396 scaffolds and N50 value 14.97 Mb (Table S4). A Hi-C (*in vivo* fixation of chromosomes) library was then employed to refine the first version of the reference genome, utilizing additionally LACHESIS for improved results (Fig. S2, Table S4). This improved the assembly slightly to give 376.97 Mb sorted into 356 super scaffolds, with a super-scaffold N50 of 26.00 Mb with the longest super-scaffold being 44.71 Mb (Table S5). The super scaffolds could be aligned into 13 groups (Table S6), hereafter referred to as pseudochromosomes, which represented an assembly of 90.84% of the estimated genome size. The genome of *S. barbata* has a GC content of 33.84% with N comprising 0.20% (Table S7); while SNP calling based on the genome sequence revealed a heterozygosity rate of 0.0132% (Fig. S3; Table S8). The coverage of the final version of the genome assembly was tested by mapping the Illumina short reads, which resulted in 96.50% mapping rate and 99.78% overall genome coverage (Table S9). Expressed sequenced tags (ESTs) from RNA sequencing were also mapped on the assembled genome; 89.85% of ESTs aligned more than 90% of their sequences in one scaffold, while 99.33% of ESTs aligned more than 50% of their sequences in one scaffold (Table S10). CEGMA (Core Eukaryotic Genes Mapping Approach) (Table S11) and BUSCO (Benchmarking Universal Single-Copy Orthologs) (Table S12; Fig. S4) evaluations of the genome sequence indicated 95.56% and 98.50% completeness, respectively, suggesting, overall, a high-quality assembly. A pipeline combining *de novo* predictions, homology-based predictions, and RNA sequencing data was used to construct gene models for the *S. barbata* genome. A total of 25,899 genes were annotated this way, with an average length of 3,065 bp and an average coding sequence length of 1,199 bp (Table S13). Long terminal repeats (LTRs) of retroelements were the most abundant interspersed repeat, accounting for 42.80% of the genome, followed by DNA transposable elements at 3.53% (Table S14). Genes annotated as encoding non-coding RNAs (ncRNAs) in the current genome included 623 microRNAs (miRNAs), 566 transfer RNAs (tRNAs), 1599 ribosomal RNAs (rRNAs), and 513 small nuclear RNAs (snRNAs) (Table S15). An overview of the genes, repeats, non-coding RNA densities, and all detected segmental duplications is presented in Figure 2. Functional annotation for the gene sets was obtained by searching against publicly available databases, including SwissProt (http://www.uniprot.org), NR (www.ncbi.nlm.nih.gov/protein), Pfam (pfam.xfam.org), KEGG (http://www.genome.jp/kegg) InterPro (www.ebi.ac.uk/interpro). The best match from each database was assigned to the protein-coding gene.

**Figure 2.**
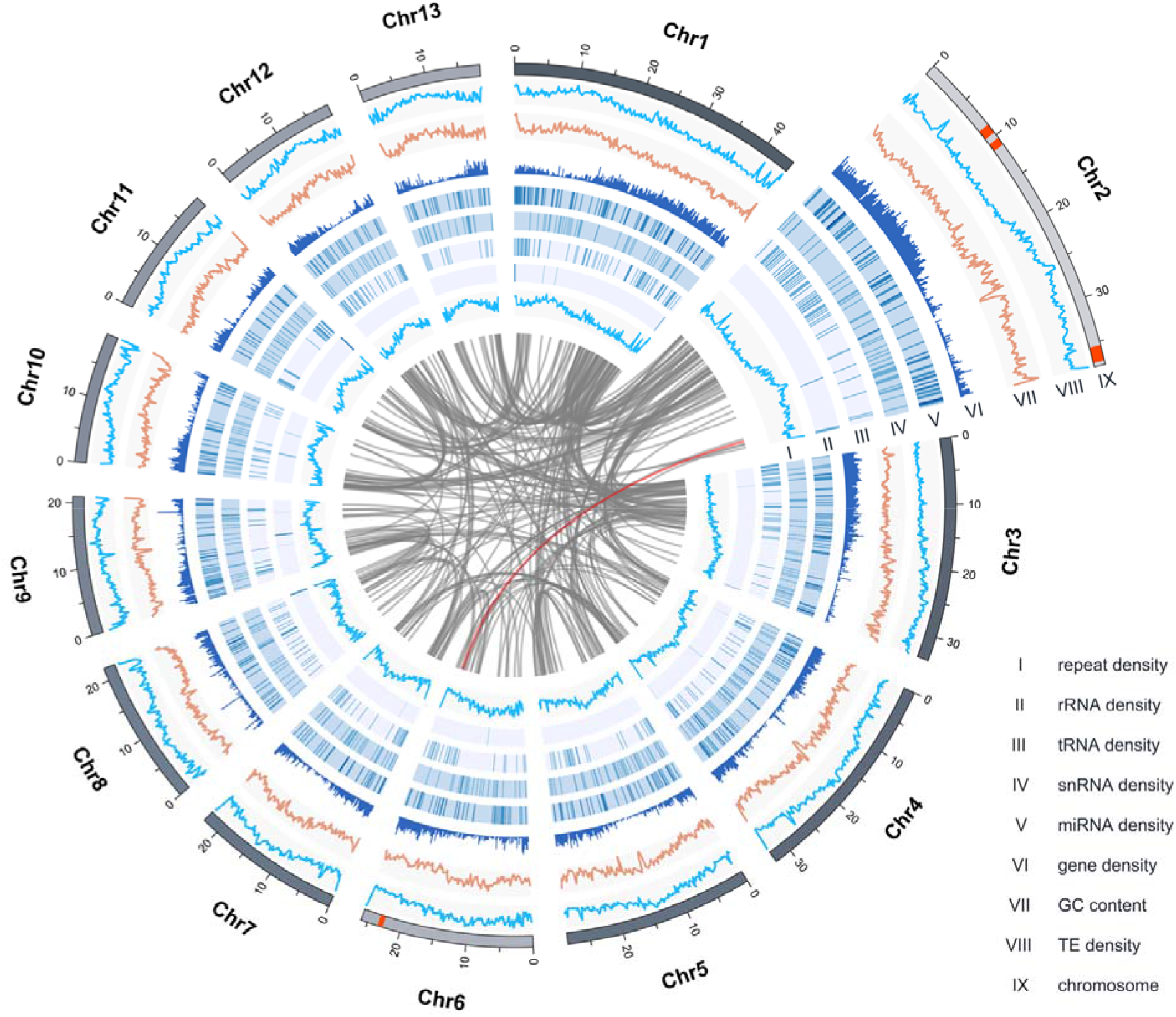
Overview of *Scutellaria barbata* draft genome assembly. The CIRCOS plot represents the genomic features of the *S. barbata* genome (pseudomolecule size 414.98 Mb). From inside to outside: the grey links indicate intraspecies syntenic relationships; the Roman numerals refer to: I, simple sequence repeat (SSR) density in a 100kb sliding window; II, rRNA density in a 100kb sliding window; III, tRNA density in a 100kb sliding window; IV, snRNA density in a 100kb sliding window; V, miRNA density in a 100kb sliding window; VI, gene density in a 100kb sliding window; VII, GC content density in a 100kb sliding window; VIII transposon elements (TE) density in a 100kb sliding window; and IX the outer lines represent pseudochromosomes. The red blocks in pseudochromosomes Chr2 and Chr6 indicate the position of diterpene synthases (Fig. S10), and the link highlighted in red illustrates their syntenic relationships.

We compared our genome assembly for *S. barbata* with 15 other sequenced genomes, including three species of the family Lamiaceae with high quality sequenced genomes (*Salvia splendens, Salvia miltiorrhiza, Scutellaria baicalensis*), three species of the order Lamiales (*Antirrhinum majus, Andrographis paniculata, Sesamum indicum*), two euasterid species (*Solanum lycopersicum, Coffea canephora*), four eurosid species (*Vitis vinifera, Glycine max, Arabidopsis thaliana, Populus trichocarpa*), a monocot (*Oryza sativa*); with *Nymphaea colorata* and *Amborella trichopoda* representing a sister group at the base of the angiosperms (Fig. S5). We identified 28370 gene families, of which 6706 were shared by all 16 species, and 284 of these gene families consisted of single-copy orthologous genes. In *S. barbata*, we identified expansion of 926 gene families and contractions of 1934 gene families. The 284 single-copy orthologous genes were used to construct a phylogenetic tree (Fig. S6). The resulting phylogenetic tree showed that the plants of the family Lamiaceae split from the rest of the Lamiales lineage around 57 Million Years Ago (MYA) as calculated for *S. indicum*, while the genera Scutellaria and Salvia diverged around 45 MYA. The species *S. barbata* and *S. baicalensis* diverged around 16 MYA (Fig. S6). The branching order in the tree based on genome comparisons is consistent with previously proposed phylogenetic orders (Consortium, 2018; Dong *et al*., 2018; Xu et al., 2016; Zhao *et al*., 2019b).

We compared the complexity of gene families between *S barbata* (Lamiaceae), *A majus* (Lamiales), *C canephora* (Lamiids), *A thaliana* (eurosids) and *O sativa* (monocots). A maximum of 9064 gene families were shared between all the five species while 750 gene families were shared between the four eudicot species and 749 gene families were specific to *S. barbata* (Fig. S7). Gene ontology enrichment analysis showed significant enrichments of genes with the functional annotation “terpene synthase activity” and “terpenoid biosynthetic process” in the *S. barbata* genome (Fig. S8) suggesting a pivotal role of terpenoid metabolism in the physiology of *S. barbata*.

Intragenomic and self-alignment analysis between and within the chromosomes revealed paralogous relationships among the 13 *Scutellaria barbata* chromosomes with 258 major duplications (Figure 2). Collectively *S. barbata* has 5059 pairs of paralogous genes. Based on calculation of the density distribution of synonymous substitution rate per gene (Ks) between the collinear paralogous genes, a peak at around Ks=0.8 (Fig. S9) indicated a whole genome duplication (WGD) event around 61.5 million years ago. Our estimate of the timing of the WGD observed in *S. barbata* is consistent with what has been proposed *for S. baicalensis* (Zhao *et al*., 2019b). We also examined the synteny between the genomes of *S. barbata* and *S. baicalensis* and we found 90 syntenic blocks shared by the two species of the genus, Scutellaria (Fig. S10, S11). Although, the two Scutellaria species share a lot of common morphological traits, they show significant differences in diterpene metabolism. While more than 150 clerodane diterpenoids have been isolated from *S. barbata* (Table S1) (Chen et al., 2020; Dai et al., 2006; Li et al., 2016; Merritt and Ley, 1992; Wang et al., 2020a), there is only one report of a clerodane (scutebaicalin) isolated from *S. baicalensis* (Hussein et al., 1996).

Clerodane diterpenoids are widespread throughout the family Lamiaceae with some considered to be chemotaxonomic markers for some species (Fontana et al., 2006; Johnson *et al*., 2019; Li *et al*., 2016; Merritt and Ley, 1992; Roncero et al., 2018). Genomic data therefore provide the unique opportunity to investigate the evolution of a specialized metabolic pathway within the mint family.

### Identification and characterization of diterpene synthases

Mining the genome of *S. barbata* by matching the amino acid sequences of the annotated genes with the relevant Hidden Markov Models (HMM) protein families of terpene synthases (TPS) (PF01397 & PF03936) (Mistry et al., 2020), we constructed a terpene synthase (TPS) dataset and used these protein sequences to build a TPS phylogenetic tree. Using similar methods, we mined the genomes of *S baicalensis, S. splendens (Dong et al*., *2018; Fontana et al*., *2006)* and teak, *Tectona grandis (Macías et al*., *2010; Zhao et al*., *2019a)* which are all members of the Lamiaceae which make clerodane diterpenes, and we compared these proteins to diTPSs from *Solanum lycopersicum* which does not make clerodanes (Fig. S12). The phylogenetic tree defined different clades of class II and class I diterpene synthases, based on homology to characterized diterpene synthases from species of the family Lamiaceae. To check that we were identifying active diterpene synthases (diTPS), we searched for the aspartic acid-rich active site domains, DXDD for class II diTPSs and DDXXD for class I diTPSs (Chen et al., 2011; Zerbe and Bohlmann, 2015). By this filtering, we identified 8 class II diTPSs and 4 class I diTPS in *S. barbata* (Table 1).

**Table 1.** **Summary of characterized activities of class I and class II diterpene synthases and cytochrome P450 enzymes identified from the three species of Lamiaceae (*S. barbata, S. baicalensis* and *S. splendens*).**

| Enzymatic activity | Plant species |  |  |
| --- | --- | --- | --- |
| Class II diTPS | *Scutellaria barbata* | *Scutellaria baicalensis* | *Salvia splendens* |
| **ent-Copalyl diphosphate synthase (eCPS)** | SbbdiTPS2.2 [N.b.]* | SbdiTPS2.5, SbdiTPS2.6 [N.b.] | SspdiTPS2.3, SspdiTPS2.6 [N.b.] |
| **Copalyl diphosphate synthase (CPS)** | SbbdiTPS2.4, SbbdiTPS2.5 [N.b.; E.c.] | SbdiTPS2.1, SbdiTPS2.2, SbdiTPS2.3 [N.b.] | SspdiTPS2.4, SspdiTPS2.5 [N.b.; S.c.] |
| **Kolavenyl diphosphate synthase (KPS)** | SbbdiTPS2.1 [N.b.; E.c.] | - | SspdiTPS2.1 [N.b.; E.c.] |
| **Isokolavenyl diphosphate synthase (IKPS)** | SbbdiTPS2.3 [N.b.; E.c.] | SbdiTPS2.7, SbdiTPS2.8 [N.b.; E.c.] | - |
| **Non identified activity** | SbbdiTPS2.6, SbbdiTPS2.7, SbbdiTPS2.8 [N.b.] | SbdiTPS2.4, SbdiTPS2.9, SbdiTPS2.10 [N.b.] | SspdiTPS2.2, SspdiTPS2.7, SspdiTPS2.8 [N.b.] |
| Class I diTPS |  |  |  |
| **Kaurene synthase (KS)** | SbbdiTPS1.3 [N.b.] | SbdiTPS1.2 [N.b.] | SspdiTPS1.1, SspdiTPS1.2 [N.b.] |
| **Miltiradiene synthase (MS)** | SbbdiTPS1.1 [N.b.; E.c.] | SbdiTPS1.1 [N.b.] | SspdiTPS1.3, SspdiTPS1.4 [N.b.; S.c.] |
| **Kolavenol/ Isokolavenol synthase (KLS/IKLS)** | SbbdiTPS1.2, SbbdiTPS1.4 [N.b.; E.c.] | SbdiTPS1.3 [N.b.; E.c.] | SspdiTPS1.1, SspdiTPS1.2, SspdiTPS1.3, SspdiTPS1.5 [N.b.; E.c.] |
| Cytochrome P450 |  |  |  |
| **Ferruginol synthase (FS)** | SbbCYP2, SbbCYP3, SbbCYP4 [S.c.] | SbCYP1, SbCYP2, SbCYP3, SbCYP4 [S.c.] | SspCYP2, SspCYP3, SspCYP4 [S.c.] |

\*In brackets are denoted the host organism used for heterologous protein expression; N.b. for *N. benthamiana*; E.c. for *E. coli*; S.c. for *Saccharomyces cerevisiae*.

To identify which genes encode the first steps of the clerodane biosynthetic pathway in *S. barbata*, it was necessary to characterize the activity of the encoded diterpene synthases, functionally. Since the majority of clerodane diterpenoids have been reported from plants of the family Lamiaceae, and we wanted to understand the microevolution of clerodane diterpene metabolism, we decided to characterize the diTPSs from *S. baicalensis* and *S. splendens* functionally (Table 1), in addition to those from *S. barbata*. The identified class II and class I diTPSs from the three investigated species, were expressed in *Nicotiana benthamiana* and *E. coli* to identify their enzymatic products. We used GCMS for the analysis of enzyme assays both *in planta* and *in vitro* and the MS spectra were analysed using the NIST17 database. As controls, we synthesized the characterized genes, *TwTPS14* from *Tripterygium wilfordii* which has kolavenyl diphosphate synthase activity (KPS) (Andersen-Ranberg *et al*., 2016) and *ArTPS2* from *Ajuga reptans* which has isokolavenyl diphosphate synthase activity (IKPS) (Johnson *et al*., 2019), and we expressed both *in planta* and *in vitro* and used their enzymatic products as standards. Diterpene metabolism occurs predominantly in plastids using precursors from the methyl-erythritol pathway (MEP) (Chen *et al*., 2011; Zerbe and Bohlmann, 2015), and both class II and class I diTPSs have plastid transit peptides. The transit peptide for each diterpene synthase protein was removed based on ChloroP predictions (Emanuelsson et al., 1999) and truncated versions of each protein were expressed in *E. coli*.

We expressed class II diTPSs in *N. benthamiana* (Lindbo, 2007), yeast (Božić et al., 2015) and *E. coli* (Berrow et al., 2007) and our findings are summarized in Table 1. The heterologously expressed and purified proteins from *E. coli* were assayed *in vitro* against geranylgeranyl diphosphate (GGPP). Based on GCMS analysis of both types of assay, we established that the genes from *S. barbata, SbbdiTPS2*.*2, S. baicalensis, SbdiTPS2*.*5* and *SbdiTPS2*.*6*, and *S. splendens, SspdiTPS2*.*3* and *SspdiTPS2*.*6* encode enzymes with *ent*-copalyl diphosphate synthase (*e*CPS) activity (Fig. S13) and the genes *SbbdiTPS2*.*4* and *SbbdiTPS2*.*5*, from *S. barbata, SbdiTPS2*.*1, SbdiTPS2*.*2* and *SbdiTPS2*.*3* from *S. baicalensis*, and *SspdiTPS2*.*4* and *SspdiTPS2*.*5* from *S. splendens*, encode enzymes with copalyl diphosphate synthase (CPS) activity (Fig. S14-S16). The enzymes SbbdiTPS2.1 from *S. barbata* and SspdiTPS2.1 from *S. splendens* showed kolavenyl diphosphate synthase activity (KPS) (Figure 3, Fig. S17) with a characteristic MS fragment at *m/z* 189 while the enzymes SbbdiTPS2.3 from *S. barbata* and SbdiTPS2.7 and SbdiTPS2.8 from *S. baicalensis* demonstrated isokolavenyl diphosphate synthase activity (IKPS) (Figure 3, Fig. S17) with a characteristic fragment at *m/z* 191. Enzymes with KPS and IKPS activities, have been reported to catalyse the first dedicated step in biosynthesis of clerodanes (Chen *et al*., 2017; Li *et al*., 2016; Pelot *et al*., 2017). There were also a number of class II diTPSs from each of the three species that did not show any activity *in planta or in vitro* towards GGPP (Table 1) (Fig. S14-S16).

**Figure 3.**
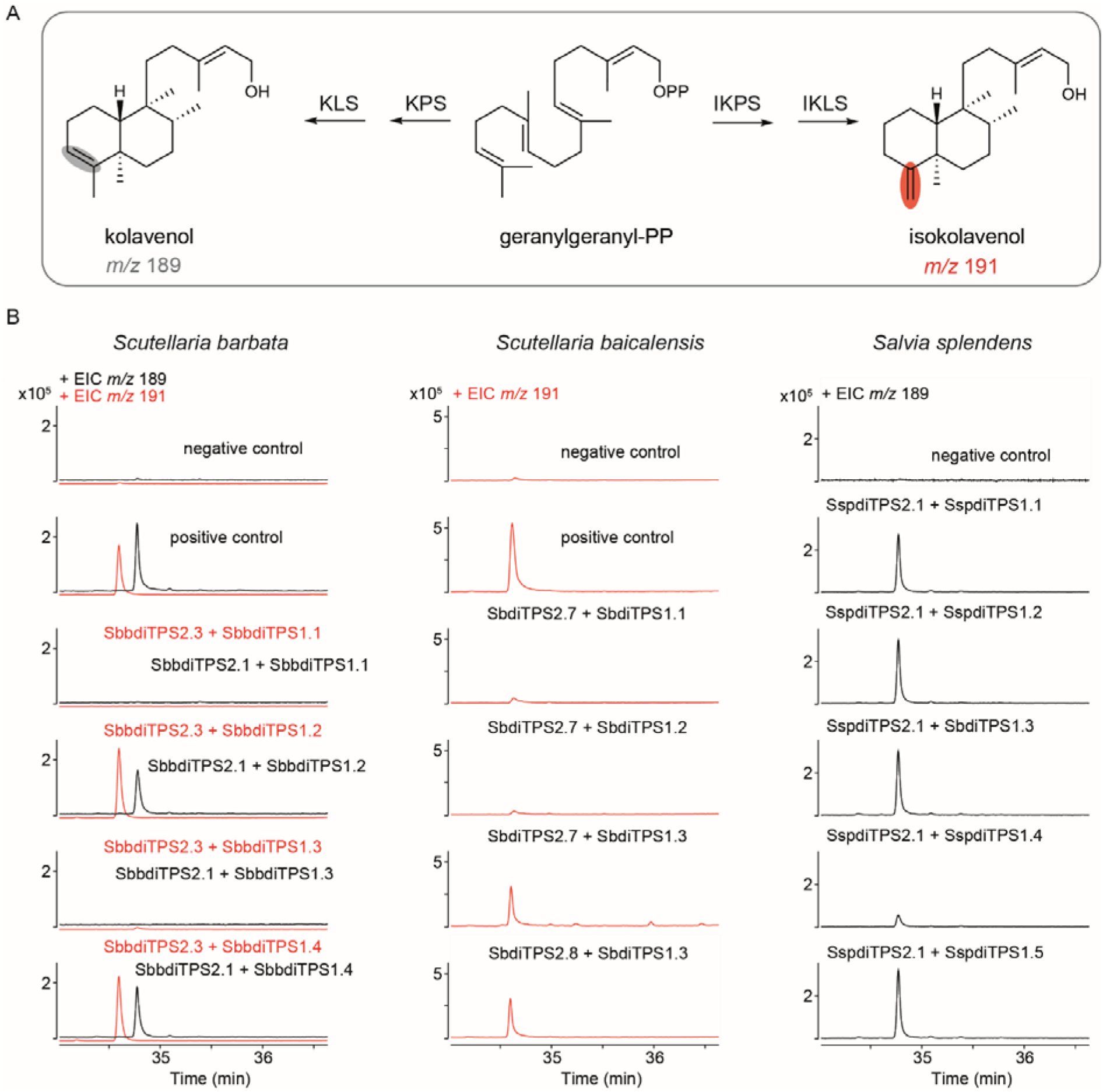
Identification of enzymes involved in clerodane diterpene biosynthesis in *S. barbata, S. baicalensis* and *S. splendens*. **A**, kolavenol and isokolavenol biosynthetic pathway in *S. barbata, S. baicalensis* and *S. splendens*: GGPP; KPS; IKPS; KLS; IKLS. **B**, GCMS analysis (selected *m*/*z* signals: 189 and 191) of extracts from enzyme assays expressing class II diTPS or combinations of class II and class I diTPSs. Negative control: enzyme assays expressing class II diTPS without alkaline phosphatase treatment. Positive control: enzyme assays expressing class II diTPS treated with alkaline phosphatase.

Since class I diTPSs display substrate promiscuity towards a broad range of enzymatic products of class II diTPSs (Jia *et al*., 2019; Jia *et al*., 2016), we combined the different class I diTPSs with the active class II diTPSs from *S. barbata* and the other Lamiaceae species either in *N. benthamiana* or *in vitro* (*E. coli* expressed) to characterize the enzymatic activity of the class I diTPSs. The co-expression of SbbdiTPS1.3, SbdiTPS1.2, SspdiTPS1.1, and SspdiTPS1.2 with the characterized *e*CPSs from the same species resulted in the synthesis of kaurene, and therefore these enzymes were assigned as kaurene synthases (KS) (Figure 3, Fig. S13, Table 1). The coupling of SbbdiTPS1.1, SbdiTPS1.1, SspdiTPS1.3 and SspdiTPS1.4 with the CPS enzymes from the same species, showed miltiradiene synthase (MS) activity (Figure 3, Fig. S14-16, Table 1) and were therefore assigned as miltiradiene synthases.

Both the SbbdiTPS1.2 and SbbdiTPS1.4 class I diTPSs of *S. barbata*, were able to convert the KPP (product of SbbdiTPS2.1 enzyme) as well as the IKPP (product of SbbdiTPS2.3 enzyme) to kolavenol and isokolavenol, respectively (Figure 3). These class I diTPSs catalyse the synthesis of either kolavenol or isokolavenol and represent activities involved in the second step in the clerodane biosynthetic pathway (Kwon *et al*., 2022; Muchlinski et al., 2021). Heterologously expressed (*E. coli*) and purified SbdiTPS1.3, a class I diTPS from *S. baicalensis*, catalysed the transformation of IKPP to isokolavenol in coupled enzyme assays with SbdiTPS2.7 and SbdiTPS2.8 (Figure 3). In the coupled assays of SspdiTPS2.1 (KPS) with class I diTPSs from *S. splendens*, SspdiTPS1.1, SspdiTPS1.2, SspdiTPS1.3, and SspdiTPS1.5 all showed kolavenol synthase (KLS) activity (Figure 3).

By combining different class II and class I diterpene synthases, the functional characterization of diTPSs allowed us to map diterpene biosynthesis in three species. The combined activities of class II and class I diTPSs lead to the formation of three distinct diterpene structural scaffolds: *ent*-kaurene, miltiradiene and kolavenol-isokolavenol (clerodanes) as summarized in Table 1. In *S. barbata* we identified two class II clerodane synthases, SbbdiTPS2.1 with KPS activity and SbbdiTPS2.3 with IKPS activity, and two class I clerodane synthases SbbdiTPS1.2 and SbbdiTPS1.4 with KLS and IKLS activities, respectively.

### Expression of genes encoding diTPSs in *S. barbata*

We have shown that the clerodanes of *S. barbata* accumulate mainly in leaves (Tomlinson *et al*., 2022). Targeted LCMS metabolomics showed (Tomlinson *et al*., 2022) that leaves and flowers produce most scutebarbatine A, the major clerodane in *S. barbata*. MALDI imaging showed that scutebarbatine A accumulates in large glandular trichomes on the leaves, and this was confirmed by isolation and LCMS analysis of glandular trichomes (Tomlinson *et al*., 2022). We measured the expression of all the characterised *S. barbata* diTPSs across different tissues (roots, young and old stems, young and old leaves, flowers, trichomes and trichome depleted leaves) by qRT-PCR (Fig. S18). Expression of the kaurene pathway *diTPSs* was constitutive and the *diTPSs* of miltiradiene pathway were expressed most highly in roots. The clerodane *diTPSs* appeared to have the highest expression in flowers and glandular trichomes from leaves with *SbbdiTPS1*.*4* showing unique expression in these trichomes. The expression profiles of clerodane *diTPSs* coincided with the presence of clerodanes across different tissues based on our analytical data (Tomlinson *et al*., 2022) (Fig. S18).

### Evolution of clerodane biosynthetic pathway in the family Lamiaceae

#### Clerodane biosynthesis in the family Lamiaceae is result of positive selection

We constructed phylogenetic trees for class II proteins (Fig. S19) using the diTPSs from *S. barbata, S. baicalensis, S. splendens* and the diTPSs that have been characterised biochemically from other species of the family Lamiaceae available from NCBI. The phylogenetic tree of class II diTPSs (Figures 4A, S19) showed two major clades, one with diterpene synthases encoding CPS activity and a second with *e*CPS activity. The class II clerodane synthases represent an emerging subclade from the *e*CPSs. Important in the phylogenetic relationships between *e*CPS and KPS is the catalytic site of *e*CPS where mutations of histidine (H) to phenylalanine (F) or tyrosine (Y) in the LLXSL motif of this active site, result in transition of *e*CPS activity to KPS and vice versa (Karunanithi and Zerbe, 2019; Lemke *et al*., 2019; Pelot *et al*., 2017; Potter *et al*., 2016). We analysed further the *e*CPS/KPS clade of class II diTPS to investigate the role of positive selection in emergence of class II clerodane synthases (Fig. S20). In principle, the ratio of non-synonymous base pair substitutions (mutations) K_a_ to synonymous base pair substitutions (mutations) K_s_ of each codon can define whether a gene is under positive (K_a_/K_s_ >> 1), neutral (K_a_/K_s_ ∼ 1) or purifying selection (K_a_/K_s_ << 1) (Lichman et al., 2020b; Wu et al., 2022; Yang and Nielsen, 2008). We found that the branch (highlighted in red in Fig. S20) was under positive selection with the calculated ratio K_a_/K_s_=9.0215 and a value for P=7.25×10^−3^, suggesting that the evolution of *e*CPS to KPS or IKPS activity in the family Lamiaceae was the result of positive selection.

**Figure 4.**
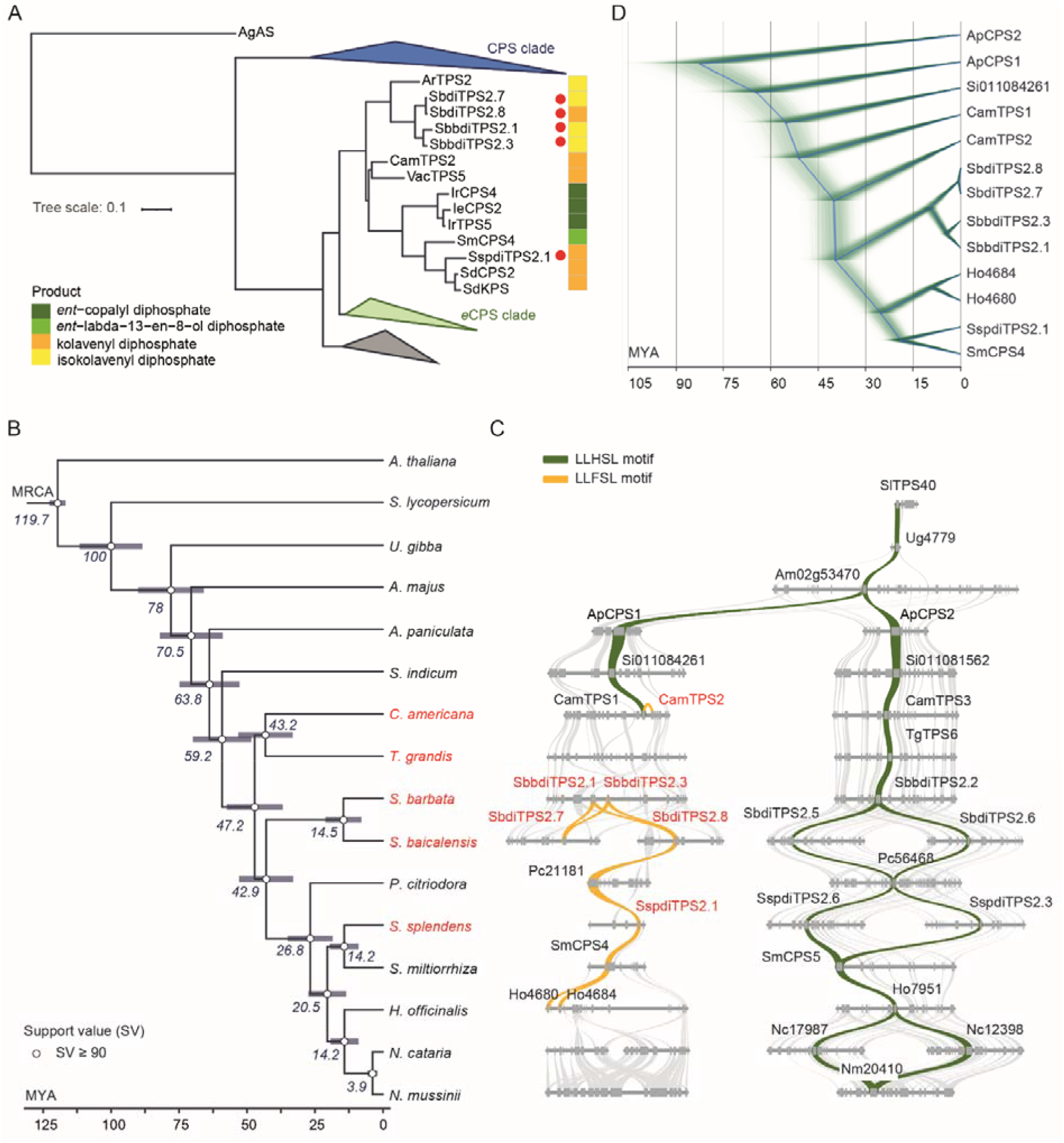
Phylogenetic and syntenic analyses of class II diTPSs from the order Lamiales indicating changes in the active site motif. **A**, Phylogenetic tree of characterized class II diterpene synthases from Lamiaceae species focused on class II clerodane synthases. The enzymes characterized in the present study are denoted by the red dots. The products produced by class II diTPSs from the substrate, GGPP, are shown in boxes with different colours. The full version is shown in Fig. S19. **B**, Phylogram based on whole genome analyses of species from the family Lamiaceae and the order Lamiales. In red are highlighted the species producing clerodanes. **C**, Syntenic analysis of loci encoding *e*CPS and KPS/IKPS from species of the family Lamiaceae and the order Lamiales. The gold ribbon shows the syntenic relationships between the diterpene synthases with the LLFSL active site motif, while the green ribbon links the diterpene synthases with the LLHSL active site motif associated with *e*CPS activity. The class II clerodane synthases are labelled with red font. **D**, Bayesian inference tree for class II clerodane synthases based on the calculated species divergence time (panel B). A more detailed view is shown in Fig. S26.

Syntenic analysis of genomes can trace genomic inheritance and variance across species and uncover intra-species duplication events related to pathway diversification (Franke et al., 2019; Li et al., 2021; Wu *et al*., 2022; Zhao *et al*., 2019b). To understand better the homology, genomic organization and phylogenetic relationships between the different activities and functionalities of diTPSs, we performed syntenic analysis using the genomes of *S. barbata, S. baicalensis, S. splendens, S. miltiorhiza, T. grandis* and *A. paniculata*. We used *A. paniculata* as an “outgroup reference” from the order Lamiales because it is rich in diterpenoid metabolism (Sun et al., 2019). *S. miltiorrhiza* is a well-known medicinal sage producing tanshinones, a class of diterpenoids derived from miltiradiene, but there are no reports of it producing clerodanes (Wang and Peters, 2022). There are reports of simple clerodanes in *teak (T. grandis)* (Macías *et al*., 2010), and diTPSs from *A. paniculata* (Sun *et al*., 2019), *T. grandis* (Zhao *et al*., 2019a) and *S. miltiorrhiza* (Cui et al., 2015; Su et al., 2016) have been characterised biochemically.

#### Class II clerodane synthases originated from an ancient *e*CPS gene duplication

Syntenic analysis of the loci encoding either class II KPS or IKPS activities revealed that the *KPS*/*IKPS* genes from *S. barbata, S. baicalensis* and *S. splendens* are syntenic (Fig S21). In *S. barbata* a tandem duplication gave rise to SbbdiTPS2.1 and SbbdiTPS2.3 followed by subfunctionalization to KPS and IKPS activities respectively. In *S. baicalensis*, the two copies of the *IKPS* genes are more distant ∼ 1.5 Mb apart (Fig S21) suggesting a larger, independent duplication. The class II *KPS/IKPS* genes are syntenous to *SmCPS4* which encodes *ent*-ladbenyl diphosphate synthase activity (Cui *et al*., 2015) and to ApCPS1 which encodes *e*CPS activity (Sun *et al*., 2019). We extended our comparative genome analysis using genomic data from more species in the family Lamiaceae and the order Lamiales. This extended syntenic analysis showed that the genes from *A. paniculata, ApCPS1* and *ApCPS2* encoding *e*CPS activity (Sun *et al*., 2019) are both syntenous to the gene *Am02g53470* (annotated as *e*CPS) from *A. majus* (Li et al., 2019) and *SlTPS40* from *S. lycopersicum* encoding *e*CPS activity (Falara et al., 2011) (Figure 4C). These findings led us to explore the phylogenomic relationships and extend the syntenic analysis to 15 species, supported by a phylogenetic tree with the estimated divergence times for each species (Figure 4B).

Figure 4C shows two, distinct syntenic blocks of class II diTPSs: on the right are genes encoding enzymes with *e*CPS activity with a LLHSL motif in the active site of each enzyme, highlighted by the green ribbons, On the left, the gold ribbon highlights the class II diTPSs containing an LLFSL motif in their active sites, including those with KPS/IKPS activity. The genes with *e*CPS function (or are annotated as such), are organized in a highly conserved genomic region in all species (Figures 4B, S22). The phylogenetic tree and the synteny suggest a time of between 70.5 - 63.8 MYA between the divergence of *A. majus* and *A. paniculata*, involving either a WGD event or a large segmental duplication which resulted in a duplicated region with an additional *eCPS* gene. We cloned the gene Si011084261 (Figure 4C) from *Sesamum indicum* and expressed it in *N. benthamiana*, but neither *e*CPS nor KPS activity were detected (Fig. S23). The enzymes we characterized functionally, either as KPS or IKPS, are encoded by genes that are organized in the duplicated region and are indicated in red font in Figure 4C. The golden ribbon shows the syntenic relationships between class II diTPSs with the LLFSL motif in their active sites (Figure 4C, Fig. S25, S26). Several reports (Lemke *et al*., 2019; Potter *et al*., 2016) (Pelot et al., 2019; Pelot *et al*., 2017) have suggested that mutation of histidine (H) to phenylalanine (F) or tyrosine (Y) in the LLXSL motif of the active site of *e*CPS, switches the catalytic activity from *e*CPS to KPS (Potter *et al*., 2016). However, the existence of SmCPS4 with *ent*-labdenol diphosphate synthase activity suggests that the LLFSL motif is necessary but not sufficient for KPS/IKPS enzymatic activities. The syntenous genes, Ho4680 and Ho4684 from *H. officinalis* were cloned and expressed, but no catalytic activity towards GGPP was detected (Fig. S24), possibly due to “corrupted” aspartic acid-rich active sites (DXDD: which in the case of Ho4680 and Ho4684 are EIDN and DIDV, respectively). No syntenous genes were identified in two Nepeta species (*N. cataria* and *N. mussini*).

The combination of functional characterization of enzymes with phylogenetic and genomic analysis revealed that the *KPS/IKPS* genes in Lamiaceae evolved from a second copy of *e*CPS which was the result of this ancient WGD or large segmental duplication, before the divergence of the Lamiaceae from other families in the order Lamiales. This was supported by the functional characterization of the proteins CamTPS1 and CamTPS2 from *Callicarpa americana* which have *e*CPS and KPS activities, respectively (Hamilton *et al*., 2020). From the Bayesian phylogenetic tree (Zhao *et al*., 2019b) of class II diTPSs in this duplicated genomic area (Figures 4D, S26), the divergence of CamTPS2 (KPS) from CamTPS1 (*e*CPS) is estimated to have taken place around 51.3 MYA, before the split of *C. americana* from the genus Scutellaria around 47.2 MYA (Figure 4A). This suggests that evolution of KPS activity in Callicarpa and Scutellaria was monophyletic. Phylogenetic analysis indicated that the Scutellaria class II diTPSs are closely related to CamTPS2 (Figures 4A, D, S19). Although, clerodanes have been isolated from *T. grandis* (Macías *et al*., 2010) our syntenic analysis did not show any diTPS in the syntenic region, which could imply an independent origin of clerodane synthesis in *T. grandis*. In *S. barbata*, the class II clerodane synthase activity most likely evolved by neofunctionalization of *e*CPS, followed by a tandem duplication (estimated around 4.4 MYA) and subsequent subfunctionalization of SbbdiTPS2.1 and SbbdiTPS2.3 for KPS and IKPS activity, respectively. In the clade of Lamiaceae (mostly including plants from Nepetoideae subfamily) that diverged 26.8 MYA, there are examples of loss of activity (*H. officinalis*) and loss of diTPS genes (*N. cataria & N. mussini*). The activity of the enzyme from the common ancestor of SspdiTPS2.1 and SmCPS4 remains unclear; either this had *e*CPS activity (with the LLFSL motif) and SspdiTPS2.1 represents an additional late neofunctionalization acquiring KPS activity, or SmCPS4 has regained *ent*-labdane diphosphate synthase activity despite having the LLFSL motif, from an ancestor with KPS activity. Altogether, the phylogenomic data do not support a monophyletic origin of class II diTPSs involved in clerodane biosynthesis in the family Lamiaceae, with suggestions of independent neofunctionalizations in Tectona, Callicarpa/Scutellaria and possibly in Salvia (Figure 4A, D).

#### Class I clerodane synthases have polyphyletic evolutionary origins

In the phylogenetic tree of characterized class I diTPSs from the family Lamiaceae (Figure 5A), there are two major clades: the first consists of the enzymes that act on *ent-*copalyl diphosphate (*e*CPP), predominantly with kaurene synthase activity, and the second contains the enzymes that act mainly on copalyl diphosphate (CPP), usually with miltiradiene synthase activity. The newly characterized enzymes with KLS/IKLS activity, SbbdiTPS1.2, SbbdiTPS1.4 and SbdiTPS1.3, from the two species of Scutellaria belong to the clade of enzymes within the class I diTPSs that act on CPP. Although in *S. splendens* the dedicated KLS SspdiTPS1.5 is placed in the clade with other class I diTPS that act on CPP, it is phylogenetically distant (Figure 5A) from the promiscuous miltiradiene synthase SspdiTPS1.3 which also shows KLS enzymatic activity, and from the KLS/IKLS enzymes from Scutellaria species. The paired activity of class II and class I diTPS in plastids, is essential for the generation of kolavenol and later utilization by downstream enzymes (De La Peña and Sattely, 2021; Kwon *et al*., 2022). The phylogeny of class I clerodane synthases supports the polyphyletic origin of clerodane biosynthetic pathway in the family Lamiaceae.

**Figure 5.**
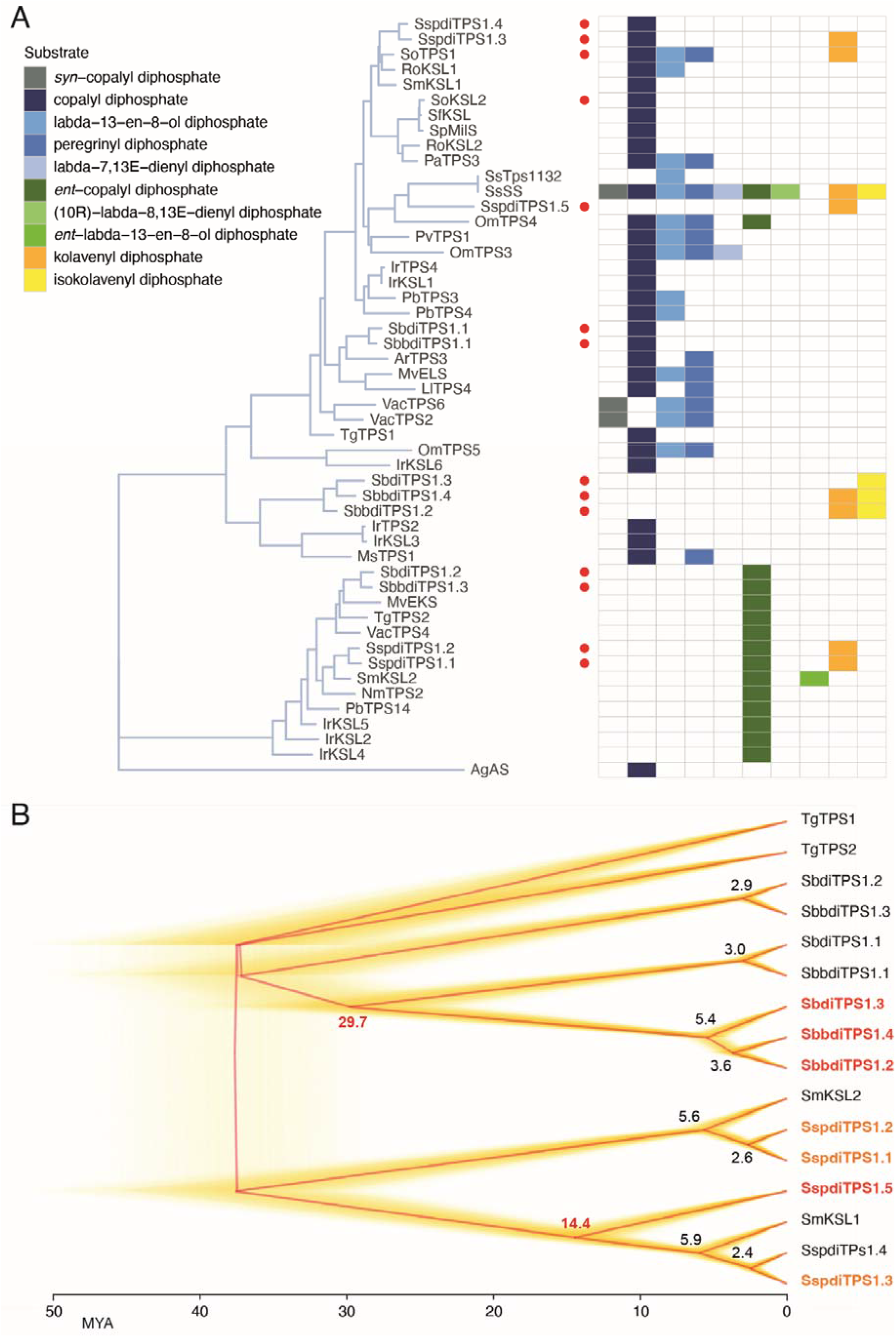
Phylogeny of class I diTPS. **A**, Phylogenetic tree of characterized class I diterpene synthases from Lamiaceae species. The phylogeny was built from class I diTPSs from the family Lamiaceae (Table S20) and those characterized in this study (highlighted with red dots) using 1000 bootstrap values. Abietadiene synthase from *Abies grandis* (AgAS) was used as the outgroup. The different substrates used by the class II diTPSs are shown in boxes with different colours. **B**, Bayesian inferred phylograms for class I diTPSs in *S. barbata, S. baicalensis, S. splendens, S. miltiorrhiza* and *T. grandis* showing estimated times of divergence on the X axis in millions of years based on the times calculated for species divergence shown in Figure 4. The runs used for constructing the phylogenetic tree are shown in yellow and the consensus tree is shown in red. The estimated divergence times of genes were calculated based on species divergence times. Genes encoding dedicated KLS enzymes are shown in red while genes encoding proteins showing promiscuity towards KPP in *S. splendens* are shown in orange colour.

A Bayesian inferred phylogeny of class I clerodane synthases (Figure 5B) showed that, in the lineage shared by the two Scutellaria species, the emergence of class I clerodane synthase was the result of tandem replication of MS genes (Figure 6) and that divergence occurred around 29.7 MYA (Figure 5B). The projected time for the duplication event that resulted in SbbdiTPS1.2 and SbbdiTPS1.4 was approximately 3.6 MYA. Consequently, the rise of class II clerodane synthase in Scutellaria lineage was partnered by neofunctionalization of genes encoding MS activity to KLS/IKLS activity and the establishment of the clerodane biosynthetic pathway in Scutellaria spp.

**Figure 6.**
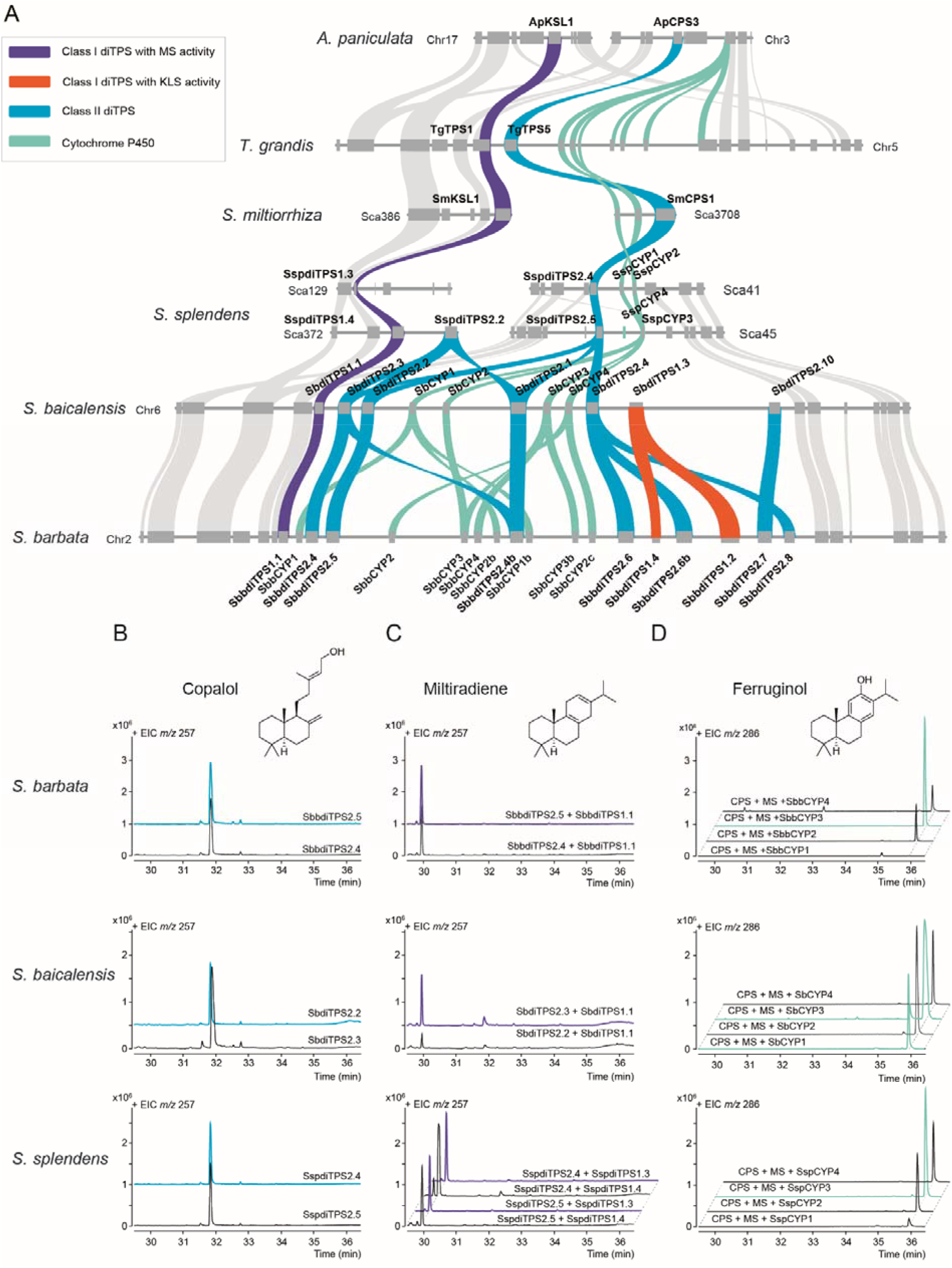
Ferruginol biosynthetic gene cluster in the Lamiaceae. **A**, Genomic region and syntenic analysis among the species of the order Lamiales order; Purple ribbons show the class I diTPSs that act on copalyl diphosphate with MS activity except ApKSL1 which has pimaradiene synthase activity (Sun *et al*., 2019); Light blue ribbons show the class II diTPSs; cyan ribbons show genes encoding cytochrome P450s; and orange ribbons show the newly duplicated class I diTPSs that act in the clerodane pathway as KLS/IKLS. **B-D**, GCMS profile (selected *m*/*z* signals: 257 and 286) of extracts from *Nicotiana benthamiana* leaves expressing class II diTPS **(B)** and combinations of class II and class I diTPSs **(C)** or yeast strains expressing GGPPS (Erg20(F96C)), CPS, MS and FS **(D)**. Three biological replicates of each assay shown in panel D were performed

In contrast, the enzymes catalysing the second step in the clerodane pathway in *Salvia splendens* appear to be result of independent evolutionary events again supporting the polyphyletic origin of clerodane biosynthesis in the Lamiaceae. The split of the KLS gene SspdiTPS1.5 from the clade of class I diTPSs with MS activity is projected to have occurred 14.4 MYA (Figure 5B). To explore whether the substrate promiscuity of class I diTPSs from *S. splendens* with KS activity (SspdiTPS1.1, SspdiTPS1.2) and MS activity (SspdiTPS1.3) towards kolavenyl diphosphate is genus- or species-specific, we assayed the SoTPS1 and SoKSL2 from *Salvia officinalis* against KPP (Figures 5A, S27). Only SoTPS1 showed KLS activity (Fig. S27) indicating that substrate promiscuity and KLS activity are most likely shared among class I diTPS from Salvia species.

Comparative genomic analysis showed the conserved organization of the genomic loci encoding KS activity (Fig. S28). The class I clerodane synthases, SbbdiTPS1.2, SbbdiTPS1.4 and SbdiTPS1.3, from the two Scutellaria species, lie in the same region of the *S. barbata* and *S. baicalensis* genomes (Figure 6A) and they are physically close to the diTPS genes encoding CPS and MS activity. Syntenic analysis revealed that the proximity between the CPS and MS genes was conserved among the genomes of the family Lamiaceae. The physical proximity of the KLS genes to the MS genes in both Scutellaria genomes, indicated that the KLS genes arose as a result of a tandem duplication followed by neofunctionalization of the encoded activities in the lineage common to the two Scutellaria species. In contrast, syntenic analysis revealed a unique insertion of SspdiTPS1.5 in a genomic area which is present only in *S. splendens* (Fig. S29). The phylogeny of class I diTPSs (Figure 5) and their genome organization (Figures 6, S29) suggest that the duplication and neofunctionalization of KLSs occurred after the divergence of the Scutellaria lineage from that of Salvia, reflecting different evolutionary origins of KLS in Scutellaria and Salvia. Consequently, the phylogenomic data from the genes encoding KLS activity support the polyphyletic origin of clerodane biosynthesis in the family Lamiaceae.

#### The genes encoding diterpene synthases in clerodane biosynthesis are not arranged in biosynthetic gene clusters

The organization of genes involved in specialized plant diterpenoid metabolism, including class I & II diTPSs and cytochrome P450s, in the form of biosynthetic gene clusters (BGCs) has been frequently reported (Polturak et al., 2022; Smit and Lichman, 2022; Zhan et al., 2022). In the region of the genome of *S. barbata* where the genes encoding CPS, MS and KLS/IKLS are located, there are also 8 genes encoding cytochromes P450 (CYP), most of them annotated as ferruginol synthase (FS). The evolution of KLS/IKLS activities in Scutellaria species following duplication of MS revealed that BGCs can provide new enzymes for newly emerged metabolic pathways through duplication and neofunctionalization. To explore whether the expansion of cytochrome P450s close to class I clerodane synthases in the ferruginol BGC was associated with new activities involving oxidation of kolavenol as a substrate, we tested the CYPs from *S. barbata* against kolavenol, following expression in yeast. No oxidation activity was observed against kolavenol (Fig. S30), although most of the CYPs showed ferruginol synthase activity against miltiradiene (Figure 6).

## Discussion

*S. barbata* is a medicinal plant, with long standing use in TCM of plant extracts (bàn-zhī-lián) from aerial parts (leaves, stems, flowers) which are rich in clerodanes with more than 150 compounds reported (Table S1) (Chen *et al*., 2020; Li *et al*., 2016; Shen *et al*., 2021; Wang *et al*., 2020a). In our previous work we demonstrated that the major clerodane from leaf trichomes, scutebarbatine A, shows selective cytotoxic activity against cancer cells (Tomlinson *et al*., 2022). *S. baicalensis* is another species of the genus Scutellaria and its medicinal properties have been attributed to 4’-deoxy-flavones from the roots (Zhao *et al*., 2016). Consequently, although the two Scutellaria plants are medicinally important, their therapeutic properties are based largely on different bioactive compounds. Clerodanes are considered as an important chemotaxonomic marker for plants of the family Lamiaceae, since more than half of the reported clerodanes in nature, have been isolated from species of the mint family, mainly from genera Ajuga, Stachys, Teucrium, Salvia and Scutellaria (Frezza et al., 2019; Li *et al*., 2016; Vestri Alvarenga et al., 2001).

Genome sequence provides an essential resource to establish the molecular mechanisms of specialized metabolism (Lichman *et al*., 2020b; Zhao *et al*., 2019a; Zhao *et al*., 2019b). For comparison to the genome of *S. baicalensis* (Zhao *et al*., 2019b), we provide a reference genome of *S. barbata* (414.98 Mb) with high-quality (BUSCO 98.50% completeness) in the subfamily Scutellarioideae, which enriches the genomic information available for the family, Lamiaceae and provides a resource for discovering the relationship between the evolution of chemo-diversity and the evolution of biosynthetic pathways.

We mapped diterpene metabolism by identifying all the genes encoding diterpene synthases in *S. barbata, S. baicalensis*, and *S. splendens*. By characterizing their enzymatic activities, *in vitro* and *in planta*, using the entire set of class II and class I diterpene synthases, we identified three major diterpene biosynthetic pathways: the kauranes which serve as precursors in the synthesis of gibberellins, miltiradiene which serves as a precursor of abietane diterpenes and clerodanes, in these three species. Several class II clerodane synthases, which form either kolavenyl diphosphate or isokolavenyl diphosphate have been reported (Chen *et al*., 2017; Hamilton *et al*., 2020; Heskes *et al*., 2018; Johnson *et al*., 2019; Pelot *et al*., 2017), but class I diTPS acting in the clerodane pathway have not been reported before, despite suggestions that a specialized phosphatase enzyme might be involved in the synthesis of kolavenol (Kwon *et al*., 2022; Pelot *et al*., 2017). The mechanisms of any phosphatase and class I diTPS in kolavenol biosynthesis must be different (Fig. S27). Our holistic approach led to the identification of class I clerodane synthases in all the three species. Our approach overcame deficiencies in functional annotation as in the case of SbdiTPS1.3 (Sb06g19700) from *S. baicalensis*, which has isokolavenol synthase activity, but which was originally annotated as “*uncharacterized protein LOC105975168 from Erythranthe guttatus*”.

Specialized metabolites play important roles in adaptation and survival of plants in different environments including responses to abiotic and biotic stresses, and their presence is taxonomically restricted. Evolution of biosynthetic pathways in the family Lamiaceae is driven mainly by gene duplication, followed by neofunctionalization (Godden et al., 2019; Lichman et al., 2020a; Zhao *et al*., 2019a; Zhao *et al*., 2019b). The majority of clerodanes have been isolated from plants of the family Lamiaceae, but no clerodanes have been reported from other families in the order Lamiales. Consequently, our work sheds new light onto the evolution of clerodane biosynthesis from diterpene metabolism.

Class II clerodane synthases catalyse the first step committed to clerodane biosynthesis, by converting the geranylgeranyl diphosphate to either the kolavenyl or isokolavenyl diphosphate. The enzymes evolved from an *ent*-copalyl diphosphate synthase which acts in gibberellin biosynthesis, and involved a whole genome or segmental duplication that occurred approximately 65 MYA (Figure 4). Although this event was shared between plants of the order Lamiales, neofunctionalization seems to have taken place only in the family Lamiaceae. Within the Lamiaceae, class II clerodane synthases have a phenylalanine in a pocket of the active site (LLFSL) which is necessary for their activity, while the syntenous genes from order Lamiales have a histidine (LLHSL) which confers *ent*-copalyl diphosphate synthase activity.

Class I diTPSs can be considered as metabolic gatekeepers since their activity on polar products of class II diTPSs is an essential part of plastidial diterpene metabolism, necessary for the departure of non-polar diterpene scaffolds from plastids to the cytoplasm (Mehrshahi et al., 2014), there to be accessed by other metabolic enzymes such as cytochromes P450 and acyltransferases (Bathe and Tissier, 2019; Wang and Peters, 2022; Zhou and Pichersky, 2020). Class I clerodane synthases catalyse the second dedicated step in clerodane biosynthesis by transforming the kolavenyl diphosphate and isokolavenyl diphosphate to kolavenol and isokolavenol, respectively. In Scutellaria species, genes encoding class I clerodane synthases are the result of tandem duplication and neofunctionalization of miltiradiene synthase (Figures 5, 6), whereas phylogenomic data suggest that in *S. splendens* the duplication and neofunctionalization of class I clerodane synthases involved an independent evolutionary event(s). The different evolutionary paths of class I clerodane synthases in Scutellaria and Salvia lineages suggest a polyphyletic origin of clerodane biosynthetic pathway in plastids, with the clerodane pathway in the genus Scutellaria evolving approximately 29.7 MYA while in *S. splendens* it evolved around 14.4 MYA although the observed promiscuity of other class I diTPS in Salvia towards kolavenyl diphosphate (Figures 3, S27) might also support earlier activity time before emergence of *bona-fide* class I clerodane synthases. The evolution of the clerodane pathway in Scutellaria involved gene duplication (Mint Evolutionary GenomicConsortium, 2018; Lichman *et al*., 2020b; Zhao *et al*., 2019a) and neofunctionalization as suggested for other examples of terpene metabolism in the family Lamiaceae. In Salvia, the pathway evolved by a dual mechanism involving adoption of new functionality and increased enzyme promiscuity (Zhao *et al*., 2019b), again supporting polyphyletic origins of clerodane biosynthesis.

There is no evidence that the genes encoding diTPSs or associated enzymes such as cytochromes P450 (CYP) or acyltransferases of clerodane biosynthesis, are organized in a cluster of biosynthetic genes. Although abietane diterpenoids have not been reported in extracts of *S. barbata, S. baicalensis* and *S. splendens*, our work established the common presence of biosynthetic gene cluster (Figure 6) involved in the synthesis of the abietane diterpene, ferruginol. This BCG includes the gene encoding the class I diTPS with miltiradiene synthase activity that gave rise to the class I diTPSs with KLS/IKLS activity in *S. barbata* and *S. baicalensis* illustrating that the BGC can be the source of new genes for diversification in diterpene synthesis. The duplication of class I diTPSs in the two Scutellaria species resulted in enzyme neofunctionalization in clerodane biosynthesis.

Figure 6 shows that biosynthetic gene clusters can serve an important role as “metabolic toolboxes” providing, through gene duplication, new biocatalysts for recruitment into emerging metabolic pathways. However, tandem gene duplications of CYP genes in the ferruginol BGC did not lead to new activities operating in the oxidation of clerodanes in any of these three species although this type of clustering has been reported in several other examples (Polturak et al., 2021; Zhan *et al*., 2022).

We provide a high-quality genome of medicinal plant *S. barbata* accompanied by genome mining and biochemical analysis to map diterpene metabolism in three closely related species in the mint family. Comparing phylogenetically closely related species, results in a much clearer and more focused picture of the microevolution of specialized metabolism, than possible by simple comparisons over large evolutionary time scales. We identified genes encoding class I clerodane synthases from three different species in the Lamiaceae and traced the polyphyletic origins of clerodane biosynthesis in the Lamiaceae and suggests that the widespread occurrence of clerodanes in the mint family is due to convergence by repeated evolution (Cseke et al., 1998; Pichersky and Lewinsohn, 2011). This is reminiscent of other specialized metabolic pathways in plants, such as pyrrolizidine alkaloid defence metabolism (Reimann et al., 2004) and cyanogenic glucoside biosynthesis (Takos et al., 2011). Our results offer pivotal insight for the full elucidation of the biosynthesis of bioactive clerodanes from *S. barbata*, like scutebarbatine A. The polyphyletic origins of clerodanes in Scutellaria and Salvia suggest that the downstream steps in clerodane biosynthesis evolved by convergence which, in turn, may underpin the differences in clerodanes produced by Scutellaria and Salvia genera.

## Accession Numbers

Reference genome data are deposited in GenBank under project number PRJNA649842, and transcriptome sequence reads are deposited in the Sequence Read Archive (SRA) under accession number SRA: SRP275961.

The NCBI accession number of the genes reported in this study can be found in tables S17 and S19.

## Author Contributions

CM and ECT initiated and conceived the project, designed the experimental strategy, and acquired the funding. JY, QZ and AKK advised on the experimental strategy and methodologies. TH, YZ, and XZ performed the genome sequencing. Genome evaluation and annotation, comparative genomic analysis were performed by SW. Gene cloning was performed by HL (*S. barbata, S. baicalensis*), SW (*S. splendens, H. officinalis, S. indicum*), RL (*S. splendens*), YW (*S. splendens*), ALMM (*S. baicalensis*); protein expression in *N. benthamiana* was performed by HL; protein expression in *E. coli* was performed by HL, RL, XY, YW, ALMM; enzyme and GCMS analysis were performed by HL and RL; RL assembled the synthetic biology expression systems for synthesis of miltiradiene and ferruginol in *E. coli* and *S. cerevisiae*; ALMM performed synthesis of GGPP, isolation and characterization of miltiradiene. HL, SW, RL and ECT prepared the figures. HL, SW, CM and ECT wrote the paper, with invaluable input from all the authors.

## Acknowledgments

The authors would like to thank the staff and management of the CEMPS Core Facility Center for the excellent support in metabolomics (GCMS, LCMS, NMR) and microscopy services, as well as the personnel at the CEMPS glasshouse facilities.

Authors gratefully acknowledge Prof A. M. Makris (INAB-CERTH) for providing the AM119 yeast strain; Prof R. Peters (Iowa State University) for providing the pIRS, pGG plasmids; Dr R. Hughes and Dr. M. Franceschetti (JIC) for preparing the modified pTRBO vector; Prof. Yong Wang (CEMPS) for the gift of kaurene.

## References

Andersen, N.R., Lorck, H.O., and Rasmussen, P.R. (1983). Fermentation, isolation and characterization of antibiotic PR-1350. J Antibiot (Tokyo) 36:753–760. 10.7164/antibiotics.36.753.

Andersen-Ranberg, J., Kongstad, K.T., Nielsen, M.T., Jensen, N.B., Pateraki, I., Bach, S.S., Hamberger, B., Zerbe, P., Staerk, D., Bohlmann, J., et al. (2016). Expanding the Landscape of Diterpene Structural Diversity through Stereochemically Controlled Combinatorial Biosynthesis. Angew Chem Int Ed Engl 55:2142–2146. 10.1002/anie.201510650.

Bathe, U., and Tissier, A. (2019). Cytochrome P450 enzymes: A driving force of plant diterpene diversity. Phytochemistry 161:149–162. 10.1016/j.phytochem.2018.12.003.

Berrow, N.S., Alderton, D., Sainsbury, S., Nettleship, J., Assenberg, R., Rahman, N., Stuart, D.I., and Owens, R.J. (2007). A versatile ligation-independent cloning method suitable for high-throughput expression screening applications. Nucleic Acids Res 35:e45. 10.1093/nar/gkm047.

Božić, D., Papaefthimiou, D., Brückner, K., de Vos, R.C., Tsoleridis, C.A., Katsarou, D., Papanikolaou, A., Pateraki, I., Chatzopoulou, F.M., Dimitriadou, E., et al. (2015). Towards Elucidating Carnosic Acid Biosynthesis in Lamiaceae: Functional Characterization of the Three First Steps of the Pathway in Salvia fruticosa and Rosmarinus officinalis. PLoS One 10:e0124106. 10.1371/journal.pone.0124106.

Chen, F., Tholl, D., Bohlmann, J., and Pichersky, E. (2011). The family of terpene synthases in plants: a mid-size family of genes for specialized metabolism that is highly diversified throughout the kingdom. Plant J 66:212–229. 10.1111/j.1365-313X.2011.04520.x.

Chen, Q., Rahman, K., Wang, S.J., Zhou, S., and Zhang, H. (2020). Scutellaria barbata: A Review on Chemical Constituents, Pharmacological Activities and Clinical Applications. Curr Pharm Des 26:160–175. 10.2174/1381612825666191216124310.

Chen, X., Berim, A., Dayan, F.E., and Gang, D.R. (2017). A (-)-kolavenyl diphosphate synthase catalyzes the first step of salvinorin A biosynthesis in Salvia divinorum. J Exp Bot 68:1109–1122. 10.1093/jxb/erw493.

Christianson, D.W. (2017). Structural and Chemical Biology of Terpenoid Cyclases. Chem Rev 117:11570–11648. 10.1021/acs.chemrev.7b00287.

Consortium, M.E.G. (2018). Phylogenomic Mining of the Mints Reveals Multiple Mechanisms Contributing to the Evolution of Chemical Diversity in Lamiaceae. Mol Plant 11:1084–1096. 10.1016/j.molp.2018.06.002.

Cseke, L., Dudareva, N., and Pichersky, E. (1998). Structure and evolution of linalool synthase. Mol Biol Evol 15:1491–1498. 10.1093/oxfordjournals.molbev.a025876.

Cui, G., Duan, L., Jin, B., Qian, J., Xue, Z., Shen, G., Snyder, J.H., Song, J., Chen, S., Huang, L., et al. (2015). Functional Divergence of Diterpene Syntheses in the Medicinal Plant Salvia miltiorrhiza. Plant Physiol 169:1607–1618. 10.1104/pp.15.00695.

Dai, S.J., Tao, J.Y., Liu, K., Jiang, Y.T., and Shen, L. (2006). neo-Clerodane diterpenoids from Scutellaria barbata with cytotoxic activities. Phytochemistry 67:1326–1330. 10.1016/j.phytochem.2006.04.024.

De La Peña, R., and Sattely, E.S. (2021). Rerouting plant terpene biosynthesis enables momilactone pathway elucidation. Nat Chem Biol 17:205–212. 10.1038/s41589-020-00669-3.

Dong, A.X., Xin, H.B., Li, Z.J., Liu, H., Sun, Y.Q., Nie, S., Zhao, Z.N., Cui, R.F., Zhang, R.G., Yun, Q.Z., et al. (2018). High-quality assembly of the reference genome for scarlet sage, Salvia splendens, an economically important ornamental plant. Gigascience 710.1093/gigascience/giy068.

Emanuelsson, O., Nielsen, H., and von Heijne, G. (1999). ChloroP, a neural network-based method for predicting chloroplast transit peptides and their cleavage sites. Protein Sci 8:978–984. 10.1110/ps.8.5.978.

Falara, V., Akhtar, T.A., Nguyen, T.T., Spyropoulou, E.A., Bleeker, P.M., Schauvinhold, I., Matsuba, Y., Bonini, M.E., Schilmiller, A.L., Last, R.L., et al. (2011). The tomato terpene synthase gene family. Plant Physiol 157:770–789. 10.1104/pp.111.179648.

Feng, P.P., Qi, Y.K., Li, N., and Fei, H.R. (2021). Scutebarbatine A induces cytotoxicity in hepatocellular carcinoma via activation of the MAPK and ER stress signaling pathways. J Biochem Mol Toxicol:e22731. 10.1002/jbt.22731.

Fontana, G., Savona, G., and Rodríguez, B. (2006). Complete 1H and 13C NMR assignments of clerodane diterpenoids of Salvia splendens. Magn Reson Chem 44:962–965. 10.1002/mrc.1869.

Franke, J., Kim, J., Hamilton, J.P., Zhao, D., Pham, G.M., Wiegert-Rininger, K., Crisovan, E., Newton, L., Vaillancourt, B., Tatsis, E., et al. (2019). Gene Discovery in Gelsemium Highlights Conserved Gene Clusters in Monoterpene Indole Alkaloid Biosynthesis. Chembiochem 20:83–87. 10.1002/cbic.201800592.

Frezza, C., Venditti, A., Serafini, M., and Bianco, A. (2019). Phytochemistry, chemotaxonomy, ethnopharmacology, and nutraceutics of Lamiaceae. Studies in natural products chemistry 62:125–178.

Godden, G.T., Kinser, T.J., Soltis, P.S., and Soltis, D.E. (2019). Phylotranscriptomic Analyses Reveal Asymmetrical Gene Duplication Dynamics and Signatures of Ancient Polyploidy in Mints. Genome Biol Evol 11:3393–3408. 10.1093/gbe/evz239.

Hamilton, J.P., Godden, G.T., Lanier, E., Bhat, W.W., Kinser, T.J., Vaillancourt, B., Wang, H., Wood, J.C., Jiang, J., Soltis, P.S., et al. (2020). Generation of a chromosome-scale genome assembly of the insect-repellent terpenoid-producing Lamiaceae species, Callicarpa americana. Gigascience 910.1093/gigascience/giaa093.

He, J., Xin, P., Ma, X., Chu, J., and Wang, G. (2020). Gibberellin Metabolism in Flowering Plants: An Update and Perspectives. Frontiers in Plant Science 1110.3389/fpls.2020.00532.

Heskes, A.M., Sundram, T.C.M., Boughton, B.A., Jensen, N.B., Hansen, N.L., Crocoll, C., Cozzi, F., Rasmussen, S., Hamberger, B., Hamberger, B., et al. (2018). Biosynthesis of bioactive diterpenoids in the medicinal plant Vitex agnus-castus. Plant J 93:943–958. 10.1111/tpj.13822.

Hill, S.J., Brion, A., and Shenvi, R.A. (2020). Chemical syntheses of the salvinorin chemotype of KOR agonist. Nat Prod Rep 37:1478–1496. 10.1039/d0np00028k.

Hussein, A.A., de la Torre, M.C., Jimeno, M.-L., Rodríguez, B., Bruno, M., Piozzi, F., and Servettaz, O. (1996). A neo-clerodane diterpenoid from Scutellaria baicalensis. Phytochemistry 43:835–837. https://doi.org/10.1016/0031-9422(96)00380-9.

Jia, M., Potter, K.C., and Peters, R.J. (2016). Extreme promiscuity of a bacterial and a plant diterpene synthase enables combinatorial biosynthesis. Metab Eng 37:24–34. 10.1016/j.ymben.2016.04.001.

Jia, M., Mishra, S.K., Tufts, S., Jernigan, R.L., and Peters, R.J. (2019). Combinatorial biosynthesis and the basis for substrate promiscuity in class I diterpene synthases. Metab Eng 55:44–58. 10.1016/j.ymben.2019.06.008.

Johnson, S.R., Bhat, W.W., Bibik, J., Turmo, A., Hamberger, B., and Hamberger, B. (2019). A database-driven approach identifies additional diterpene synthase activities in the mint family (Lamiaceae). J Biol Chem 294:1349–1362. 10.1074/jbc.RA118.006025.

Karunanithi, P.S., and Zerbe, P. (2019). Terpene Synthases as Metabolic Gatekeepers in the Evolution of Plant Terpenoid Chemical Diversity. Front Plant Sci 10:1166. 10.3389/fpls.2019.01166.

Kutrzeba, L., Dayan, F.E., Howell, J., Feng, J., Giner, J.L., and Zjawiony, J.K. (2007). Biosynthesis of salvinorin A proceeds via the deoxyxylulose phosphate pathway. Phytochemistry 68:1872–1881. 10.1016/j.phytochem.2007.04.034.

Kwon, M., Utomo, J.C., Park, K., Pascoe, C.A., Chiorean, S., Ngo, I., Pelot, K.A., Pan, C.-H., Kim, S.-W., Zerbe, P., et al. (2022). Cytochrome P450-Catalyzed Biosynthesis of a Dihydrofuran Neoclerodane in Magic Mint (Salvia divinorum). ACS Catalysis 12:777–782. 10.1021/acscatal.1c03691.

Lemke, C., Potter, K.C., Schulte, S., and Peters, R.J. (2019). Conserved bases for the initial cyclase in gibberellin biosynthesis: from bacteria to plants. Biochem J 476:2607–2621. 10.1042/bcj20190479.

Lhullier, C., de Oliveira Tabalipa, E., Nienkötter Sardá, F., Sandjo, L.P., Zanchett Schneider, N.F., Carraro, J.L., Oliveira Simões, C.M., and Schenkel, E.P. (2019). Clerodane Diterpenes from the Marine Sponge Raspailia bouryesnaultae Collected in South Brazil. Mar Drugs 1710.3390/md17010057.

Li, M., Zhang, D., Gao, Q., Luo, Y., Zhang, H., Ma, B., Chen, C., Whibley, A., Zhang, Y., Cao, Y., et al. (2019). Genome structure and evolution of Antirrhinum majus L. Nat Plants 5:174–183. 10.1038/s41477-018-0349-9.

Li, R., Morris-Natschke, S.L., and Lee, K.H. (2016). Clerodane diterpenes: sources, structures, and biological activities. Nat Prod Rep 33:1166–1226. 10.1039/c5np00137d.

Li, Y., Leveau, A., Zhao, Q., Feng, Q., Lu, H., Miao, J., Xue, Z., Martin, A.C., Wegel, E., Wang, J., et al. (2021). Subtelomeric assembly of a multi-gene pathway for antimicrobial defense compounds in cereals. Nat Commun 12:2563. 10.1038/s41467-021-22920-8.

Lichman, B.R., Godden, G.T., and Buell, C.R. (2020a). Gene and genome duplications in the evolution of chemodiversity: perspectives from studies of Lamiaceae. Curr Opin Plant Biol 55:74–83. 10.1016/j.pbi.2020.03.005.

Lichman, B.R., Godden, G.T., Hamilton, J.P., Palmer, L., Kamileen, M.O., Zhao, D., Vaillancourt, B., Wood, J.C., Sun, M., Kinser, T.J., et al. (2020b). The evolutionary origins of the cat attractant nepetalactone in catnip. Sci Adv 6:eaba0721. 10.1126/sciadv.aba0721.

Lindbo, J.A. (2007). TRBO: a high-efficiency tobacco mosaic virus RNA-based overexpression vector. Plant Physiol 145:1232–1240. 10.1104/pp.107.106377.

Macías, F.A., Lacret, R., Varela, R.M., Nogueiras, C., and Molinillo, J.M. (2010). Isolation and phytotoxicity of terpenes from Tectona grandis. J Chem Ecol 36:396–404. 10.1007/s10886-010-9769-3.

Marconett, C.N., Morgenstern, T.J., San Roman, A.K., Sundar, S.N., Singhal, A.K., and Firestone, G.L. (2010). BZL101, a phytochemical extract from the Scutellaria barbata plant, disrupts proliferation of human breast and prostate cancer cells through distinct mechanisms dependent on the cancer cell phenotype. Cancer Biol Ther 10:397–405. 10.4161/cbt.10.4.12424.

Mehrshahi, P., Johnny, C., and DellaPenna, D. (2014). Redefining the metabolic continuity of chloroplasts and ER. Trends Plant Sci 19:501–507. 10.1016/j.tplants.2014.02.013.

Merritt, A.T., and Ley, S.V. (1992). Clerodane diterpenoids. Nat Prod Rep 9:243–287. 10.1039/np9920900243.

Mistry, J., Chuguransky, S., Williams, L., Qureshi, M., Salazar, Gustavo A., Sonnhammer, E.L.L., Tosatto, S.C.E., Paladin, L., Raj, S., Richardson, L.J., et al. (2020). Pfam: The protein families database in 2021. Nucleic Acids Research 49:D412–D419. 10.1093/nar/gkaa913.

Muchlinski, A., Jia, M., Tiedge, K., Fell, J.S., Pelot, K.A., Chew, L., Davisson, D., Chen, Y., Siegel, J., Lovell, J.T., et al. (2021). Cytochrome P450-catalyzed biosynthesis of furanoditerpenoids in the bioenergy crop switchgrass (Panicum virgatum L.). Plant J 108:1053–1068. 10.1111/tpj.15492.

Nakano, C., Oshima, M., Kurashima, N., and Hoshino, T. (2015). Identification of a New Diterpene Biosynthetic Gene Cluster that Produces O-Methylkolavelool in Herpetosiphon aurantiacus. ChemBioChem 16:772–781. https://doi.org/10.1002/cbic.201402652.

Ostrozhenkova, E. (2008). Metabolite and isotopologue profiling in plants. Studies on the biosynthesis of terpenoids and alkaloids.. Lehrstuhl für Biochemie der Technischen Universität München Technische Universität München.

Pelot, K.A., Hagelthorn, D.M., Hong, Y.J., Tantillo, D.J., and Zerbe, P. (2019). Diterpene Synthase-Catalyzed Biosynthesis of Distinct Clerodane Stereoisomers. Chembiochem 20:111–117. 10.1002/cbic.201800580.

Pelot, K.A., Mitchell, R., Kwon, M., Hagelthorn, L.M., Wardman, J.F., Chiang, A., Bohlmann, J., Ro, D.K., and Zerbe, P. (2017). Biosynthesis of the psychotropic plant diterpene salvinorin A: Discovery and characterization of the Salvia divinorum clerodienyl diphosphate synthase. Plant J 89:885–897. 10.1111/tpj.13427.

Perez, A.T., Arun, B., Tripathy, D., Tagliaferri, M.A., Shaw, H.S., Kimmick, G.G., Cohen, I., Shtivelman, E., Caygill, K.A., Grady, D., et al. (2010). A phase 1B dose escalation trial of Scutellaria barbata (BZL101) for patients with metastatic breast cancer. Breast Cancer Res Treat 120:111–118. 10.1007/s10549-009-0678-5.

Pichersky, E., and Lewinsohn, E. (2011). Convergent evolution in plant specialized metabolism. Annu Rev Plant Biol 62:549–566. 10.1146/annurev-arplant-042110-103814.

Polturak, G., Liu, Z., and Osbourn, A. (2022). New and emerging concepts in the evolution and function of plant biosynthetic gene clusters. Current Opinion in Green and Sustainable Chemistry 33:100568. https://doi.org/10.1016/j.cogsc.2021.100568.

Polturak, G., Dippe, M., Stephenson, M.J., Misra, R.C., Owen, C., Ramirez-Gonzalez, R., Haidioulis, J., Schoonbeek, H.-J., Chartrain, L., Borrill, P., et al. (2021). Genome Mining Uncovers Clustered Biosynthetic Pathways for Defense-Related Molecules in Bread Wheat. bioRxiv:doi:10.1101/2021.1111.1104.467362.10.1101/2021.11.04.467362.

Potter, K.C., Zi, J., Hong, Y.J., Schulte, S., Malchow, B., Tantillo, D.J., and Peters, R.J. (2016). Blocking Deprotonation with Retention of Aromaticity in a Plant ent-Copalyl Diphosphate Synthase Leads to Product Rearrangement. Angew Chem Int Ed Engl 55:634–638. 10.1002/anie.201509060.

Reimann, A., Nurhayati, N., Backenköhler, A., and Ober, D. (2004). Repeated evolution of the pyrrolizidine alkaloid-mediated defense system in separate angiosperm lineages. Plant Cell 16:2772–2784. 10.1105/tpc.104.023176.

Roncero, A.M., Tobal, I.E., Moro, R.F., Díez, D., and Marcos, I.S. (2018). Halimane diterpenoids: sources, structures, nomenclature and biological activities. Nat Prod Rep 35:955–991. 10.1039/c8np00016f.

Shen, J., Li, P., Liu, S., Liu, Q., Li, Y., Sun, Y., He, C., and Xiao, P. (2021). Traditional uses, ten-years research progress on phytochemistry and pharmacology, and clinical studies of the genus Scutellaria. J Ethnopharmacol 265:113198. 10.1016/j.jep.2020.113198.

Smit, S.J., and Lichman, B.R. (2022). Plant biosynthetic gene clusters in the context of metabolic evolution. Nat Prod Rep 39:1465–1482. 10.1039/d2np00005a.

Su, P., Tong, Y., Cheng, Q., Hu, Y., Zhang, M., Yang, J., Teng, Z., Gao, W., and Huang, L. (2016). Functional characterization of ent-copalyl diphosphate synthase, kaurene synthase and kaurene oxidase in the Salvia miltiorrhiza gibberellin biosynthetic pathway. Sci Rep 6:23057. 10.1038/srep23057.

Sun, W., Leng, L., Yin, Q., Xu, M., Huang, M., Xu, Z., Zhang, Y., Yao, H., Wang, C., Xiong, C., et al. (2019). The genome of the medicinal plant Andrographis paniculata provides insight into the biosynthesis of the bioactive diterpenoid neoandrographolide. Plant J 97:841–857. 10.1111/tpj.14162.

Takos, A.M., Knudsen, C., Lai, D., Kannangara, R., Mikkelsen, L., Motawia, M.S., Olsen, C.E., Sato, S., Tabata, S., Jørgensen, K., et al. (2011). Genomic clustering of cyanogenic glucoside biosynthetic genes aids their identification in Lotus japonicus and suggests the repeated evolution of this chemical defence pathway. Plant J 68:273–286. 10.1111/j.1365-313X.2011.04685.x.

Tomlinson, M., Man, Z., Elaine, J.B., Jie, L., Haixiu, L., Juri, F., Lionel, H., Gerhard, S., Martin, R., Dongfeng, Y., et al. (2022). Diterpenoids from Scutellaria barbata induce tumour-selective cytotoxicity by taking the brakes off apoptosis. Medicinal Plant Biology 1:1–16. 10.48130/MPB-2022-0003.

Tudzynski, B. (2005). Gibberellin biosynthesis in fungi: genes, enzymes, evolution, and impact on biotechnology. Appl Microbiol Biotechnol 66:597–611. 10.1007/s00253-004-1805-1.

Vestri Alvarenga, S.A., Pierre Gastmans, J., do Vale Rodrigues, G., Moreno, P.R., and de Paulo Emerenciano, V. (2001). A computer-assisted approach for chemotaxonomic studies--diterpenes in Lamiaceae. Phytochemistry 56:583–595. 10.1016/s0031-9422(00)00424-6.

Wang, L., Chen, W., Li, M., Zhang, F., Chen, K., and Chen, W. (2020a). A review of the ethnopharmacology, phytochemistry, pharmacology, and quality control of Scutellaria barbata D. Don. J Ethnopharmacol 254:112260. 10.1016/j.jep.2019.112260.

Wang, M., Ma, C., Chen, Y., Li, X., and Chen, J. (2019). Cytotoxic Neo-Clerodane Diterpenoids from Scutellaria barbata D.Don. Chem Biodivers 16:e1800499. 10.1002/cbdv.201800499.

Wang, M., Chen, Y., Hu, P., Ji, J., Li, X., and Chen, J. (2020b). Neoclerodane diterpenoids from Scutellaria barbata with cytotoxic activities. Nat Prod Res 34:1345–1351. 10.1080/14786419.2018.1514399.

Wang, Z., and Peters, R.J. (2022). Tanshinones: Leading the way into Lamiaceae labdane-related diterpenoid biosynthesis. Curr Opin Plant Biol 66:102189. 10.1016/j.pbi.2022.102189.

Wu, S., Malaco Morotti, A.L., Wang, S., Wang, Y., Xu, X., Chen, J., Wang, G., and Tatsis, E.C. (2022). Convergent gene clusters underpin hyperforin biosynthesis in St John’s wort. New Phytologist 235:646–661. https://doi.org/10.1111/nph.18138.

Xu, H., Song, J., Luo, H., Zhang, Y., Li, Q., Zhu, Y., Xu, J., Li, Y., Song, C., Wang, B., et al. (2016). Analysis of the Genome Sequence of the Medicinal Plant Salvia miltiorrhiza. Mol Plant 9:949–952. 10.1016/j.molp.2016.03.010.

Yang, G.C., Hu, J.H., Li, B.L., Liu, H., Wang, J.Y., and Sun, L.X. (2018). Six New neo-Clerodane Diterpenoids from Aerial Parts of Scutellaria barbata and Their Cytotoxic Activities. Planta Med 84:1292–1299. 10.1055/a-0638-8255.

Yang, Z., and Nielsen, R. (2008). Mutation-selection models of codon substitution and their use to estimate selective strengths on codon usage. Mol Biol Evol 25:568–579. 10.1093/molbev/msm284.

Yuan, Q.Q., Song, W.B., Wang, W.Q., and Xuan, L.J. (2017). Scubatines A-F, new cytotoxic neo-clerodane diterpenoids from Scutellaria barbata D. Don. Fitoterapia 119:40–44. 10.1016/j.fitote.2017.03.012.

Zerbe, P., and Bohlmann, J. (2015). Plant diterpene synthases: exploring modularity and metabolic diversity for bioengineering. Trends in Biotechnology 33:419–428. https://doi.org/10.1016/j.tibtech.2015.04.006.

Zhan, C., Shen, S., Yang, C., Liu, Z., Fernie, A.R., Graham, I.A., and Luo, J. (2022). Plant metabolic gene clusters in the multi-omics era. Trends in Plant Science https://doi.org/10.1016/j.tplants.2022.03.002.

Zhao, D., Hamilton, J.P., Bhat, W.W., Johnson, S.R., Godden, G.T., Kinser, T.J., Boachon, B., Dudareva, N., Soltis, D.E., Soltis, P.S., et al. (2019a). A chromosomal-scale genome assembly of Tectona grandis reveals the importance of tandem gene duplication and enables discovery of genes in natural product biosynthetic pathways. Gigascience 810.1093/gigascience/giz005.

Zhao, Q., Chen, X.Y., and Martin, C. (2016). Scutellaria baicalensis, the golden herb from the garden of Chinese medicinal plants. Sci Bull (Beijing) 61:1391–1398. 10.1007/s11434-016-1136-5.

Zhao, Q., Yang, J., Cui, M.Y., Liu, J., Fang, Y., Yan, M., Qiu, W., Shang, H., Xu, Z., Yidiresi, R., et al. (2019b). The Reference Genome Sequence of Scutellaria baicalensis Provides Insights into the Evolution of Wogonin Biosynthesis. Mol Plant 12:935–950. 10.1016/j.molp.2019.04.002.

Zhou, F., and Pichersky, E. (2020). More is better: the diversity of terpene metabolism in plants. Current Opinion in Plant Biology 55:1–10. https://doi.org/10.1016/j.pbi.2020.01.005.

