## Supplementary material for "The genomes of medicinal skullcaps reveal the polyphyletic origins of clerodane diterpene biosynthesis in the family Lamiaceae": SI material

†These three authors contribute equally to present work.

**This PDF file includes:**

Materials and Methods
Figs. S1 to S35
Tables S1 to S20
Data S1 to S2
References

### **Materials and Methods**

#### Plant material

Plants of *Scutellaria barbata*, *Scutellaria baicalensis* and *Salvia splendens* were acquired from Shanghai Chenshan Botanical Garden and kept in greenhouse conditions.

#### Genome sequencing

##### ***Illumina paired-end short read sequencing***

Genomic DNA was extracted from leaves of an *S. barbata* plant using DNAsecure Plant Kit (TIANGEN, China). DNA degradation and contamination were monitored on 1% agarose gels. DNA purity was validated using a NanoPhotometer® spectrophotometer (IMPLEN, CA, USA). DNA concentration was measured using Qubit® DNA Assay Kit in Qubit® 2.0 Fluorometer (Life Technologies, CA, USA).

A total amount of 700ng DNA per sample was used as input material for the DNA sample preparations. Sequencing libraries were generated using NEB Next® Ultra DNA Library Prep Kit for Illumina® (NEB, USA) following the manufacturer's instructions and sequenced on an Illumina NovaSeq platform to generate 150bp paired-end reads.

##### ***Pacbio long-read sequencing***

For long-read sequencing, genomic DNA was extracted from the same plant using a modified Phenol-Chloroform method. Libraries for Single Molecule, Real-Time (SMRT) Sequencing were constructed following the standard instructions of Pacific Biosciences (www.pacb.com). The library was constructed by shearing whole genomic DNA to ~20 kb targeted size, followed by damage repair, ends repair, blunt-end ligation and size selection. The large insert SMRTbell library was sequenced on the Pacbio Sequel platform.

##### ***Bionano genome imaging***

The high-molecular-weight genomic DNA was isolated by BioNanoPrep Plant Tissue DNA Isolation Kit (Bionano Genomics, USA). Using the NLRs DNA Labeling Kit (Bionano, USA), a single enzymatic reaction (DLE-1) labels specific sequences across the entire genome. The long, labeled DNA molecules were linearized in nanochannel arrays on a Saphyr chip® and imaged by the Saphyr® Instrument.

##### ***Hi-C library sequencing***

Following the standard protocol described previously with modifications (Belton et al., 2012), genomic DNA from the leaves of *S. barbata* was cross-linked using 4% formaldehyde solution in a vacuum. After quenching the crosslinking reaction, an overnight digestion was applied to the samples with the restriction enzyme MboI (?). The ends of fragments were marked with biotin and the proximal chromatin DNA was re-ligated. DNA was purified by the phenol-chloroform method and biotin was removed from non-ligated fragment ends. After adding A-tails to the fragment ends and following ligation by the illumina paired-end (PE) sequencing adapters, Hi-C sequencing libraries were amplified by PCR and sequenced on Illumina NovaSeq platform.

#### Genome survey

Illumina short reads were used to evaluate the genome size and heterozygosity rate of *S. barbata* genome by *k-mer* analysis (Fig. S1). *K-mers* were counted by Jellyfish (www.genome.umd.edu/jellyfish.html) with the *k-mer* size of 31. GenomeScope (github.com/schatzlab/genomescope) uses the *k-mer* count distribution to infer the global properties of *S. barbata* genome.

#### Genome assembly

*De novo* assembly of the Pacbio long reads was conducted by FALCON v0.3.0 (github.com/PacificBiosciences/falcon). All overlaps in the raw reads were identified and the overlap information was used to error-correct the reads. Overlap detection between corrected reads generated the contig assembly. Then, the contig sequences were polished by Quiver (SMRT® Analysis suite) to produce the primary assembly. Bionano raw data were *de novo* assembled into consensus physical maps using IrysView v. Hybrid scaffolds (first version reference genome assembly) were constructed by integrating *de novo* primary assembly and Bionano genome imaging data using Bionano Solve v. The Hi-C sequencing data were mapped to the assembled scaffolds to get the order and orientation information of scaffolds using BWA v (github.com/lh3/bwa). The scaffolds were anchored to pseudochromosomes by LACHESIS v (github.com/shendurelab/LACHESIS).

#### Transcriptomic analysis

##### ***RNA sequencing***

For the global tissue-specific transcriptome analyses, young leaves, old leaves, young stems, old stems, flowers and roots were collected from *S. barbata* plants with three biological replicates.

For the trichome-specific transcriptomes, leaves, leaves without trichomes and trichomes were isolated as described previously (Tomlinson et al., 2022).

Total RNA was extracted using a modified cetyltrimethylammonium bromide (CTAB) method for each sample (Meisel et al., 2005). RNA purity was checked with OD260/OD280 value using NanoDrop (Thermo Fisher Scientific, USA). RNA degradation and potential contamination were tested by agarose gel electrophoresis. 2100 Bioanalyzer (Agilent Technologies, USA) was used to evaluate the integrity of RNA by measuring the insert size. After quality control, mRNA was enriched using oligo(dT) beads. The mRNA was then fragmented randomly, followed by cDNA synthesis using random hexamers and reverse transcriptase. Following first-strand synthesis, a custom second-strand synthesis buffer (Illumina, USA) was added with dNTPs, RNase H and *Escherichia coli* polymerase I to generate the second strand by nick-translation.

After purification, terminal repair, oligo(dA)-tailing, ligation of sequencing adapters, size selection and PCR enrichment, the final cDNA libraries were sequenced on the Illumina HiSeq X platform to generate 150 bp pair-end reads.

##### ***Reference genome-based reads mapping***

To obtain clean reads with high quality, raw reads were pruned using Trimmomatic v0.36 with following parameters: LEADING:3 TRAILING:3 SLIDINGWINDOW:4:15 MINLEN:50. The clean data were mapped to the draft genome by HISAT2 v2.0.4 (daehwankimlab.github.io/hisat2) using default parameters. Fragments Per Kilobase of transcript per Million mapped reads (FPKM) values were generated by custom scripts using read counts produced by HTSeq v0.6.1. SNPs and Indels were detected using GATK v3.5.

##### ***De Novo Transcriptome Assembly***

The clean reads were used as input for Trinity v2.4.0 to complete the *de novo* transcriptome reconstruction process. The primary contigs were clustered and filtered to yield a set of non-redundant representative sequences using CD-HIT v4.6 with default parameters.

##### **Genome annotation**

###### ***Annotation of repetitive DNA sequence***

A combined strategy based on homology alignment and *ab initio* prediction was applied to search the repeat sequences. The homology alignment method used Repbase (www.girinst.org/repbase) as the library to extract potential repeat regions in the *S. barbata* genome by applying RepeatMasker v4.1.0 (http://www.repeatmasker.org/) using default parameters. For *ab initio* prediction method, tandem repeat sequences were extracted using TRF v4.09 (tandem.bu.edu/trf/trf.html) while LTR\_FINDER (tlife.fudan.edu.cn/tlife/ltr_finder/), RepeatScout v1.0.5 (www.repeatmasker.org/) and RepeatModeler v2.0.1 (www.repeatmasker.org/RepeatModeler) provided the raw transposable element(TE) library. The combination of all libraries above was processed by uclust (www.drive5.com/usearch) to generate a non-redundant pool and supplied to RepeatMasker v4.1.0 for the identification of repetitive sequences across the genome.

#### ***Annotation of non-coding RNA***

In order to search for tRNA genes in the *S. barbata* genome, tRNAscan-SE (lowelab.ucsc.edu/tRNAscan-SE/) was used for prediction by capturing the primary sequence and secondary structure information from tRNA training data. For highly conserved rRNAs, rRNAs from relatively close species were used to search the homologous sequences by BLAST. Other ncRNAs, including miRNAs, snRNAs were identified by searching against the Rfam (rfam.xfam.org) database using infernal (http://infernal.janelia.org/) with default parameters.

#### ***Annotation of protein-coding gene***

Structural annotation of the protein-coding genes for the *S. barbata* genome incorporated *ab initio* prediction, homology-based prediction and RNA-Seq assisted prediction. Augustus v3.2.3, Geneid v1.4, Genescan v1.0, GlimmerHMM v3.04 (ccb.jhu.edu/software/glimmerhmm) and SNAP v2013.11.29 were used in *ab initio* prediction. Sequences of protein-coding genes from *S. baicalensis*, *S. splendens*, *S. miltiorrhiza*, *S. indicum* genome were mapped to the *S. barbata* genome using tblastn v2.2.26 with E-value $\leq 1e^{-5}$, and then supplied to GeneWise v2.4.1 to predict gene structure. RNA-seq mapping results from different samples were used to identify splice sites by Stringtie v1.3.3. The predictions from these three approaches were merged to a non-redundant reference gene set with EvidenceModeler v1.1.1 (EVM, evidencemodeler.sourceforge.net/). The annotation produced by EVM was further curated using splice site information by PASA (github.com/PASApipeline/PASApipeline).

Functional annotation for the gene set was obtained by searching against publicly available databases, including SwissProt (http://www.uniprot.org), Nr (www.ncbi.nlm.nih.gov/protein), Pfam (pfam.xfam.org), KEGG (http://www.genome.jp/kegg) InterPro (www.ebi.ac.uk/interpro). The best match from each database was assigned to the protein-coding gene.

For the structural annotation of the genes from *U. gibba*, *A. paniculata*, *P. citriodora* genome, publicly available RNA-seq data from NCBI (SRR5046448, SRR3722534, SRR5825981) were aligned to the corresponding genome assembly (PRJNA383049, PRJNA421867, PRJNA339064). The mapping results then assisted *ab initio* prediction by GeneMark-ES v4.61\_lic (exon.gatech.edu/GeneMark/).

##### Evaluation of genome assembly and annotation

###### ***Assessment of genome assembly and annotation completeness***

Benchmarking Universal Single-Copy Orthologs (BUSCO) v4.0.6 (github.com/ezlab/busco) was used to estimate the quality of *S. barbata* genome, while Embryophyta odb10 was chosen as the conserved gene set and tomato as the training model.

###### ***Assessment of genome assembly accuracy***

To evaluate the accuracy of the genome assembly, short reads were mapped to the assembly using BWA (bio-bwa.sourceforge.net/).

##### Comparative Genomic Analysis

###### ***Phylogenetic tree of species***

Orthologue clustering was inferred by OrthoFinder v2.3.11 (github.com/davidemms/OrthoFinder) among the genome of *S. barbata*, *S. baicalensis* (PRJNA484052), *S. splendens* (gigadb.org/dataset/100463), *Salvia miltiorrhiza* (gigadb.org/dataset/100164, ftp://danshen.ndctcm.org:10402/), *Tectona grandis* (gigadb.org/dataset/100550), *Hyssopus officinalis* (datadryad.org/stash/dataset/doi:10.5061/dryad.88tj450), *Nepeta cataria* (datadryad.org/stash/dataset/doi:10.5061/dryad.88tj450), *Nepeta mussinii* (datadryad.org/stash/dataset/doi:10.5061/dryad.88tj450), *Perilla citriodora* (PRJNA339064), *Sesamum indicum* (PRJNA186669), *Andrographis paniculata* (PRJNA421867), *Antirrhinum majus* (http://bioinfo.sibs.ac.cn/Am), *Utricularia gibba* (PRJNA383049, PRJNA207602), *Solanum lycopersicum* (SL3.0, EnsemblPlants), *Coffea canephora* (AUK\_PRJEB4211\_v1, EnsemblPlants), *Arabidopsis thaliana* (TAIR10, EnsemblPlants), *Glycine max* (B73\_RefGen\_v4, EnsemblPlants), *Populus trichocarpa* (Pop\_tri\_v3, EnsemblPlants), *Vitis vinifera* (12X, EnsemblPlants), *Oryza sativa Japonica* (IRGSP-1.0, EnsemblPlants), *Nymphaea colorata* (ASM883128v1, EnsemblPlants), *Amborella trichopoda* (AMTR1.0, EnsemblPlants). The protein sequences from each genome were then filtered by keeping the longest isoform and discarding sequences less than 50 amino acids. The polished sets were then fed to OrthoFinder with the following parameters: multiple sequence alignment MAFFT, sequence search BLASTP, tree inference FastTree.

#### ***Species divergence times***

MCMCTree from PAML v4.9j (abacus.gene.ucl.ac.uk/software/paml.html) suite was used to estimate the divergence time with the following parameters: burnin 5000000, sampfreq 50, nsample 1000000. For the general phylogram of species (Fig. S6), the restriction of certain speciation was based on calibration points described previously (Zhao et al., 2019). The divergence time generated by the general phylogram was further used to calibrate the phylogenetic tree focusing on Lamiaceae species (Fig. S6).

#### ***Gene family evolution***

Changes in gene family size were analyzed by CAFE v4.2.1 (github.com/hahnlab/CAFE) to calculate the expansions and contractions of gene families for each species.

The shared and unique gene families between *S. barbata*, *A. majus*, *C. canephora*, *A. thaliana* and *O. sativa* genome were calculated and plotted using the VennDiagram package in R, while the upset diagram was drawn using the modified UpSetR package (github.com/GuangchuangYu/UpSetR).

Genes from expanded gene families in *S. barbata* were extracted to carry out the Gene Ontology (GO) annotation using InterProScan v5.44-79.0. The results were further supplied to clusterProfiler (guangchuangyu.github.io/software/clusterProfiler/) for enrichment analysis. For Over Representation Analysis (ORA) method, the cutoff of *P* value was set to 0.05.

#### ***Synteny analysis***

Genomes and total protein sequences of species used in phylogenetic analysis were loaded to MCscan (Python version), a package from JCVI utility libraries (github.com/tanghaibao/jcvi). Comparisons between gene pairs were performed using LAST (last.cbrc.jp). After the removal of potential tandem duplications and hits with low scores, the anchors from LAST outputs were clustered into syntenic blocks. The dot plots, macrosynteny plots and microsynteny plots were generated by dotplot, karyotype, synteny functions with default parameters respectively.

#### ***Speciation and whole genome duplication (WGD) analysis***

Homologous gene pairs were obtained by BLASTP using the proteomes of *S. barbata*, *S. baicalensis*, *S. splendens*, *T. grandis*, *A. paniculata*. Syntenic genes within each genome and between genomes were inferred by MCScanX based on the combined information of gene similarity and gene order. The synonymous substitution rate (Ks) of gene pairs was calculated using CodeML function in PAML.

#### ***Identification of diterpene synthase genes***

A joint methodology including Pfam searching, homologous alignment and syntenic analysis was applied to discover diterpene synthase genes across the genomes of *S. barbata*, *S. baicalensis* and *S. splendens*. Two Pfam domains, PF01397 (terpene synthase, N-terminal domain) and PF03936 (terpene synthase family, metal binding domain), were used to search against the whole genome by HMMER v3.3 (hmmer.org). The characterized class I and class II diterpene synthase genes in Lamiaceae plants which were retrieved from NCBI were merged into the custom library separately. BLAST with e-value = $1 \times 10^{-5}$ was performed to explore homologues against the libraries. The gene pairs obtained from synteny analysis were further analyzed by selecting pairs containing known diterpene synthase genes.

#### ***Phylogeny of class I and class II diterpene synthases***

The diterpene synthase genes identified by the combined strategy from *S. barbata*, *S. baicalensis* and *S. splendens*, accompanied by the genes with verified activities, were used to construct the phylogenetic tree by RAxML-NG v0.9.0 (github.com/amkozlov/raxml-ng) using amino acid sequences. Abietadiene cyclase isolated from *Abies grandis* (AgAS) was chosen as the outgroup and the bootstrap value was set to 1000. All the sequences were aligned by MUSCLE v3.8.31 (www.drive5.com/muscle). JTT + I + G, the best-fit models of amino acid replacement for class I and class II diterpene synthase gene alignments were selected by ProtTest v3.4.2 (github.com/ddarriba/prottest3) considering models with a proportion of invariable sites and rate variation among sites and number of categories.

#### ***Positive selection analysis***

In terms of syntenic and phylogenetic relationships, genes which had been characterized as encoding *ent*-copalyl diphosphate synthase, *ent*-labda-13-en-8-yl diphosphate synthase, kolavenyl diphosphate synthase and isokolavenyl diphosphate synthase in previous research and this study were aligned by MUSCLE using amino acid sequences. The reverse translation from protein alignment to codon alignment was performed by RevTrans v1.4 (www.cbs.dtu.dk/services/RevTrans). Based on Maximum Likelihood method, CodeML estimates dN/dS value on the codon alignment. The background branches shared the same distribution of dN/dS value among sites, whereas different values could apply to the foreground branches which comprise kolavenyl diphosphate synthase and isokolavenyl diphosphate synthase genes. For the alternative model and null model, dN/dS ratio was fixed to 0 and 1 individually. *P* value for the dN/dS value of the foreground branch was tested using a chi-squared test with an inbuilt script $\chi^2$.

#### ***Protein sequence alignments***

Protein sequence alignments were generated using MUSCLE for diterpene synthases encoded by genes in each microsynteny. The secondary structure elements from AgAS crystal structure accompanied by alignments were supplied to ESPript v3.0 (escript.ibcp.fr).

### Molecular Biology and Biochemistry

#### ***Cloning of genes***

Plant tissues collected from leaves, stems, flowers and roots were pooled for RNA extraction using RNeasy Mini Kit (Qiagen) or CTAB method. The extracted RNA was purified with TURBO DNA-free kit (ThermoFisher) to remove gDNA prior to being reverse-transcribed into cDNA with SuperScript IV First-Strand Synthesis System (ThermoFisher). Candidate genes were amplified with either KOD plus Neo (Toyobo) or Phusion High-Fidelity PCR Master Mix (ThermoFisher) using gene-specific primers (Table S19). The primers for the genes were designed based on genomic and transcriptomic data. Recombinant pTRBO vectors containing candidate genes were constructed by In-Fusion cloning using ClonExpress II One Step Cloning (Vazyme). Transformed DH5a cells were screened on kanamycin selective media (50 mg L<sup>-1</sup>) and colonies that showed the correct insert size in colony PCR were picked for overnight incubation in 5 mL kanamycin selective liquid Luria-Bertani media. These cultures were used for plasmid extraction using TIANprep Mini Plasmid Kit (Tiangen), and the extracted plasmids were sent for sequencing.

#### ***Expression of diterpene synthases in Nicotiana benthamiana***

The recombinant constructs of pTRBO vectors carrying the full-length coding sequences of diterpene synthases (diTPSs) from *Scutellaria barbata*, *Scutellaria baicalensis* and *Salvia splendens* were transferred to *Agrobacterium tumefaciens* strain GV3101 and introduced into *Nicotiana benthamiana* by agro-infiltration.

Transformed *Agrobacterium tumefaciens* colonies were identified by PCR, and a positive colony was cultured in 10 mL LB containing 50 µg/ml rifampicin, 20 µg/ml gentamycin and 50 µg/ml kanamycin and grown for 2 days at 200 rpm, 28°C. Cells were centrifuged at 4000 rpm for 15 min, then resuspended in 1 × MMA (10 mM MgCl<sub>2</sub>, 10 mM MES, 150 μM acetosyringone). All transformed *Agrobacterium tumefaciens* suspensions were normalized to OD<sub>600</sub> around 0.9 and kept in dark for at least one hour. Strains of *Agrobacterium tumefaciens* harboring diTPS genes were mixed with the strain carrying the p19 gene silencing suppressor for co-infiltration in *Nicotiana benthamiana*. Co-infiltrations with empty vector pTRBO and p19 were used as the negative control.

The agrobacterium suspensions were infiltrated into whole leaves of 4-6 weeks old *Nicotiana benthamiana* plants. Plants were grown for further 5 days in the greenhouse before extraction. The infiltrated whole leaves were collected and transferred into Eppendorf tubes (2 ml) and ground thoroughly. Diterpenes were extracted with 1 mL n-hexane (Sigma-Aldrich, ReagentPlus(R) grade) at 37°C overnight in an orbital shaker at 220 rpm. The tubes were vortexed vigorously for 1 min and then centrifuged at 8000 rpm for 10 min. The supernatant was transferred into a new glass tube. The pellets were then extracted in 1 mL n-hexane (Sigma-Aldrich) after 30 min of sonification in room temperature twice. The combined extracted product was dried by nitrogen gas and resuspended in 1 mL n-hexane, then filtered through an organic filter. The resuspension was dried again and dissolved in 50 μL n-hexane with 1 mg/L 1-eicosene as the internal standard (IS) and aliquoted to GC vials for analyses. Three biological replicates of each combination were analyzed.

##### ***Recombinant constructs for protein expression in E. coli***

The length of chloroplast signal peptide in diterpene synthases was predicted based on ChloroP (http://www.cbs.dtu.dk/services/ChloroP/). Then the truncated versions of diterpene synthases were cloned into either pOPINF or pOPINM vector (Table S17) by In-Fusion cloning using ClonExpress II One Step Cloning (Vazyme) for expression in *E. coli*. Colonies were screened by colony PCR to check the size of the inserted gene, and those which had the correct band size were sent out for sequencing. Expression vectors pOPINF or pOPINM carrying the N-terminal truncated versions of diterpene synthases were transformed into *E. coli* strains (Table S17) for protein expression.

##### ***Protein expression and purification***

LB media containing selective antibiotics were inoculated with selected recombinant *E. coli* colonies and cultured overnight at 220 rpm, 37 °C. This overnight culture was used to inoculate Terrific-Broth media containing the same selective antibiotics at 1:50 dilution. The TB culture was incubated at 37 °C and shaken at 220 rpm until the optical density at 600 nm reached 0.8-1.0. Cultures were cooled to 16 °C for 30 min and induced with 1 mM isopropylthio- $\beta$ -galactoside. Cultures were incubated at 16 °C for a further 16 h before centrifugation. The pellets were resuspended in chilled lysis buffer (50 mM tris-HCl buffer, 50 mM glycine, 5% v/v glycerol, 0.5 M NaCl, 20 mM imidazole, 0.2 mg mL<sup>-1</sup> lysozyme and 1 mM phenylmethylsulfonyl fluoride or Complete Protease Inhibitor Cocktail [Roche], pH=8) and kept on ice for 30 min. The lysate was disrupted by sonication (Scientz JY92-IIN) and centrifuged at 4 °C (35000 rpm, 20 min). A slurry of 200-250 μL Ni NTA beads (Smart LifeSciences) was added into the collected supernatants to pull down the His-tagged protein. After gently shaking the mixture on a rocking platform at 4 °C for 1.5 h, the mixture was centrifuged and washed with ice cold lysis buffer (that did not contain protease inhibitor or lysozyme) twice. Proteins were eluted with 600 μL elution buffer (50 mM tris-HCl buffer pH=8, 50 mM glycine, 5% v/v glycerol, 0.5 M NaCl, 0.5 M imidazole) and the eluate were applied to a 4 mL Amicon Centrifugal 30k NMWL tube for dialysis, where the buffer content was exchanged to phosphate-buffered saline (0.137 M NaCl, 0.0027 M KCl, 0.01 M phosphate buffer, pH = 7.4). Protein concentration was measured by the A<sub>280</sub> using Nanodrop and diluted to approximately 1 mg mL<sup>-1</sup> to avoid precipitation. Purified enzymes were divided into aliquots, snap frozen in liquid N<sub>2</sub> and stored under -80 °C.

##### **In vitro enzyme assays of diterpene synthases**

Enzyme assays were carried out in 250 μL assay buffer consisting of 50 mM tris-HCl (pH = 7.2), 7.5 mM MgCl<sub>2</sub>, 100 mM KCl, 5% glycerol and 5mM dithiothreitol. The reactions were started by adding 50 μg class II diTPS alone or in combination with 50 μg class I diTPS into the buffer and incubating it with 100 μM GGPP at 37 °C for 2 h in the dark. In the enzyme assays of class II diTPS, the enzyme products were further dephosphorylated with 3 μL 30 U calf intestinal alkaline phosphatase (Takara Bio) at 37 °C for 2 h before extraction. The products were extracted with 250 μL n-hexane repeatedly, three times, using equal volumes of hexane, added every time. The hexane fractions were pooled and evaporated under N<sub>2</sub> flow, then resuspended in 30 μL hexane for GC-QTOF analysis.

##### **Cytochrome P450s expression in yeast**

Plasmids for expression of GGPPS (Erg20(F96C)), MS, CPS and Cytochrome P450s were transformed into the yeast strain AM119 (Ignea et al., 2016), and were grown on selected synthetic dropout (SD) medium. The recombinant yeast strains were cultured in 10 mL liquid SD-His-Leu or SD-His-Leu-Ura medium supplemented with 2% glucose at 30°C for 24 h with shaking. The cultures were centrifuged and the cells were washed twice with sterile ddH<sub>2</sub>O; The cells were resuspended in 20 mL liquid SD-His-Leu or SD-His-Leu-Ura medium supplemented with 2% galactose and agitated for 48 h at 30°C. Yeast cultures were centrifuged and the cells were ground by tissue lyzer in tubes containing glass balls, then the cracked cells and the supernatant were extracted three times with 10 mL hexane. After treatment with ultrasonication and separation, the organic extracts were dried using a rotary evaporator, and dissolved in hexane for GC-MS analysis.

#### **Determination of the content of Scutebarbatine A**

Different tissues (roots, flowers, young stems, old stems, leaves (0.5<leaf width<1 cm)) were ground and then dissolved in methanol (1mg per 20 μL methanol). Standard compound, scutebarbatine A, was purchased from Purifa (Chengdu, China).

The content of scutebarbatine A in different tissues was determined by AB 5500 Q-TRAP (Massachusetts, USA) using Poroshell 120 SB-C18 2.7 μm (3×100 mm) in Shanghai Institute of Plant Physiology and Ecology. Scutebarbatine A was dissolved in methanol and set at different concentrations (100 ng/mL; 50 ng/mL; 25 ng/mL; 10 ng/mL; 5 ng/mL; 1 ng/mL; 0.5 ng/mL; 0.1 ng/mL) for calibration curves. The analyses were performed using three biological replicates.

Separations were conducted using the following gradient program: 60%A+40%B for 0 to 8 min; 5%A+95%B for 8 to 10 min; 1%A+99%B for 10 to 10.1 min; 60%A+40%B for 10.1 to 12 min (Mobile phase A: 2 mM NH<sub>4</sub>FA+ 0.01% FA; mobile phase B: acetonitrile). The flow rate was 0.4 mL/min.

#### **qRT-PCR**

The tissues (roots, young stems, old stems, flowers, old leaves (leaf width>2 cm), young leaves (0.5<leaf width<1 cm), leaves without trichomes (0.5<leaf width<1 cm), large trichomes (>100μm diameter) of *S. barbata* were collected at the flowering stage, and each tissue was analysed using three biological replicates. Total RNA was extracted by the CTAB method. qRT-PCR experiments were run on an AB StepOne Plus system using SYBR Premix ExTaq Mix (Takara), and relative expression levels were calculated as described (Li et al., 2020), using *SbbqACTIN1* as the reference gene. The primers are shown in Table S19.

### GC-MS analysis

The GC-MS analysis of diterpenes was based on a method described previously (Andersen-Ranberg et al., 2016). The samples were analyzed on a GC7890B-MS7200B QTOF (Palo Alto, California, USA) using an Agilent DB-5HT (30 m × 0.25 mm × 0.1 μm) in the Shanghai Institute of Plant Physiology and Ecology and Shanghai Institute of Organic Chemistry, respectively. Injection volume was 2 μL for transfected *Nicotiana benthamiana* samples and 5 μL for enzyme reaction samples and samples were injected in split mode (split ratio, 5:1), the temperature was set to 250°C. The GC program was 50°C hold 2 min, 4°C min<sup>-1</sup> to 110°C, 8°C min<sup>-1</sup> to 250°C, 10°C min<sup>-1</sup> to 310°C and hold for 5 min. The temperature of the MS (mass spectrometer) for ion source was set to 230°C and spectra were recorded from m/z 50 to m/z 350. The flow rate was 1 mL/min.

### Chemical methods

#### Experimental procedures

Reactions were carried out under nitrogen atmosphere and the glassware was oven-dried prior to use. Double distilled water was used for the preparation of all buffers and solutions, which were freshly prepared preceding use. NMR samples were prepared with deuterated solvents and measured in Bruker AVANCE III 400MHz or 500MHz.

#### Synthesis of Geranylgeranyl pyrophosphate (GGPP)

To a stirring solution of geranylgeraniol (0.2g, 0.68 mmol) in dry Et<sub>2</sub>O (10 mL) at 0 °C, PBr<sub>3</sub> (0.02 mL, 0.27 mmol) was added. The mixture was allowed to warm up to room temperature and was kept stirring under N<sub>2</sub> atmosphere until completion, as monitored by thin layer chromatography (TLC) every 30 minutes. The reaction mixture was washed with brine, organic phase was dried with Na<sub>2</sub>SO<sub>4</sub>, filtered, and concentrated under reduced pressure. The thick yellow liquid was then used directly in the next step. To the crude geranylgeranyl bromide in dry acetonitrile (Bu<sub>4</sub>N)<sub>3</sub>HOPP was added. The reaction was kept stirring at room temperature and under N<sub>2</sub> atmosphere until completion – around 2 hours. The reaction mixture was then dried under reduced pressure (bath temperature <40 °C) and the thick liquid was then loaded onto a column containing Dowex 50WX8-200 resin (NH<sub>4</sub><sup>+</sup> form), previously equilibrated with 2-propanol/25 mM aq. NH<sub>4</sub>HCO<sub>3</sub> (1:49, v/v) and product was eluted with this same mobile phase. Product was freeze-dried and its purity was checked by TLC. When necessary, column chromatography was performed in spherical silica cartridge pre-equilibrated with 2-propanol containing 1% aq. $\text{NH}_4\text{HCO}_3$ 0.1 M (mobile phase: 1% to 80% $\text{NH}_4\text{HCO}_3$ 0.1 M in isopropanol). Fractions containing the product (as checked by TLC, spotted with sulfuric anisaldehyde) were pooled together, bulky solvent was evaporated and remaining aqueous solution was freeze-dried to give a yellow syrup with 31% yield as $(\text{Bu}_4\text{N})\text{NH}_4$ salt, seen by $^1\text{H}$ NMR. NMR data are in accordance with literature (Chow et al., 2005). The concentration of GGPP per milligram of salt was measured by quantitative NMR using DSS as reference.

**GGPP, $^1\text{H}$-NMR $\delta\text{H}$ (ppm) ($\text{CDCl}_3$; 400 MHz):** 5.30 (1H, t, $J$= 7.2 Hz H-2), 5.12 (2H, m, H-6, H-10), 4.80 (1H, m, overlapped, H-14), 4.40 (2H, t, $J$= 6.5 Hz, H-1), 2.10-1.80 (10H, m, H-4, H-5, H-8, H-9, H-12, H-13), 1.60-1.40 (15H, 5s, 5 Me).

#### **Isolation of miltiradiene as product of an enzymatic reaction**

The SbbdiTPS2.4 with CPS activity and the SbbdiTPS1.1 with MS activity, were co-expressed together in *E. coli* using the pIRS and pGG vectors (Jia et al., 2019) to isolate miltiradiene. To the crude enzyme assay, 50 mL of hexane was added, and the mixture stirred overnight. The organic and aqueous phases were separated, and the organic phase was dried with $\text{Na}_2\text{SO}_4$ and concentrated, giving 12 mg of crude extract. The mixture was loaded into 12g spherical silica cartridge and flash chromatography using hexane as solvent was performed. Miltiradiene was isolated and subjected to NMR analysis, which was in accordance with literature (Božić et al., 2015)

**Miltiradiene, $^1\text{H}$-NMR $\delta\text{H}$ (ppm) ($\text{CDCl}_3$; 500 MHz):** 5.45 (1H, m, H-12); 2.63 (2H, m, H-14); 2.49 (1H, m, H-11a); 2.41 (1H, m, H-11b); 2.18 (1H, m overlapped, H-15); 1.99 (2H, m, H-7); 1.73 (3H, m, H-1a, H-2a, H-6b); 1.52-1.44 (3H, m, H-2b, H-3, H-6a); 1.20 (1H, dd, $J$= 12.5Hz, 2Hz, H-5); 1.16 (2H, dd, $J$= 13.5 Hz; 4.1 Hz, H-1b, H-2b); 1.2 (3H, d, $J$= 6.8 Hz, Me); 1.01 (3H, d, $J$= 6.8 Hz, Me-17); 0.99 (3H, s, Me-16); 0.89 (3H, s, Me-18); 0.86 (3H, s, Me-19). $\delta\text{C}$ (ppm) ($\text{CDCl}_3$; 500 MHz) (assigned by HSQC): 116.5 (C-12); 51.4 (C-5); 41.7 (C-3); 36.9 (C-1); 34.0 (C-15); 33.3 (C-11); 33.3 (C-18); 31.9 (C-7); 25.3 (C-14); 21.6 (C-19); 21.3 (C-17); 21.2 (C-16); 18.9 (C-6); 19.2 (C-2); 18.6 (C-6).

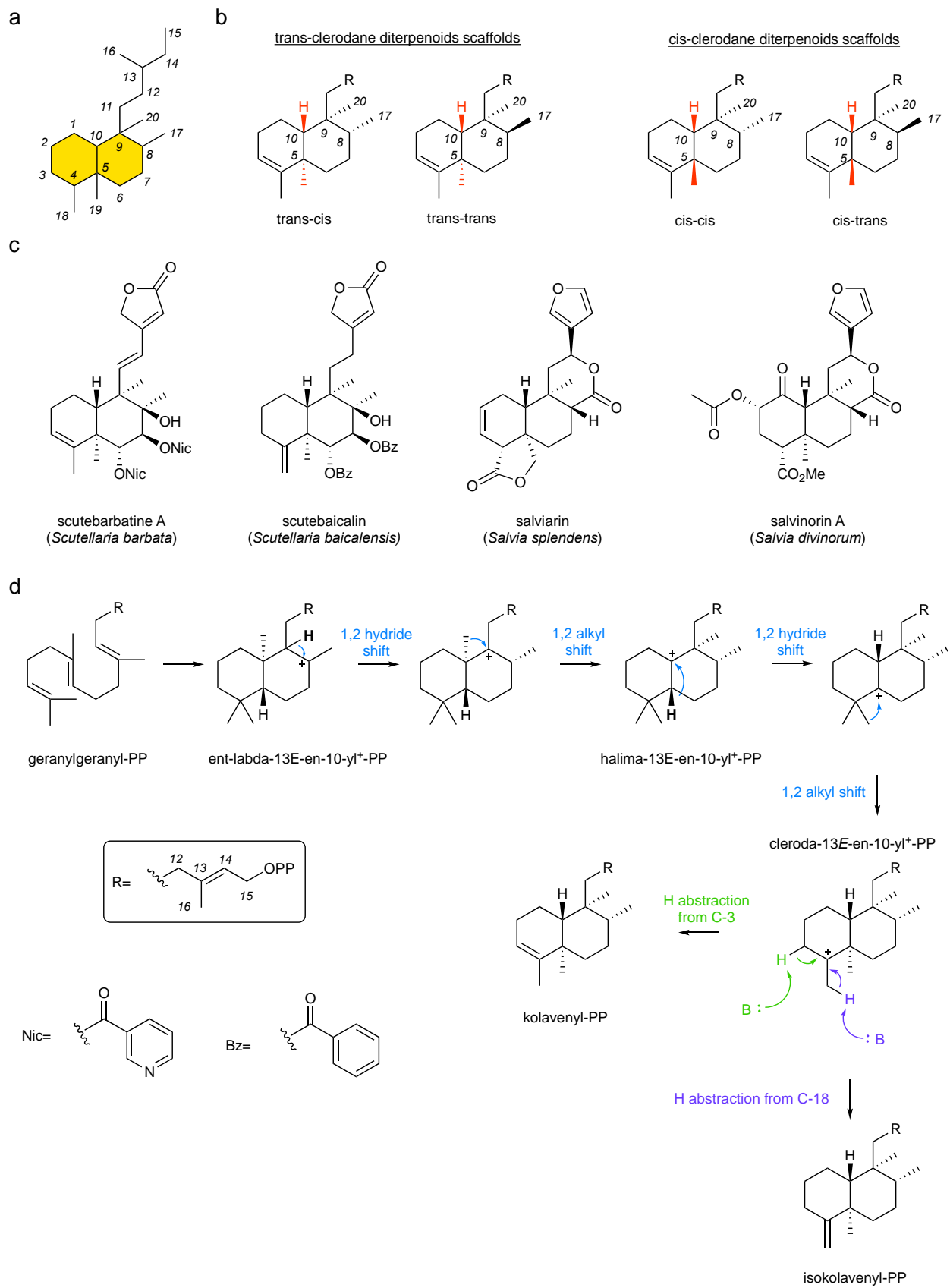

**Fig. S1. The basic structural scaffold of clerodane diterpenoids and enzymatic synthesis by class II diterpene synthase. a:** the clerodane carbon skeleton with all the carbons numbered. **b:** structural classification of clerodanes based on stereochemistry of basic skeleton. **c:** the structures of scutebarbatine A from *Scutellaria barbata*, scutebaicalin from *Scutellaria baicalensis*, salviarin from *Salvia splendens* and salvinorin A from *Salvia divinorum*. **d:** proposed catalytic mechanisms of kolavenyl diphosphate synthase KPS and isokolavenyl diphosphate synthase IKPS in Lamiaceae depicting the cyclization followed by a cascade of 1,2-hydride and methyl shifts for the formation of the clerodane scaffold.

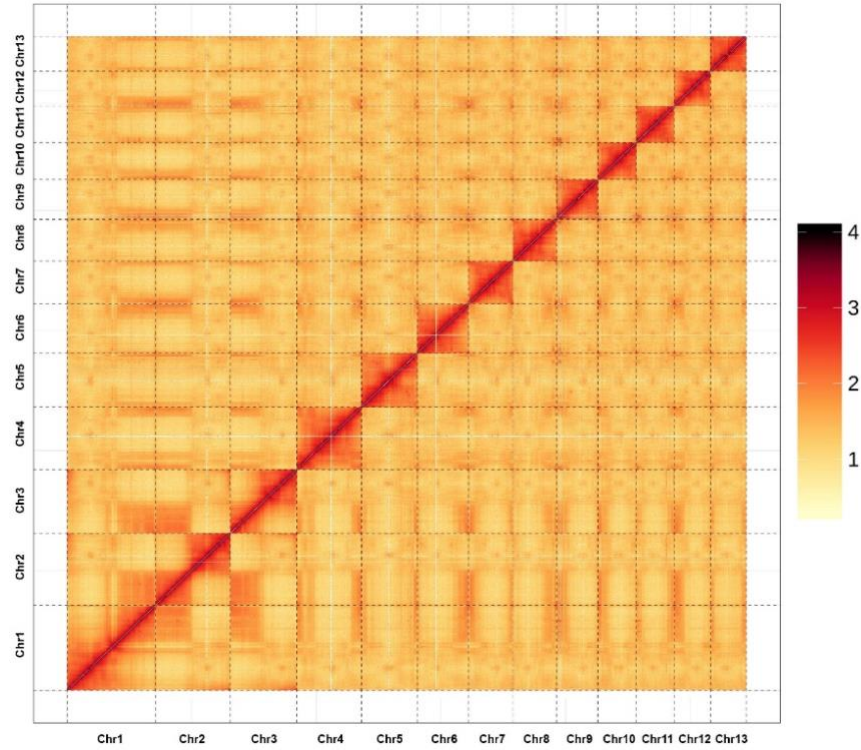

**Fig. S2. Hi-C contact frequencies within and among *S. barbata* pseudochromosomes (Chr1-Chr13).** The color bar indicates contact frequencies from low (yellow) to high (red).

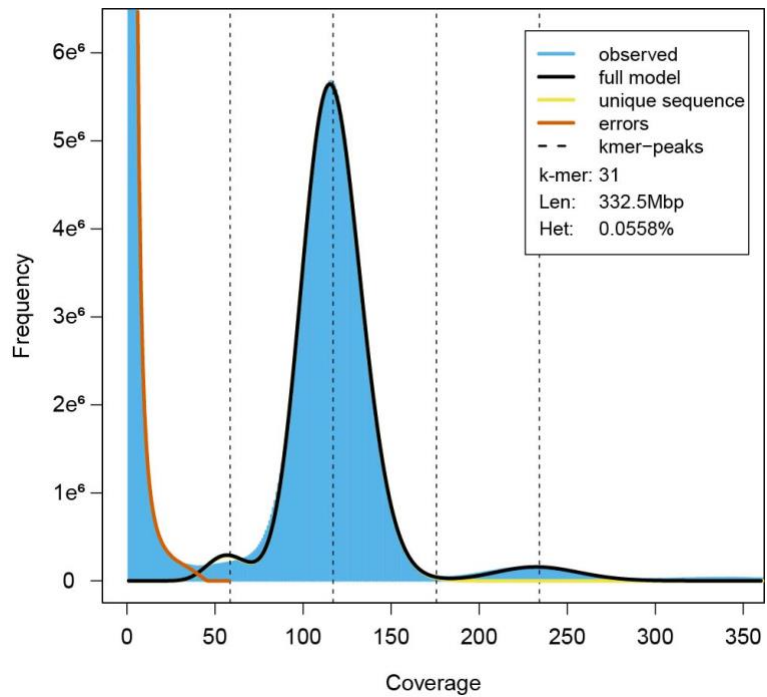

**Fig. S3. Evaluation of *Scutellaria barbata* genome size by k-mer analysis.** Distribution of k-mers was analyzed by GenomeScope1. The major peak is the homozygous portions of the *S. barbata* genome. The tiny shoulder to the left of the major peak corresponds to the heterozygous portions of the genome which lead heterozygosity rate to 0.0558%. Observed = observed 31-mer distribution; full model = mixture model excluding presumptive sequencing errors; unique sequence = sequence without higher frequency repeats in the genome; errors = sequencing errors identified by low coverage *k-mers* = *kmer-peaks*, four components of the genome including heterozygous unique region, homozygous unique region, heterozygous repetitive region and homozygous repetitive region; Len = inferred total genome length; Het = overall rate of heterozygosity

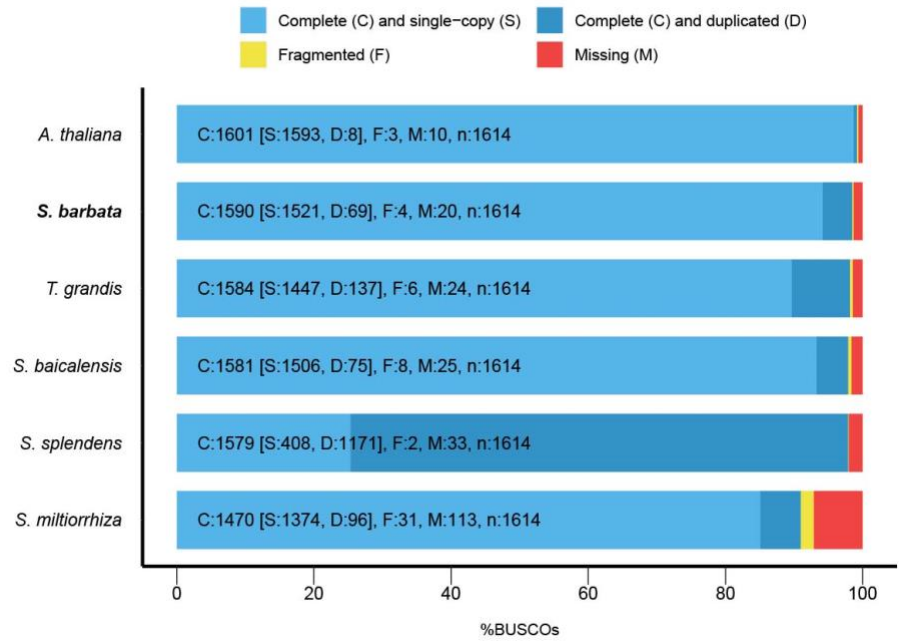

**Fig. S4. Assessment of genome assembly and completeness of annotation.** Benchmarking Universal Single-Copy Orthologs (BUSCO) software was used to evaluate the quality of the genome assemblies among five Lamiaceae species and *Arabidopsis thaliana*.

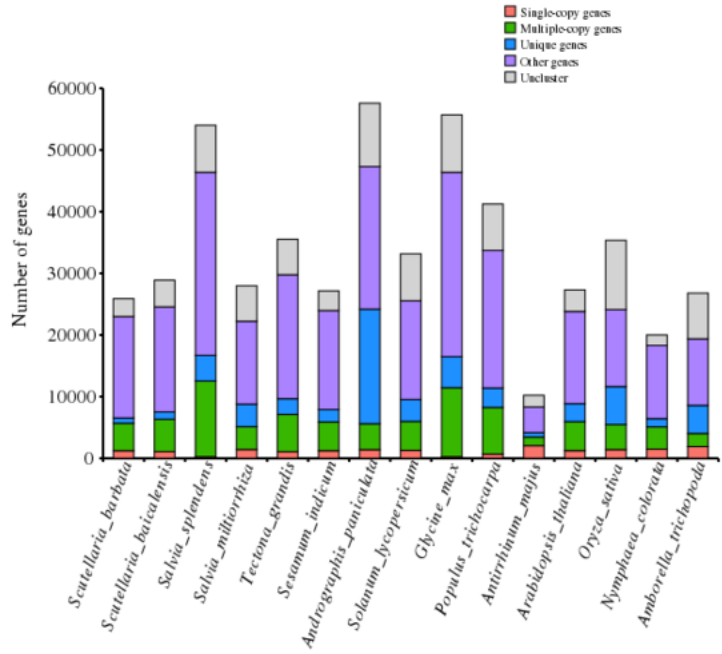

**Fig. S5. Distribution of types of genes in different species.**

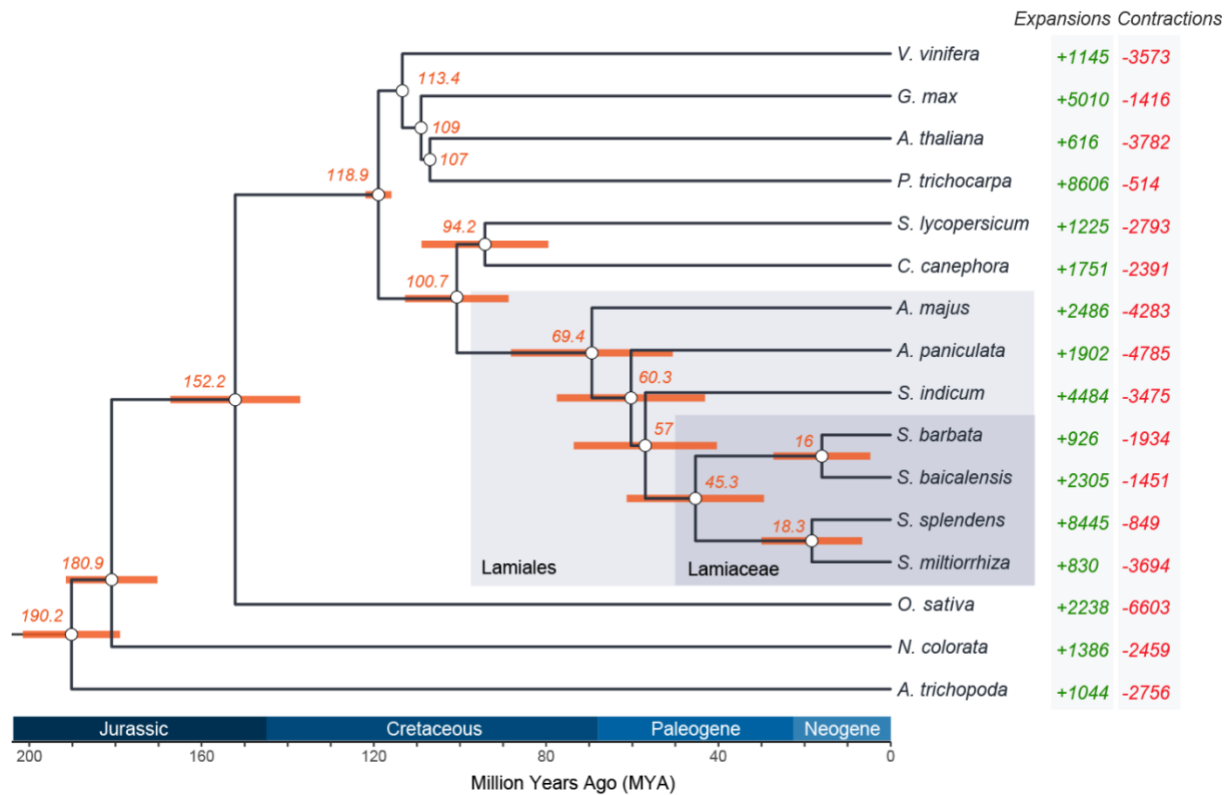

**Fig. S6. Phylogenetic tree inferred from orthogroups in 16 plant genomes.** Estimated divergence times for each node is shown in orange and 95% confidence intervals are represented as orange bars. The support values for all nodes exceed 0.95. Expansions and contractions of gene families are shown in green and red respectively. The light grey box includes the species from the order Lamiales and the species from the Lamiaceae family are highlighted by the dark grey box. The phylogenetic tree was calibrated with the following points as described previously (Kumar et al., 2017): *A. thaliana* and *P. trichocarpa* (107-109 million years ago), *A. thaliana* and *G. max* (107–109 million years ago), *S. lycopersicum* and *P. trichocarpa* (107-125 million years ago), *O. sativa* and *A. thaliana* (140-200 million years ago), *V. vinifera* and *A. thaliana* (114-113 million years ago).

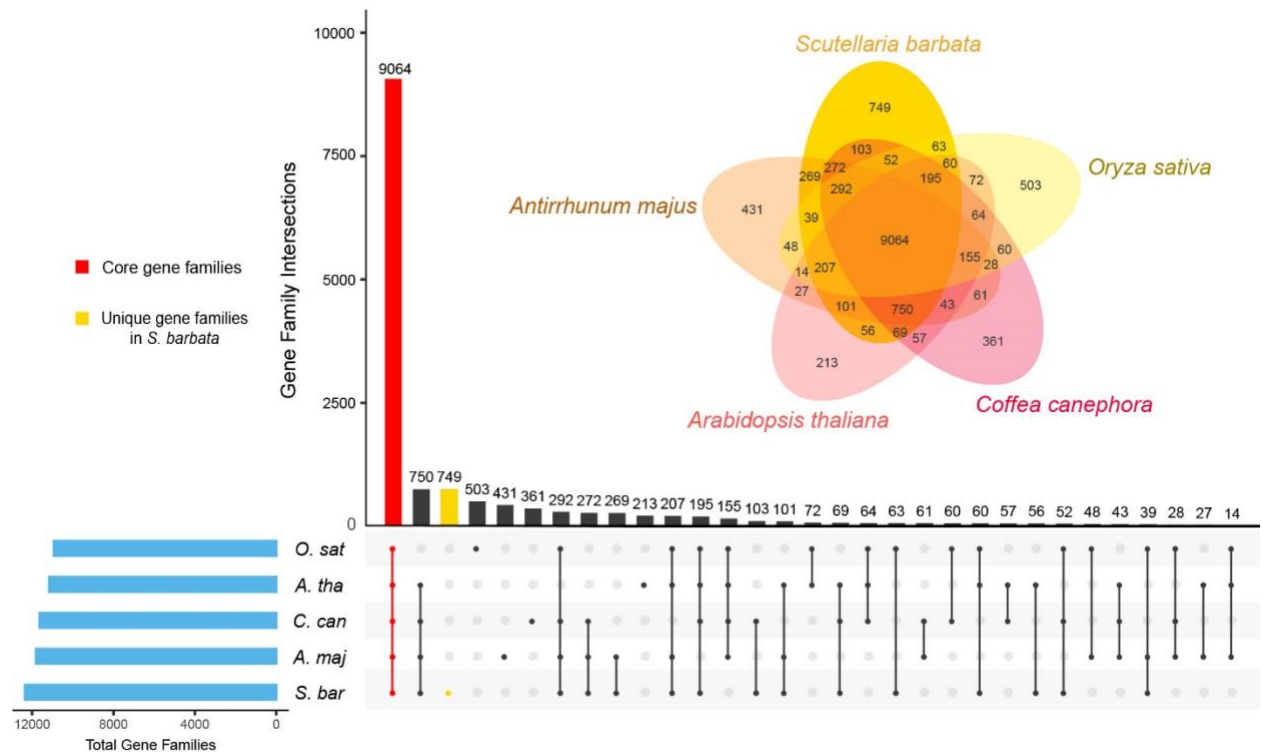

**Fig. S7. Shared and unique gene families among five species.** Upset plot and Venn diagram show the core and unique gene families across *S. barbata*, *A. majus*, *C. canephora*, *A. thaliana* and *O. sativa* genomes. Intersections of overlapping gene families are represented as linked dots and the number of each intersection is indicated by the bar plot.

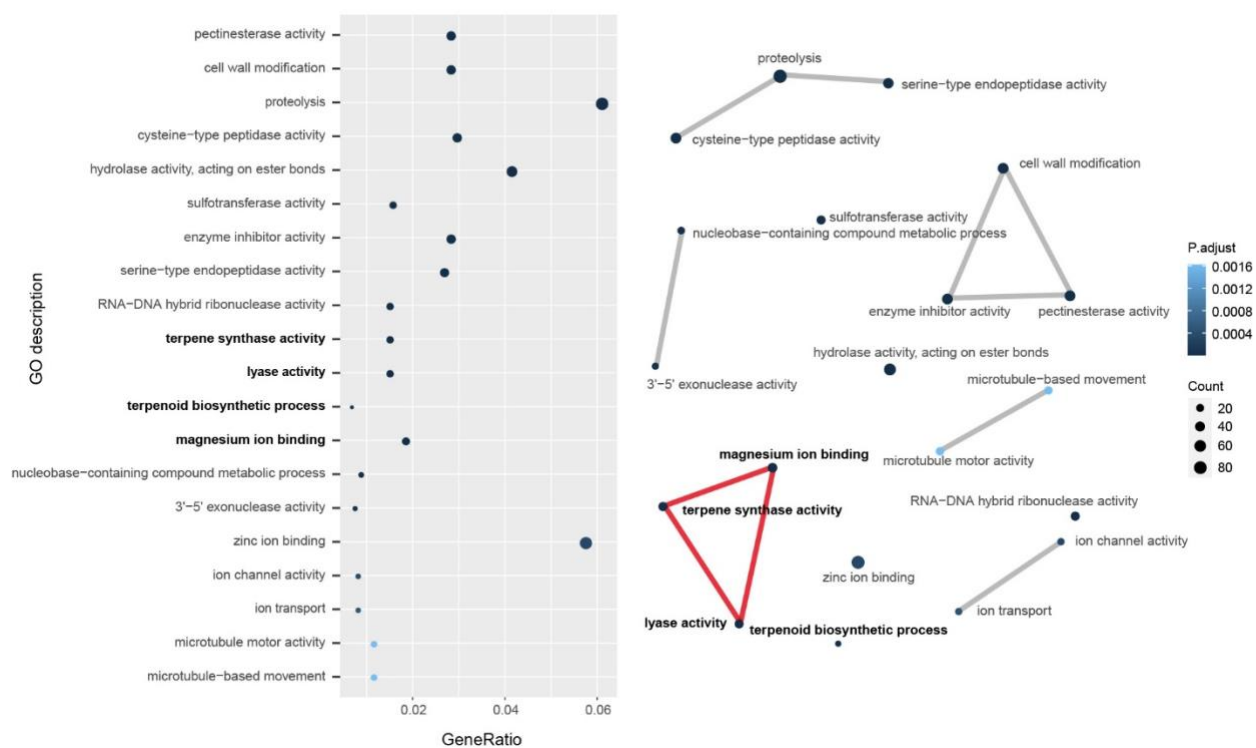

**Fig. S8. Expanding gene family enrichment analysis in *S. barbata*.** (A) The dot plot shows the top 20 enriched Gene Ontology (GO) terms including GO:0010333 (terpene synthase activity), GO:0016829 (lyase activity), GO:0016114 (terpenoid biosynthetic process) and GO:0000287 (magnesium ion binding) using genes from expanded gene families in *S. barbata*. (B) The enriched terms were built into a network with edges connecting overlapping gene sets that cluster as functional modules. The connections of GO terms related to terpene metabolism are highlighted with red.

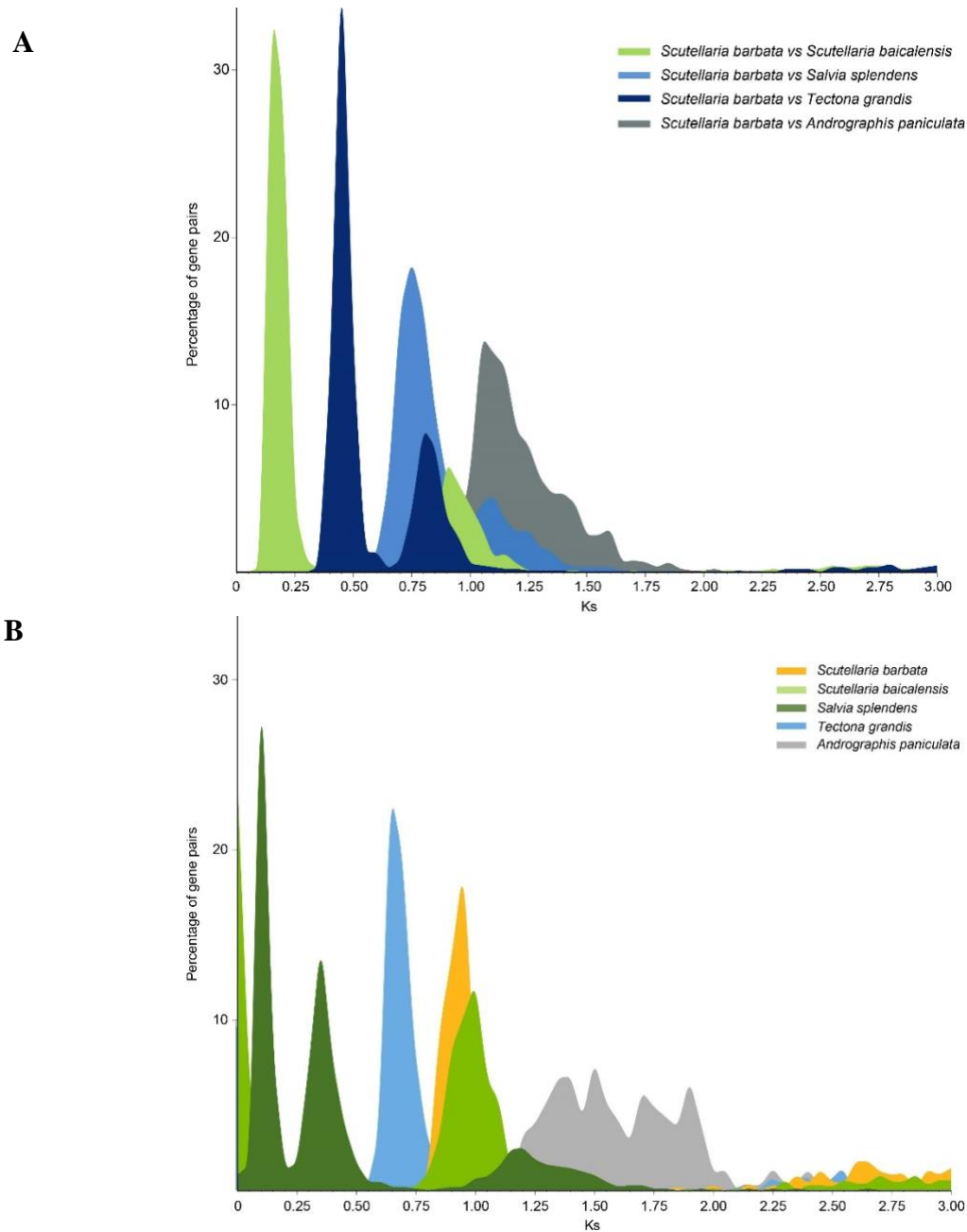

**Fig. S9.** Synonymous substitution rate (Ks) distributions of syntenic blocks for *S. barbata* paralogs and orthologs with other Lamiaceae species (*S. baicalensis*, *S. splendens*, *T. grandis*) and Lamiales species (*A. paniculata*), represented in colored regions as indicated.

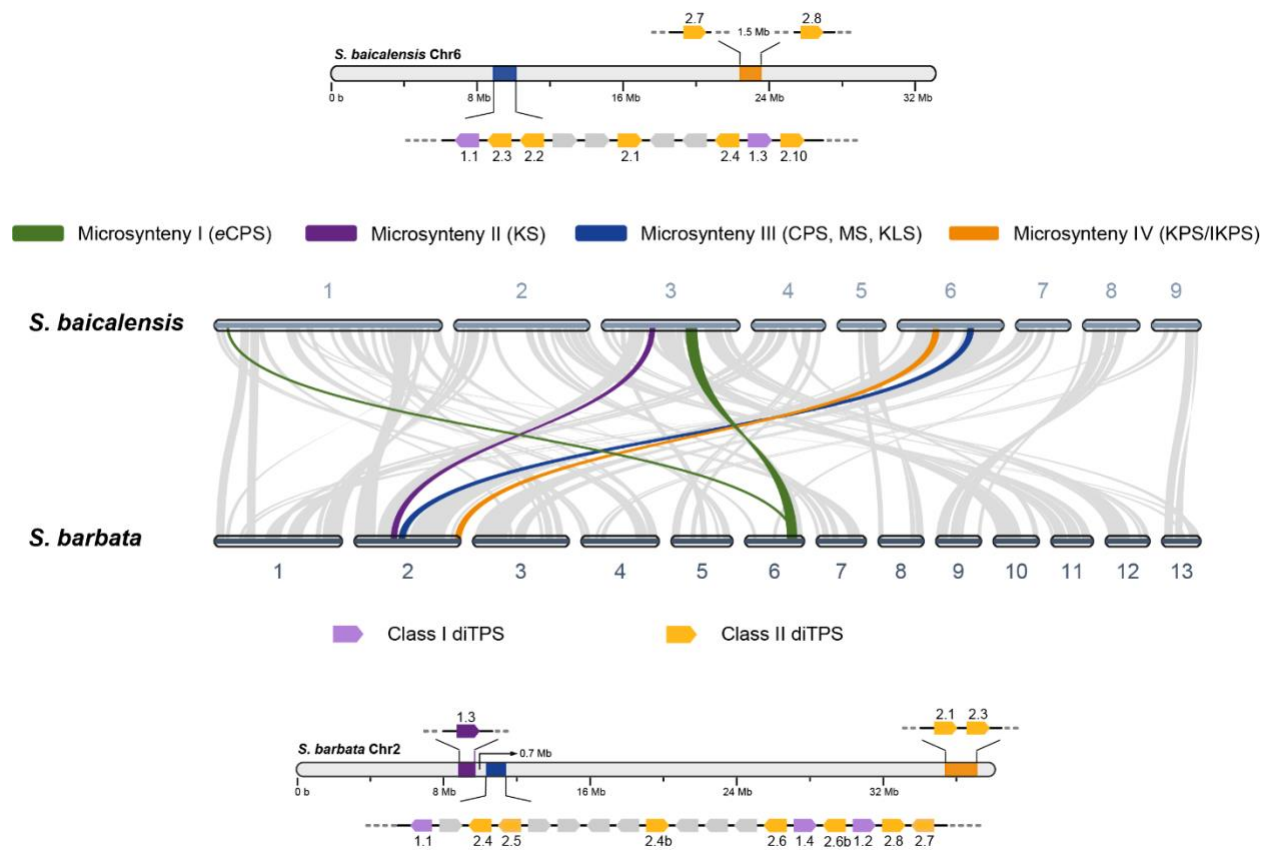

**Fig. S10. Macrosynteny and gene order between *S. barbata* and *S. baicalensis*.** Gray wedges in the background indicate major syntenic blocks spanning $\geq 100$ genes between the genomes, with syntenic regions harboring *ent*-CPS genes, highlighted by green ribbon (microsynteny I), only the class I diTPS KS genes (microsynteny II) is shown in purple ribbon, with both class I (MS, KLS, IKLS) and class II diTPS genes (CPS) shown in blue (microsynteny III). Syntenic regions harboring class II clerodane diTPS genes (KPS and IKPS) are highlighted in orange (microsynteny IV). The position of class I diTPS genes in genomic areas is highlighted light purple while class II diTPS genes are highlighted in gold.

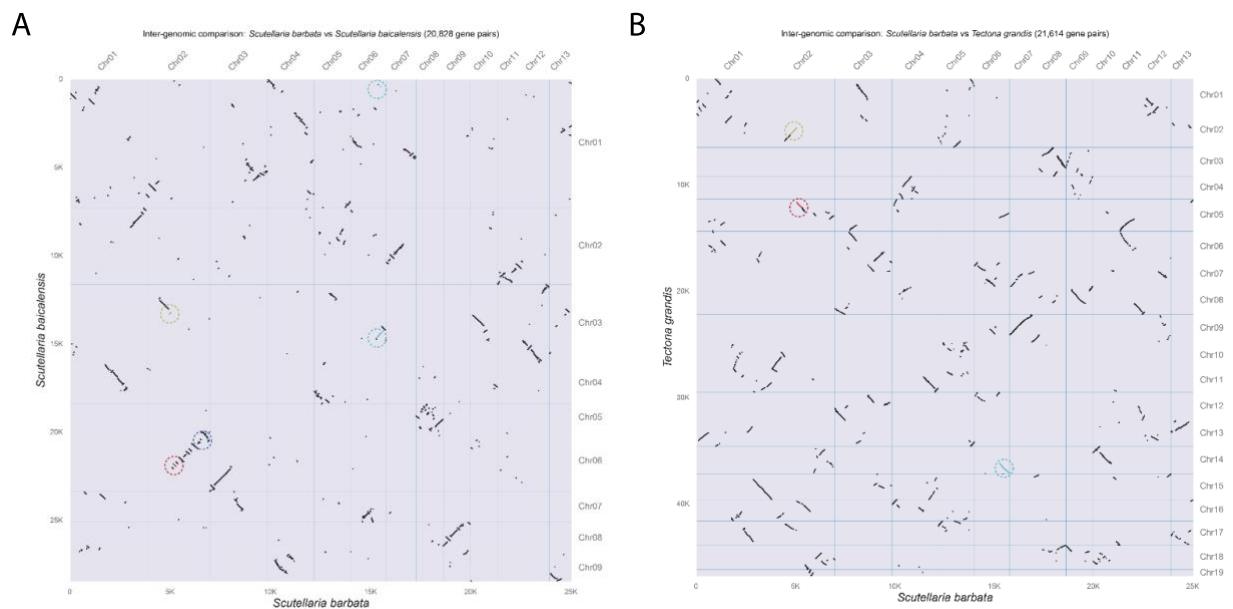

**Fig. S11. Syntenic dot plots between (A) *S. barbata* and *S. baicalensis* and (B) *S. barbata* and *T. grandis*.** The syntenic blocks containing diterpene synthase genes between genomes are highlighted with dash-lined circles.

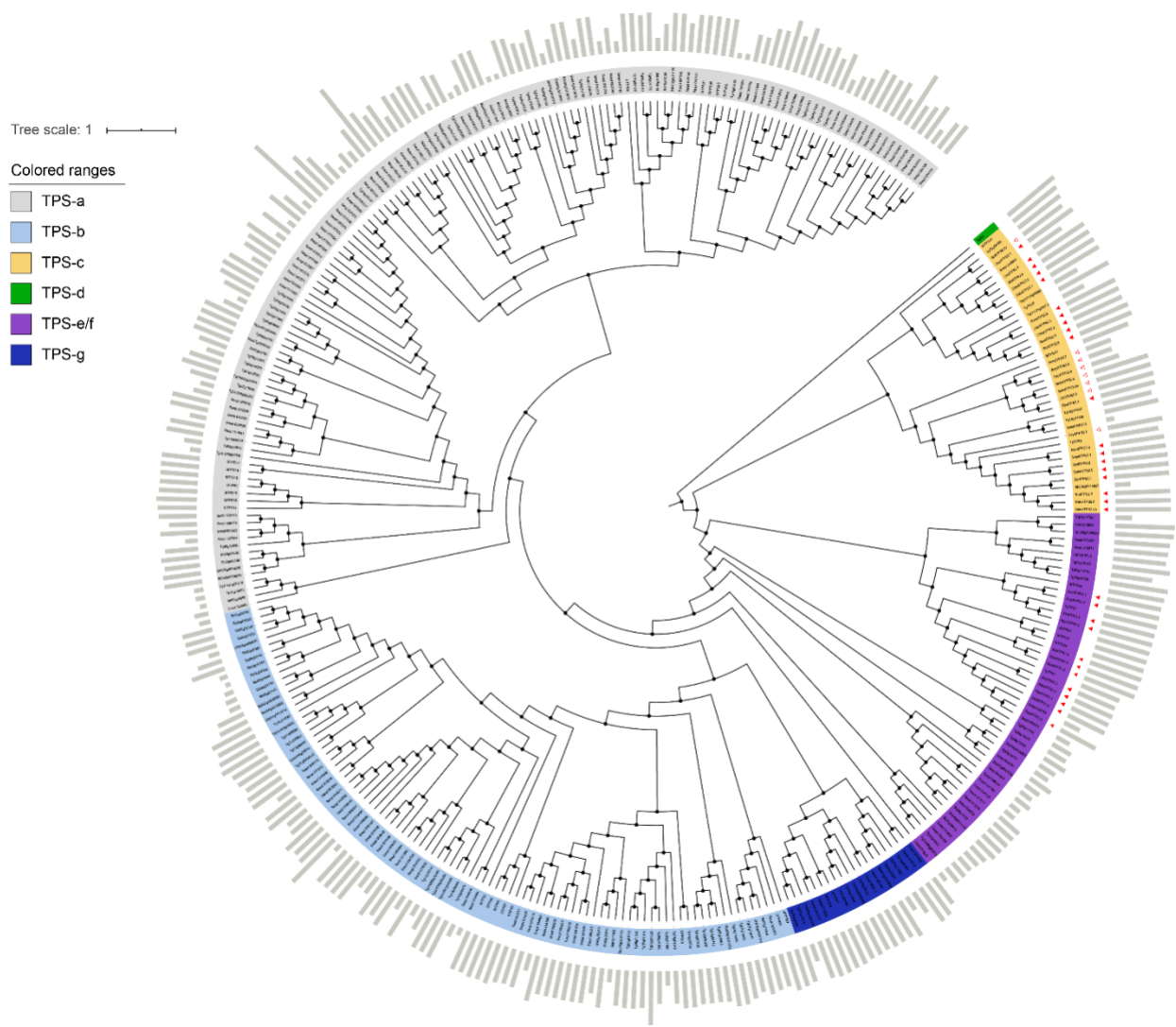

**Fig. S12. Extended phylogeny of functionally annotated terpene synthases (TPS) genes based on relevant Hidden Markov Models (HMM) protein families of terpene synthases (PF01397 & PF03936) from the genomes of *S. barbata*, *S. baicalensis*, *S. splendens*, *T. grandis* and *S. lycopersicum*.** Genes from TPS-a class are highlighted in grey, *TPS-b* in light blue, *TPS-c* in yellow, *TPS-d* in green, *TPS-e/f* in purple and *TPS-g* in navy blue. The red triangles indicate the genes cloned in the present study with the ones filled colour being the gene that were characterized functionally. The outer bars indicate the length of the genes in bp. The bifunctional diterpene synthase (*TPS-d* class, highlighted in green) is abietadiene synthase from the gymnosperm *Abies grandis* (AgAS) which was used as an outgroup. The functionally characterized *TPS* genes from *S. lycopersicum* were used for the categorization of TPS classes.

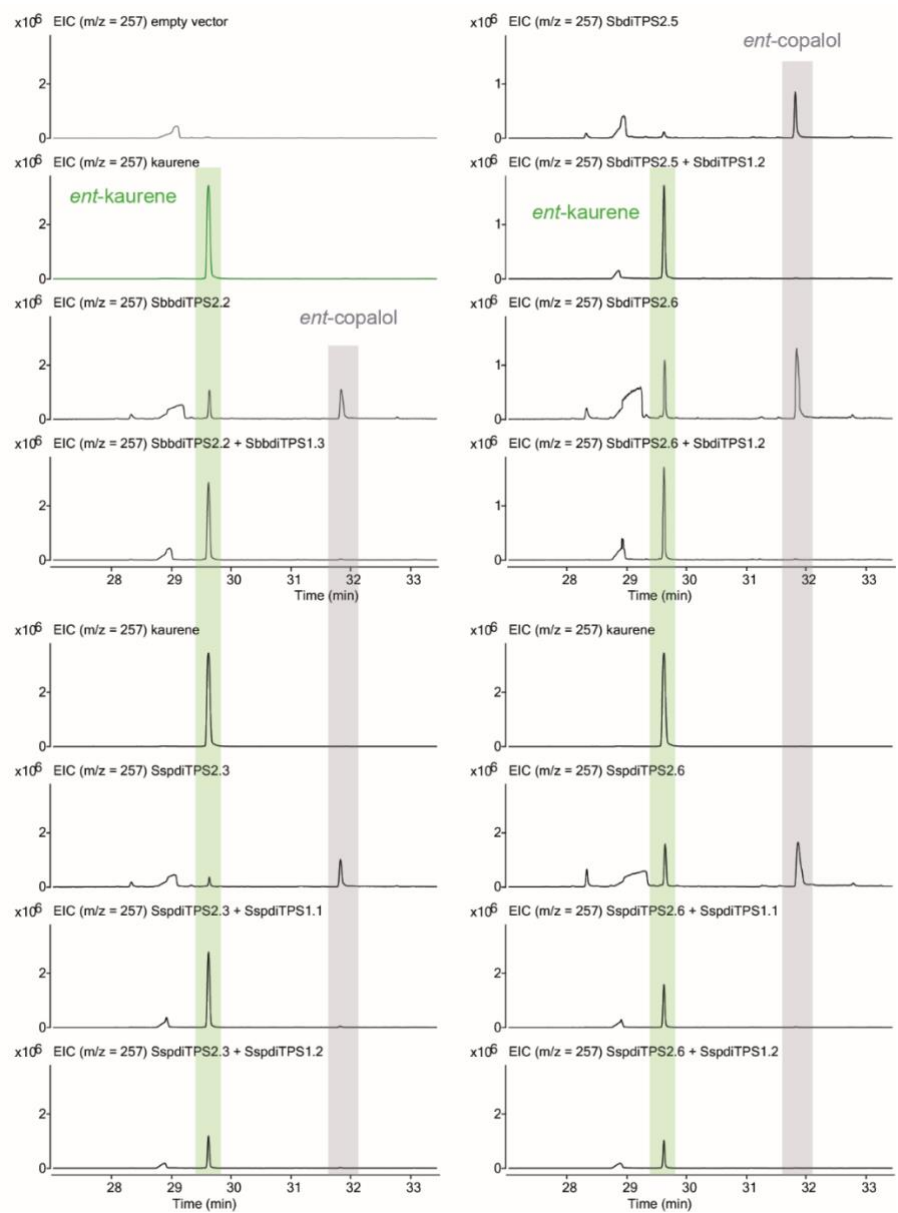

**Fig. S13.** Extracted ion chromatograms at $m/z$ 257 from GC-MS analysis of extracts from *N. benthamiana* leaves expressing class II diTPSs (eCPSs) from *S. barbata*, *S. baicalensis* and *S. splendens* and in combination with class I diTPSs from *S. barbata*, *S. baicalensis* and *S. splendens*. The class I diTPSs acting on *ent*-copalyl diphosphate catalyse the formation of kaurene. The grey background box indicates the presence of *ent*-copalol and the green background box, the presence of kaurene. The formation of *ent*-kaurene was confirmed by comparison with the kaurene standard.

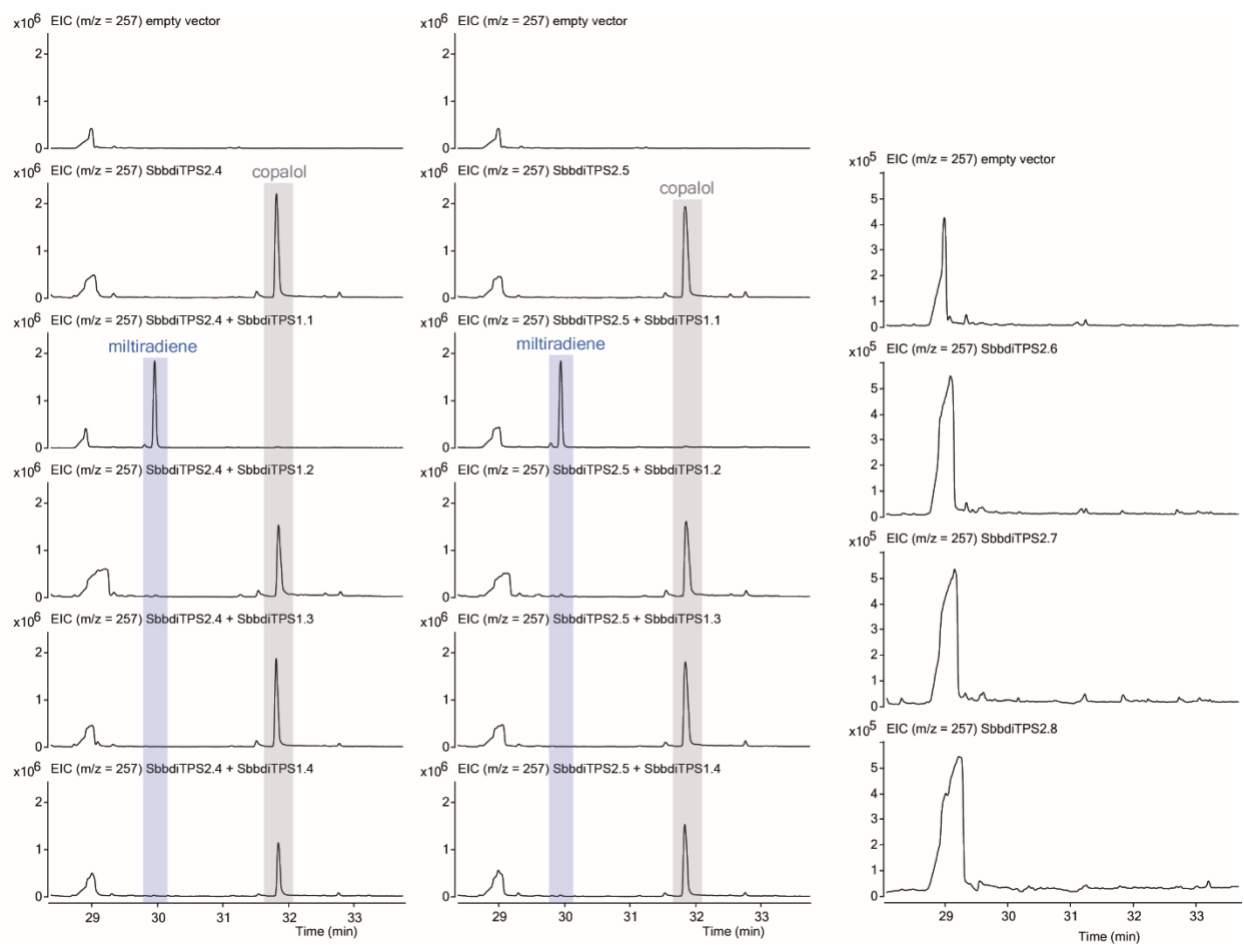

**Fig. S14. Extracted ion chromatograms at $m/z$ 257 from GC-MS analysis of extracts from *N. benthamiana* leaves expressing class II diTPSs (CPSs) from *S. barbata* and in combinations with class I diTPSs from *Scutellaria barbata*.** The grey background box indicates the presence of copalol and the blue background box, the presence of miltiradiene. Miltiradiene was isolated and characterized by NMR.

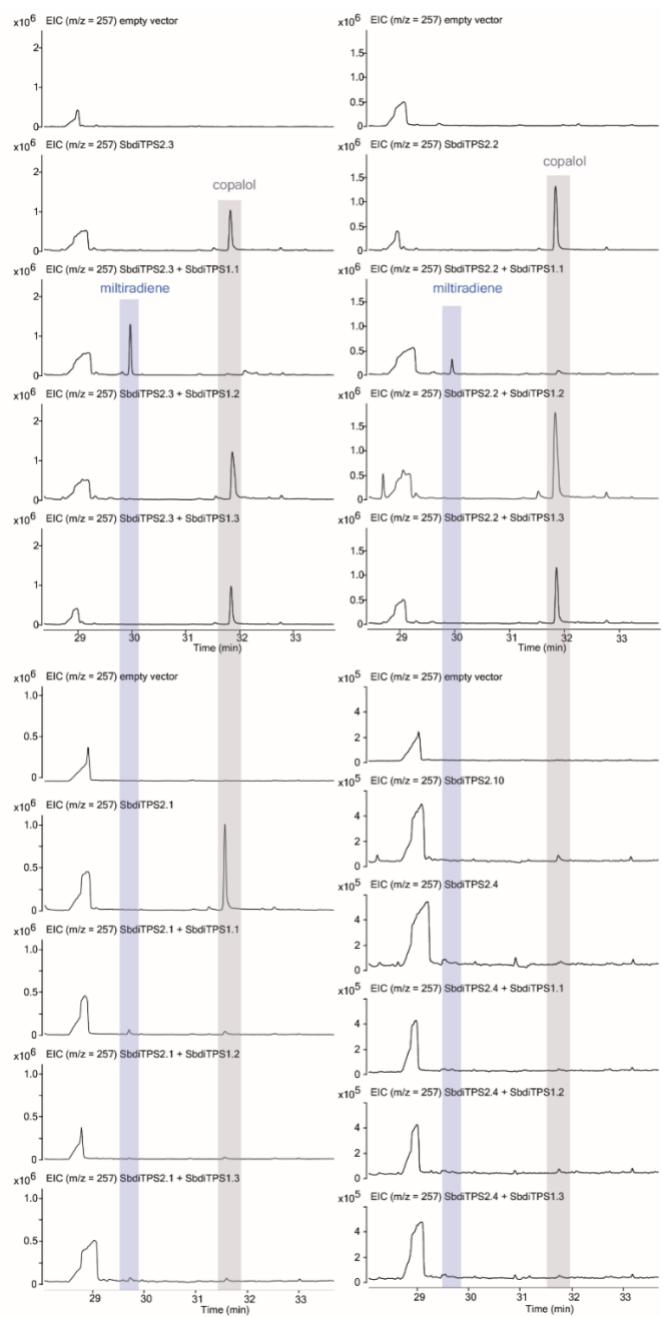

**Fig S15.** Extracted ion chromatograms at $m/z$ 257 from GC-MS analysis of extracts from *N. benthamiana* leaves expressing class II diTPSs (CPSs) from *S. baicalensis* and in combinations with class I diTPSs from *S. baicalensis*. The grey background box indicates the presence of copalol and the blue background box, the presence of miltiradiene. Miltiradiene was isolated and characterized by NMR.

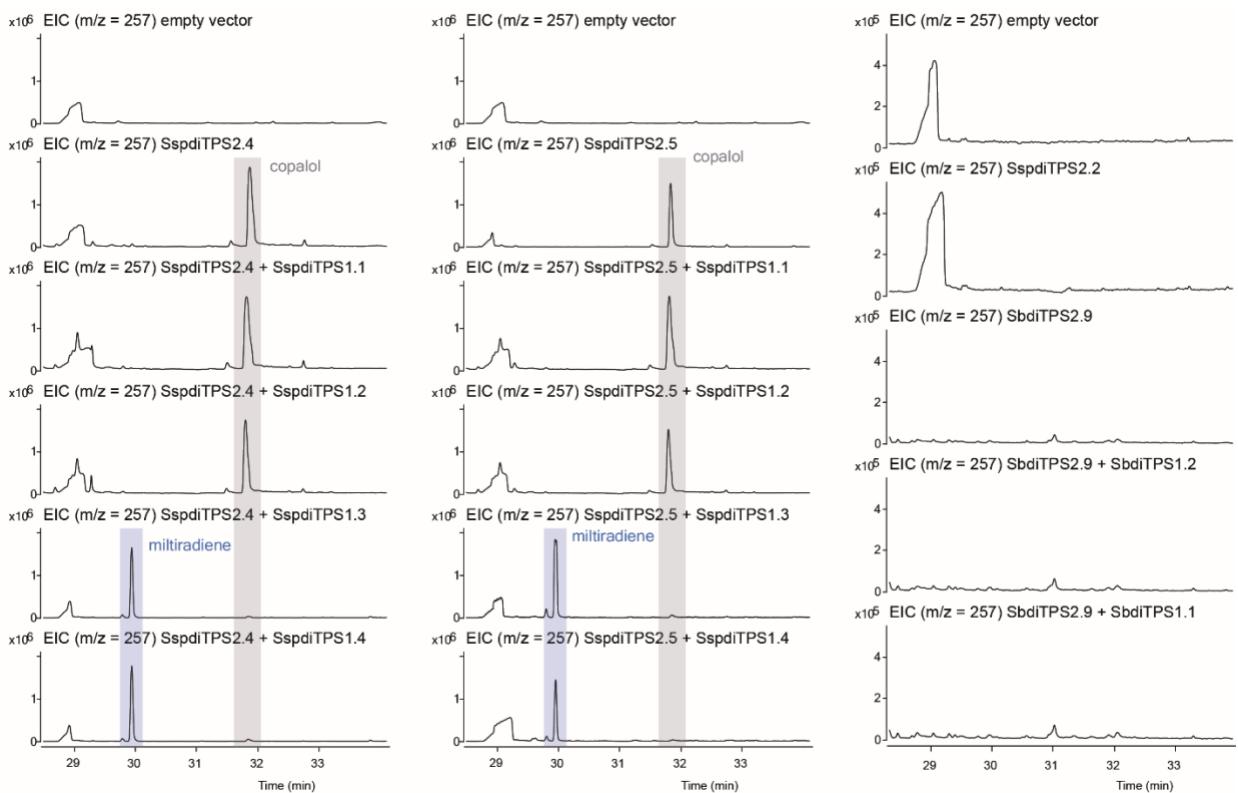

**Fig. S16.** Extracted ion chromatograms at $m/z$ 257 from GC-MS analysis of extracts from *N. benthamiana* leaves expressing class II diTPSs (CPSs) from *S. splendens* and in combinations with class I diTPSs from *S. splendens*. The grey background box indicates the presence of copalol and the blue background box, the presence of miltiradiene. Miltiradiene was isolated and characterized by NMR.

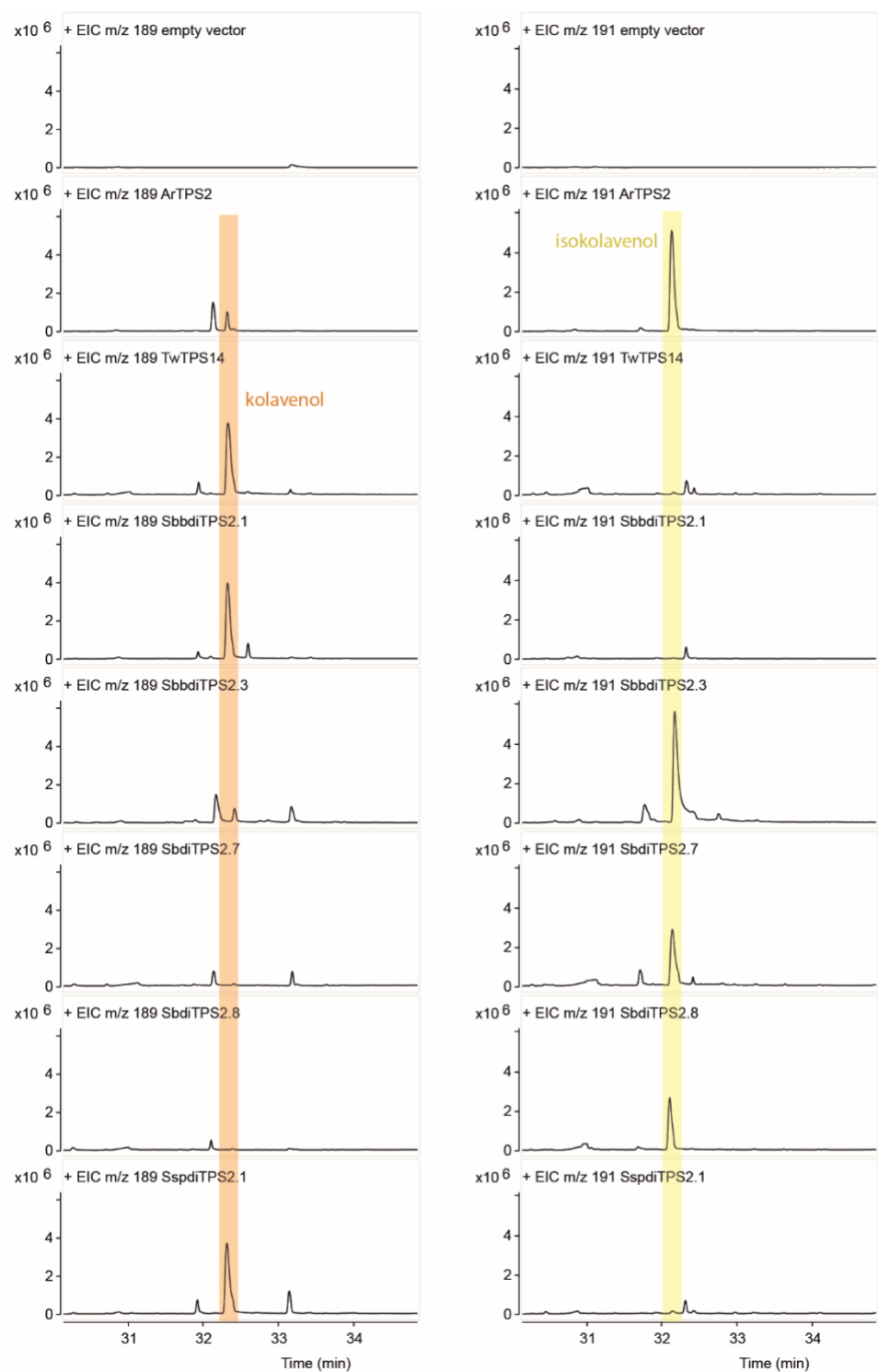

**Fig. S17. Extracted ion chromatograms (EIC) at $m/z$ 189 and at $m/z$ 191 from GC-MS analysis of extracts from *N. benthamiana* leaves expressing class II diTPSs with KPS and IKPS activities from *S. barbata*, *S. baicalensis* and *S. splendens*. The EIC at $m/z$ 189 was used for detection of kolavenol and the EIC at $m/z$ 191 was used for detection of isokolavenol. The**

orange background box indicates the presence of kolavenol and the yellow background box, the presence of isokolavenol. The enzymatic products of ArTPS2 (IKPS activity) (Johnson et al., 2019) and TwTPS14 (KPS activity) (Andersen-Ranberg *et al.*, 2016) were used as standards for isokolavenol and kolavenol, respectively.

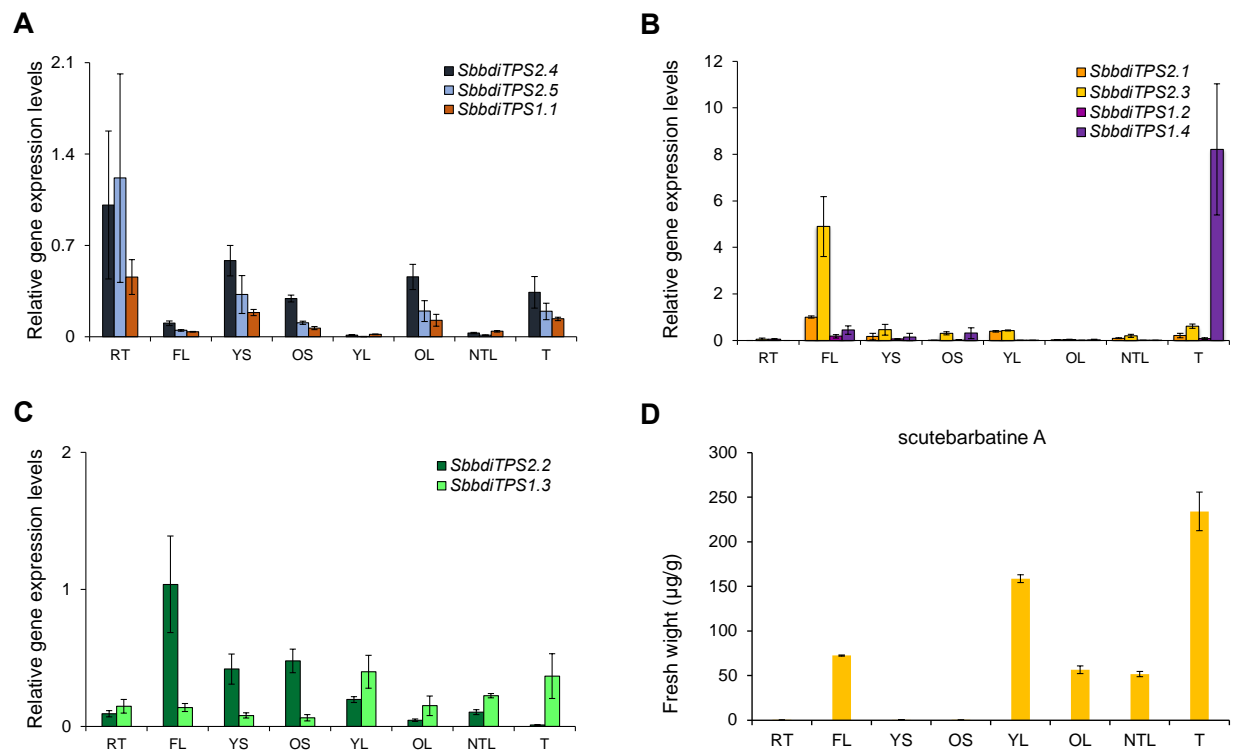

**Fig. S18. Quantitative realtime polymerase chain reaction (qRT-PCR) analysis of the expression of class II and class I diTPSs from different tissues of *Scutellaria barbata*.** RT: root; FL: flower; YS: young stem; OS: old stem; YL: young leaf (0.5 cm<leaf width<1 cm); OL: old leaf (leaf width>2 cm); NTL: non-trichome leaf (0.5 cm<leaf width<1 cm); T: trichome. A: Relative diTPS (*SbbdiTPS2.4*, *SbbdiTPS2.5* and *SbbdiTPS1.1*) gene expression levels of the miltiradiene pathway; B: Relative diTPS (*SbbdiTPS2.1*, *SbbdiTPS2.3*, *SbbdiTPS2.2* and *SbbdiTPS1.4*) gene expression levels of the clerodane pathway; C: Relative diTPS (*SbbdiTPS2.2* and *SbbdiTPS1.3*) expression levels of genes in the gibberellin pathway; D: Scutebarbatine A content (fresh weight) in different tissues of *Scutellaria barbata* from 5-month-old plants. Data were obtained from three independent biological replicates. Transcript levels were normalized to that of actin (n = 3). The colour of the bars represents different diTPS genes as highlighted in each panel. The bars represent SD.

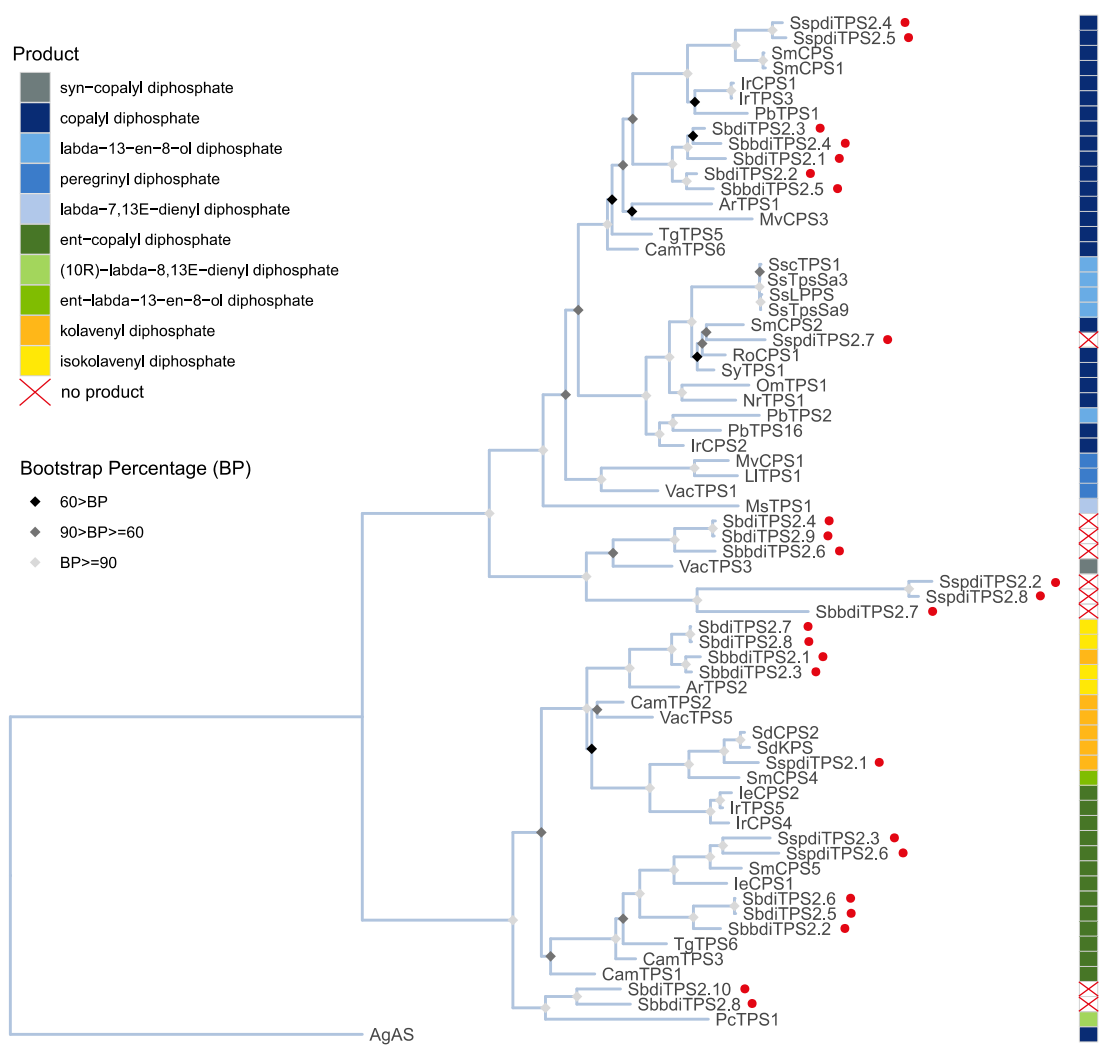

**Fig. S19. Phylogenetic tree of characterized class II diterpene synthases from Lamiaceae species.** The phylogeny was built using reported class II diTPSs from the family Lamiaceae (Table S20) and those characterized in this study are highlighted with red dots with 1000 bootstrap values. Abietadiene synthase from *Abies grandis* (AgAS) was used as the outgroup. The products produced by class II diTPS from the substrate, GGPP, are shown in boxes with different colors.

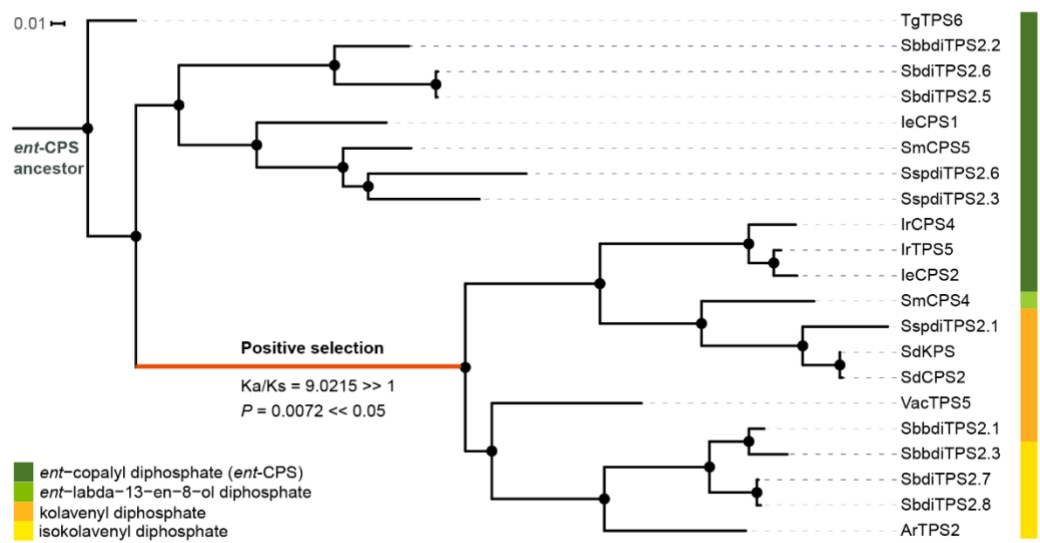

**Fig. S20. Evolution of class II clerodane synthases as result of positive selection.** Phylogram of characterized class II diTPSs with eCPS, KPS/IKPS activities from the family Lamiaceae . Coloured boxes show the *in vivo*/ *in vitro* enzyme activities of each enzyme.

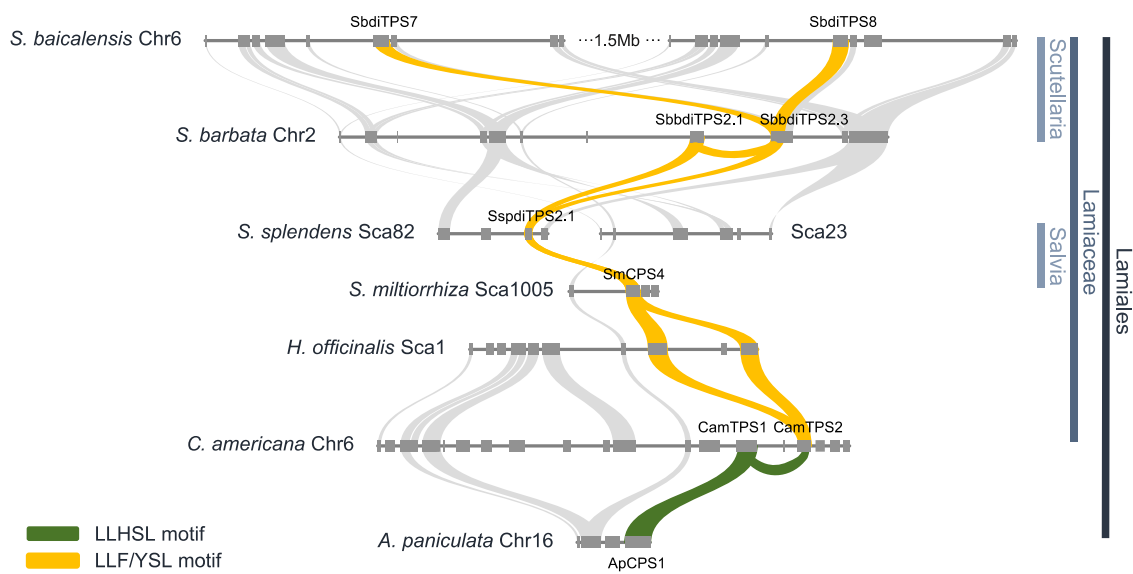

**Fig. S21. Syntenic analysis (microsynteny IV) among the species of *S. barbata*, *S. baicalensis*, *S. splendens*, *S. miltiorrhiza*, *H. officinalis*, *C. americana* (Lamiaceae family) and *A. paniculata* (Lamiales order) depicting the diTPS genes encoding eCPS, KPS and IKPS activity. In gold ribbon are highlighted the syntenic relationships between the diTPS genes with active site motif (LLFSL), while the green ribbon highlights the diterpene synthases with the active site motif (LLHSL).**

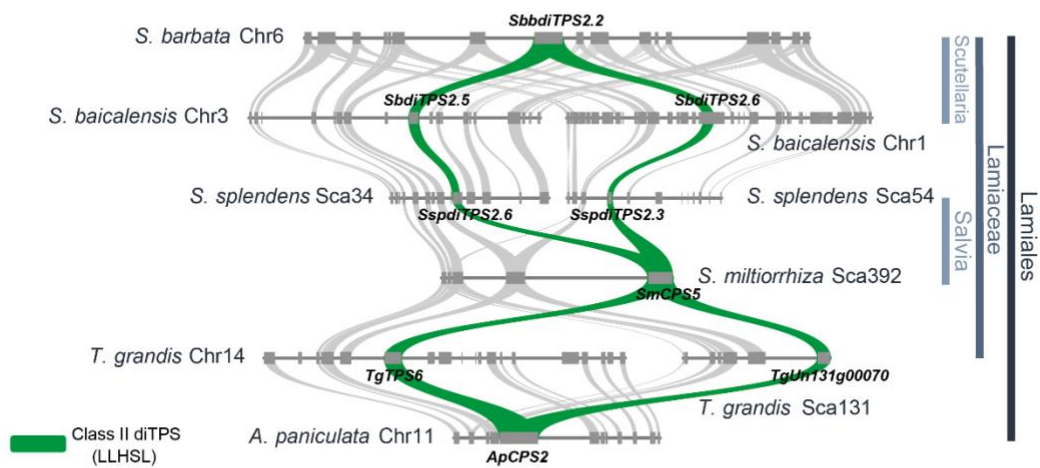

**Fig. S22. Syntenic analysis (microsynteny I) among the species of *S. barbata*, *S. baicalensis*, *S. splendens*, *S. miltiorrhiza*, *T. grandis* (Lamiaceae family) and *A. paniculata* (Lamiales order) depicting the genomic loci of diTPS genes with eCPS activity. Green colour ribbons highlight the synteny between the class II diTPSs with eCPS activity from the above-mentioned species.**

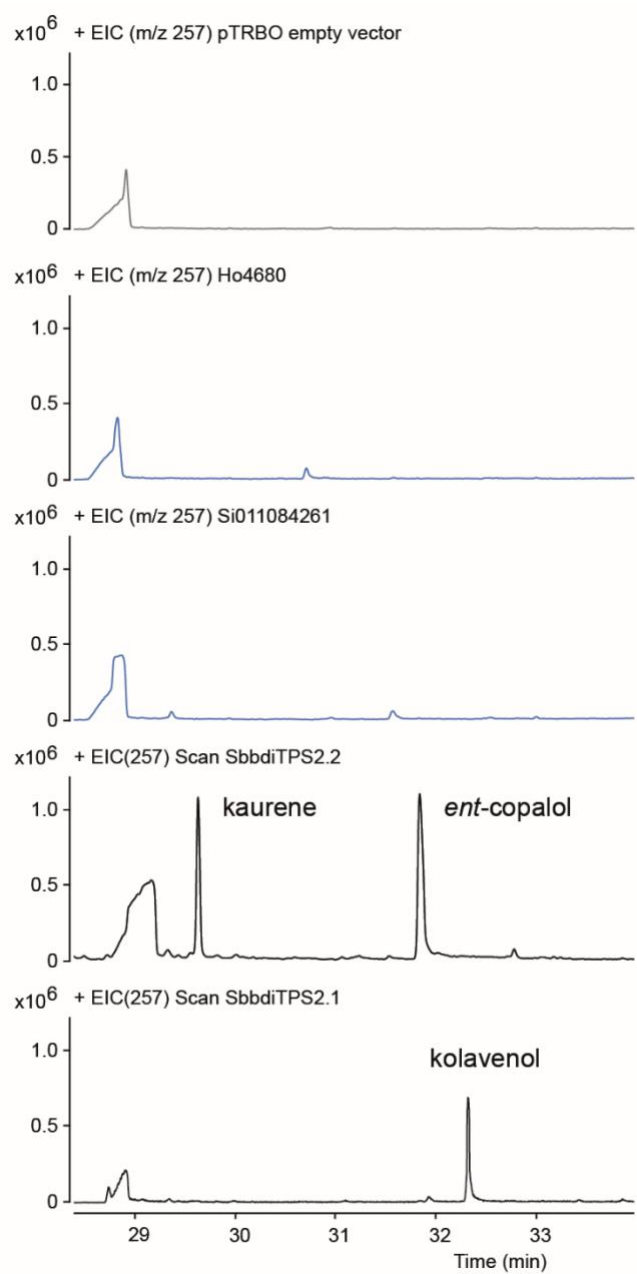

**Fig. S23. Extracted ion chromatograms (EIC) at m/z 257 from GC-MS analysis of extracts from *N. benthamiana* leaves expressing class II diTPSs Ho4680 from *H. officinalis*, and Si011084261 from *S. indicum* in comparison with SbbdiTPS2.2 (eCPS) and SbbdiTPS2.1 (KPS)**

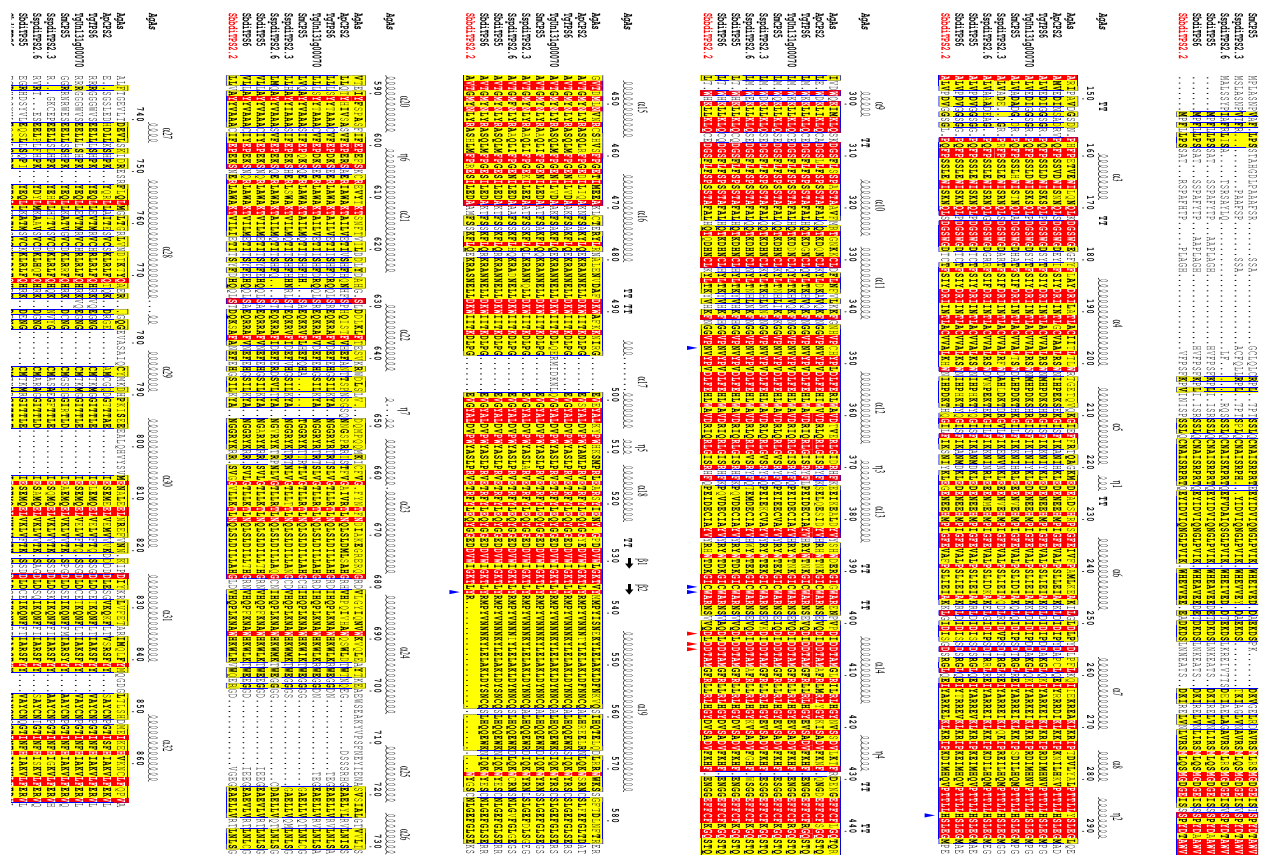

**Fig. S24. Alignment of class II diTPSs from microsynteny I (Fig. S22).** The amino acid sequences of class II diTPSs (with *eCPS* activity) were aligned with *Abies grandis* abietadiene synthase (Peters et al., 2001). Amino acid residues which are conserved within a group but not conserved from one group to the other are highlighted in yellow. Amino acid residues which are strictly conserved are highlighted in red.

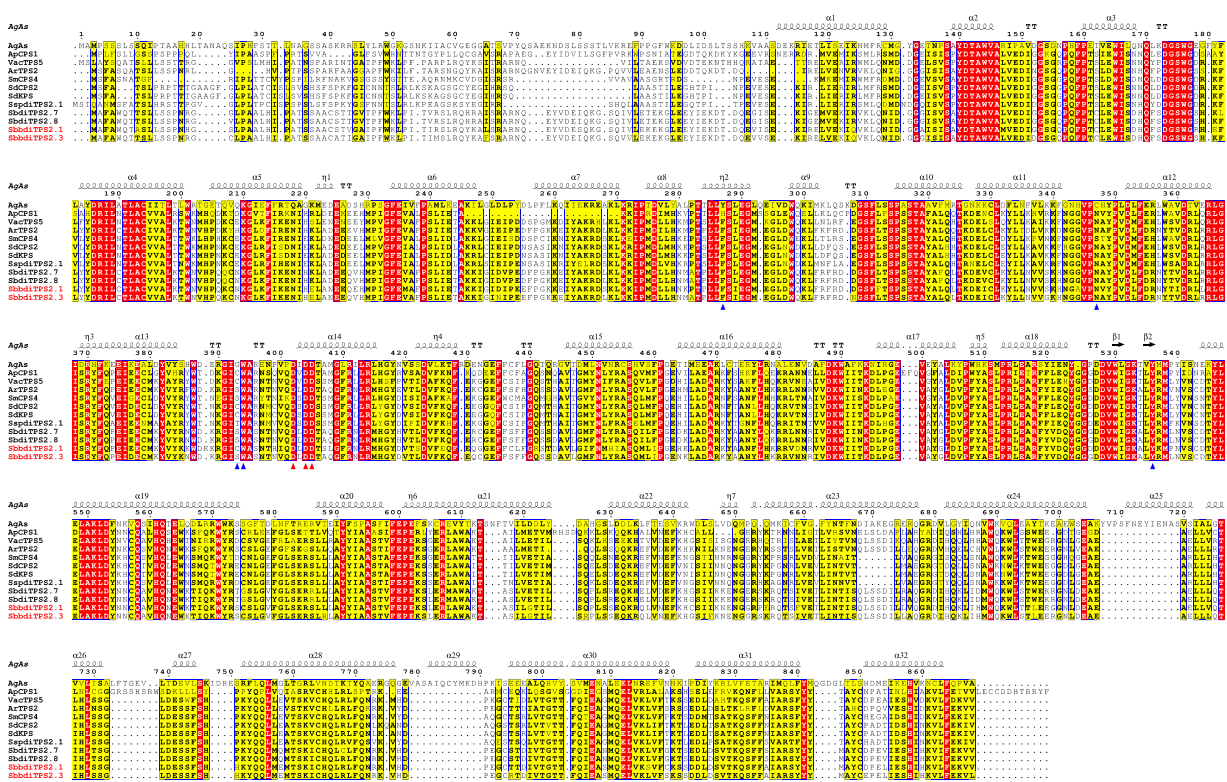

**Fig. S25. Alignment of class II diTPSs from microsynteny IV (Fig. S21).** The amino acid sequences of class II diTPSs (with KPS and IKPS activity – clerodane synthases) were aligned with *Abies grandis* abietadiene synthase (Peters *et al.*, 2001). Amino acid residues which are conserved within a group but not conserved from one group to the other are highlighted in yellow. Amino acid residues which are strictly conserved are highlighted in red.

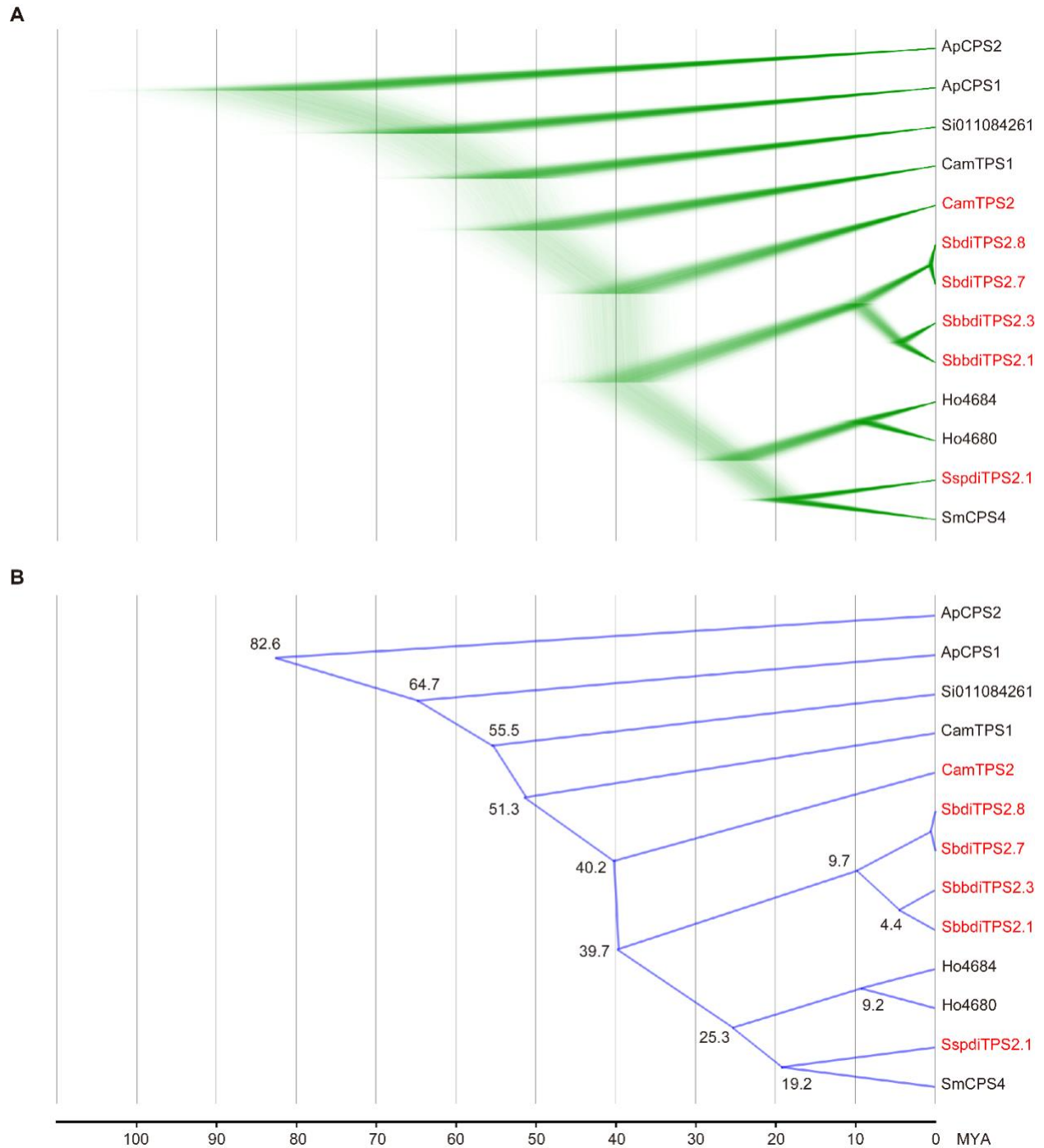

**Fig S26. Bayesian inference tree for Class II diTPSs based on the calculated species divergence time in Figure 4.** (A) shows the results from all the runs while (B) shows the consensus tree with estimated split time.

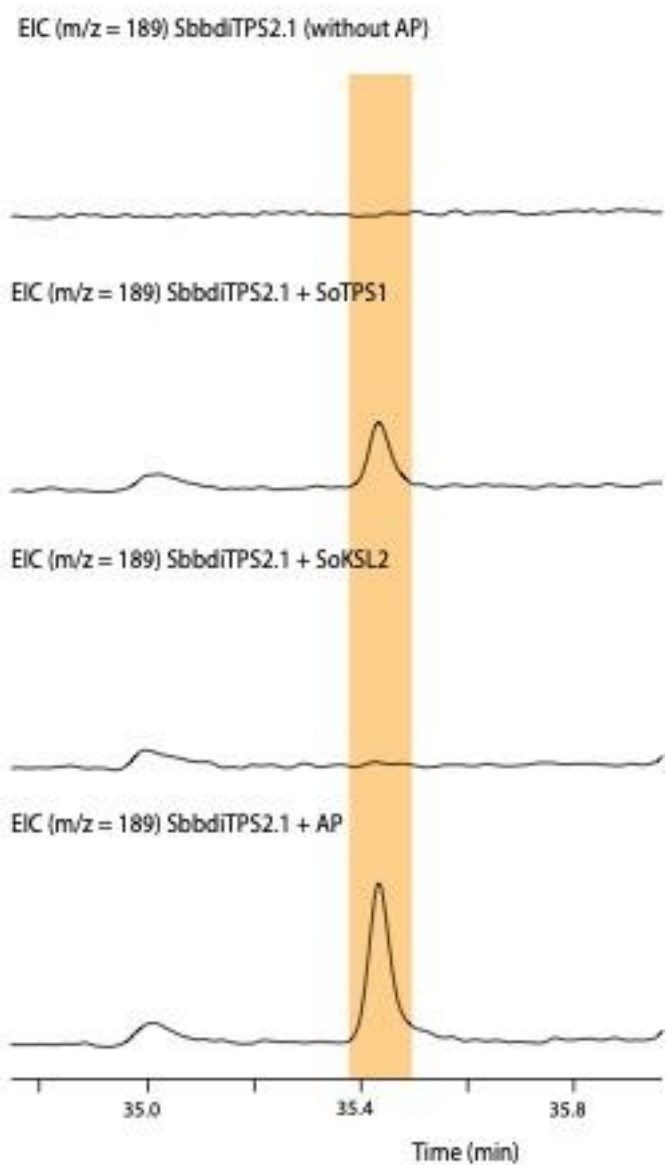

**Fig S27.** GC-MS analysis (extracted ion chromatogram $m/z$ 189) of extracts from enzyme assays expressing class II clerodane synthase SbbdiTPS2.1 in combination with class I diTPSs, SoTPS1 and SoKSL2.

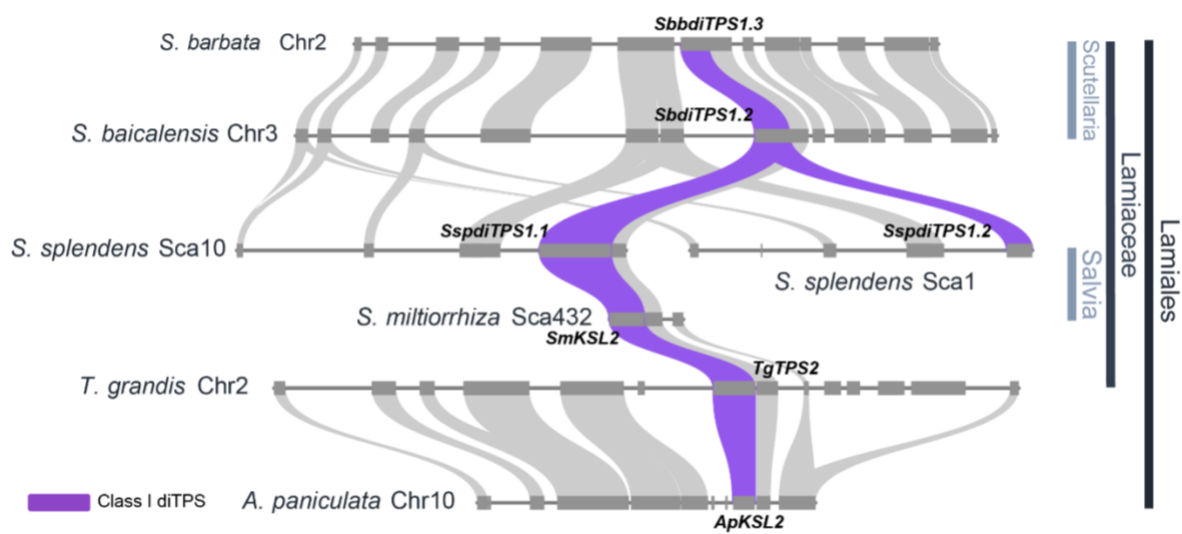

**Fig. S28. Syntenic analysis (microsynteny II) among the species of *S. barbata*, *S. baicalensis*, *S. splendens*, *S. miltiorrhiza*, *T. grandis* (Lamiaceae family) and *A. paniculata* (Lamiales order) depicting the genomic loci of diTPS genes with KS activity. The purple coloured ribbons highlight the syntenic between the class I diTPSs with KS activity from the above-mentioned species.**

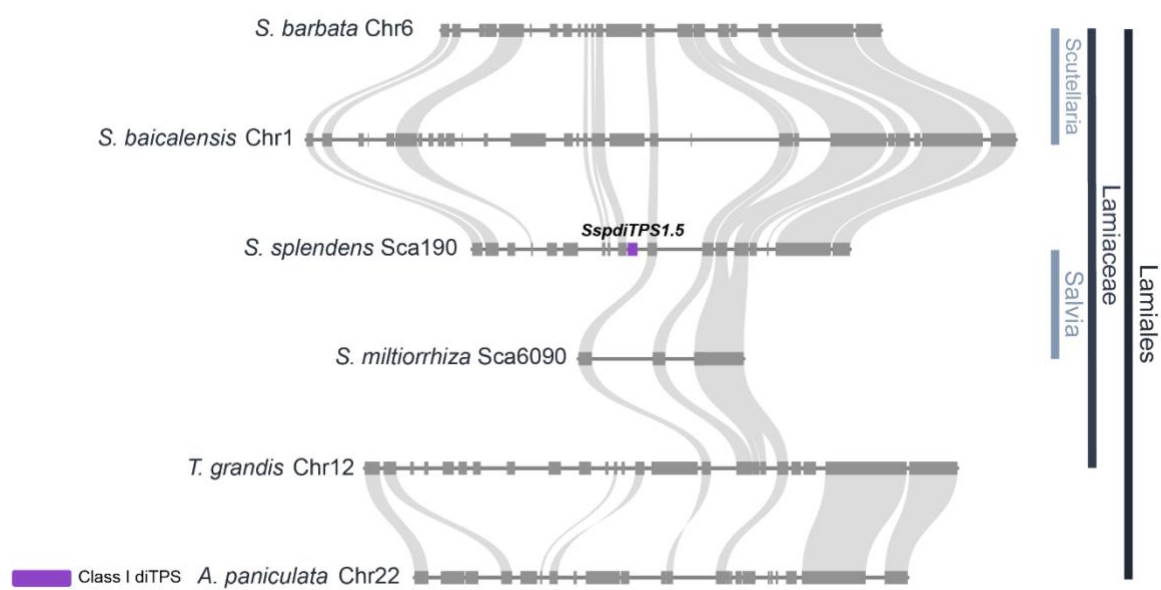

**Fig. S29.** Syntenic analysis among the species of *S. barbata*, *S. baicalensis*, *S. splendens*, *S. miltiorrhiza*, *T. grandis* (Lamiaceae family) and *A. paniculata* (Lamiales order) depicting the genomic loci of *SspdiTPS1.5* gene with KLS activity. The purple box shows the *SspdiTPS1.5* gene.

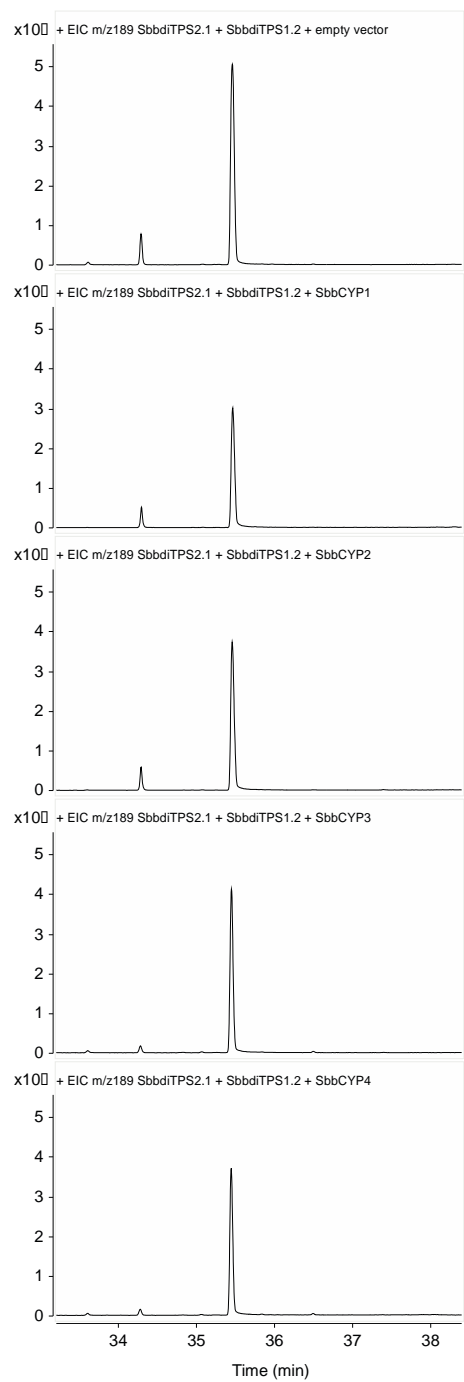

**Fig S30. GC-MS analysis (extracted ion chromatogram m/z 189) of extracts from yeast strains expressing GGPPS (Erg20(F96C)), KPS, KLS and cytochrome P450s from the *Scutellaria barbata* genomic region including the ferruginol biosynthetic gene cluster (Figure 5).**

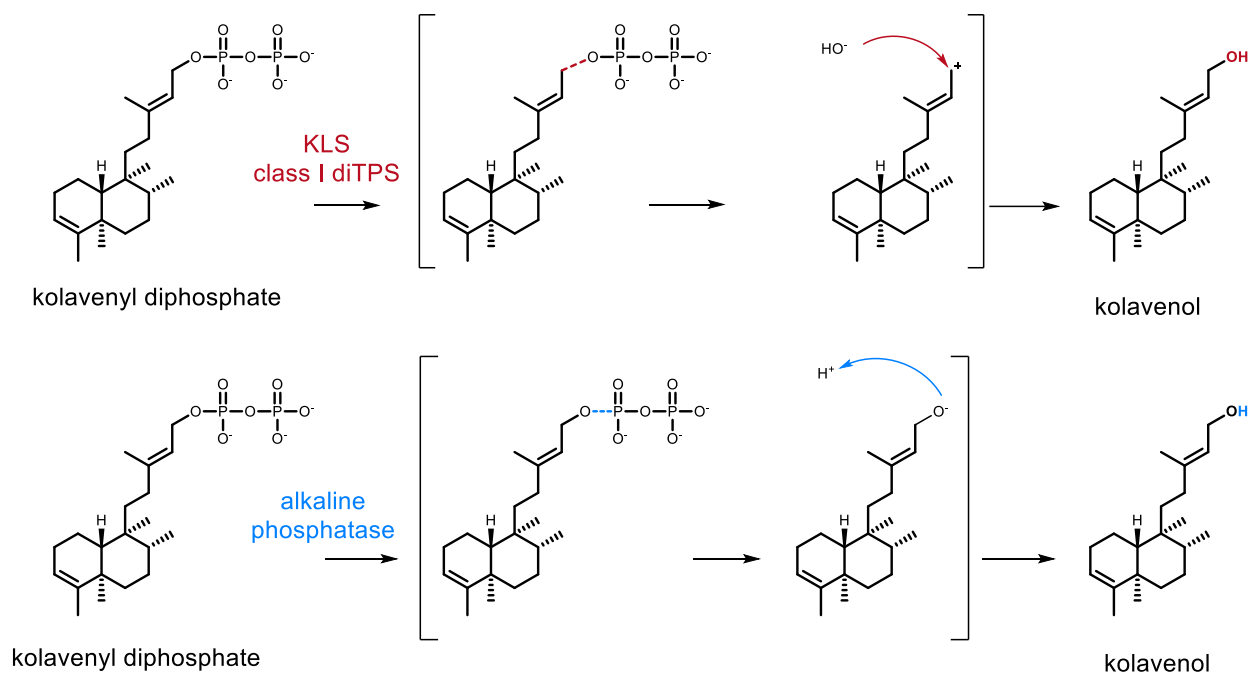

**Fig. S31. Differences in catalytic mechanisms between kolavenol synthase (KLS – class I diTPS) and alkaline phosphatase (AP) for the formation of kolavenol from kolavenyl diphosphate.** KLS as class I diTPS (lyase enzyme) catalyzes the breaking of the bond between carbon-oxygen atoms resulting in the formation of a carbocation and the later attack of an OH group for formation of kolavenol. In contrast, AP catalyzes the hydrolysis of the bond between the oxygen-phosphorus atoms resulting to formation of anion.

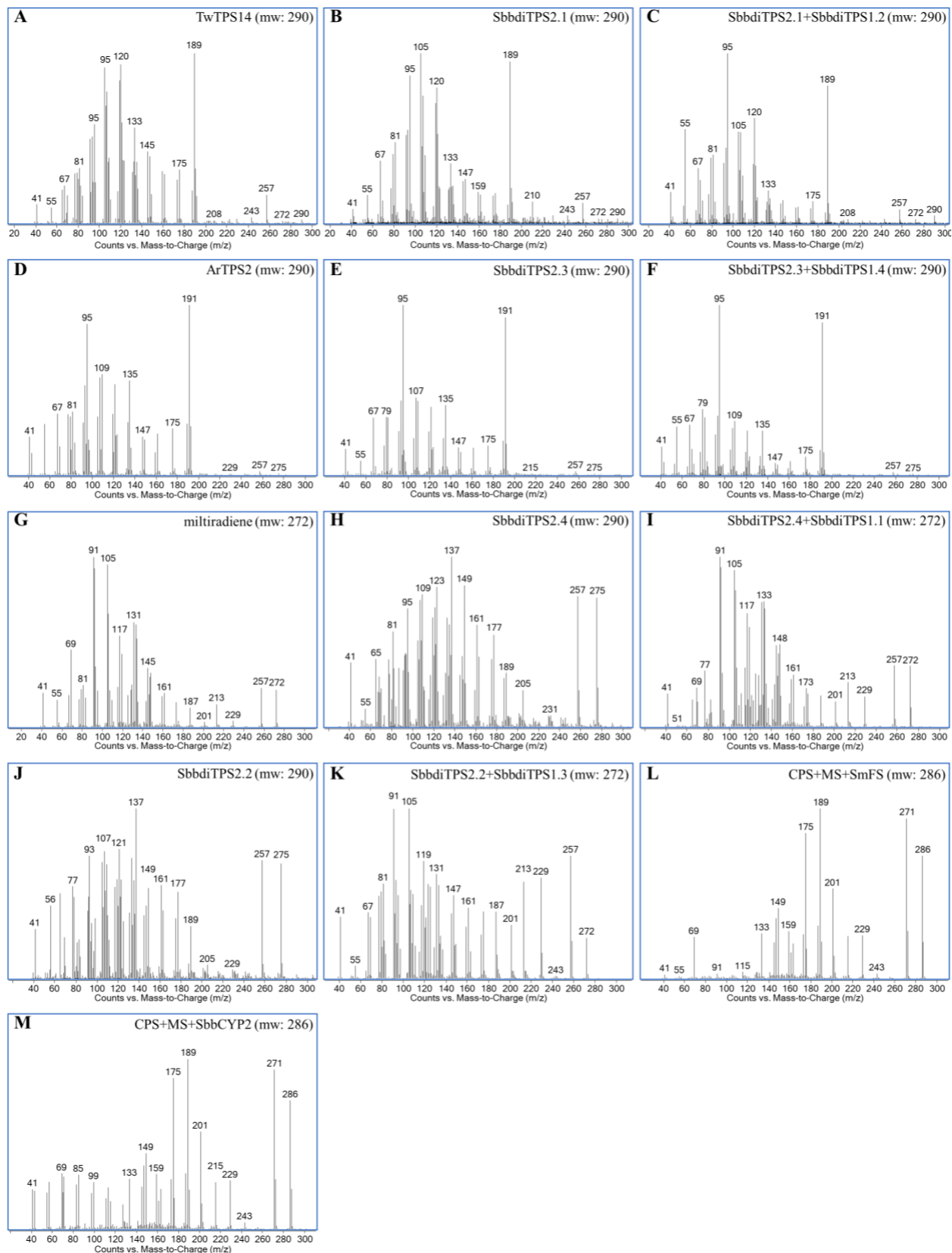

**Fig. S32. EI MS spectra from GCMS of major diterpene scaffolds as derived by combination of class II and class I diterpene synthases *in vivo* and *in vitro*.** The EI MS spectra of kolavenol as extracted from GCMS analysis of expression of (a) TwTPS14 in *N. benthamiana* (Andersen-Ranberg *et al.*, 2016), (b) SbbdiTPS2.1 in *N. benthamiana*, (c) SbbdiTPS2.1+SbbdiTPS1.2 in *E. coli*; EI MS spectra of isokolavenol as extracted from GCMS analysis of expression of (d) ArTPS2 in *N. benthamiana* (Johnson *et al.*, 2019), (e) SbbdiTPS2.3 in *N. benthamiana*, (f) SbbdiTPS2.3+SbbdiTPS1.4 in *E. coli*; EI MS spectra of miltiradiene as extracted from GCMS analysis of (g) isolated miltiradiene, (i) expression of SbbdiTPS2.4 + SbbdiTPS1.1 in *N. benthamiana*; EI MS spectra of copalol as extracted from GCMS analysis of (h) expression of SbbdiTPS2.4 in *N. benthamiana*; EI MS spectra of *ent*-copalol as extracted from GCMS analysis of (j) expression of SbbdiTPS2.2 in *N. benthamiana*; EI MS spectra of *ent*-kaurene as extracted from GCMS analysis of (k) expression of SbbdiTPS2.2+ SbbdiTPS1.3 in *N. benthamiana*; EI MS spectra of ferruginol as extracted from GCMS analysis of (l) expression of CPS+ MS+SmCYP76AH1 in yeast (*S. cerevisiae*), (l) expression of CPS+ MS+SbbCYP2 in yeast (*S. cerevisiae*).

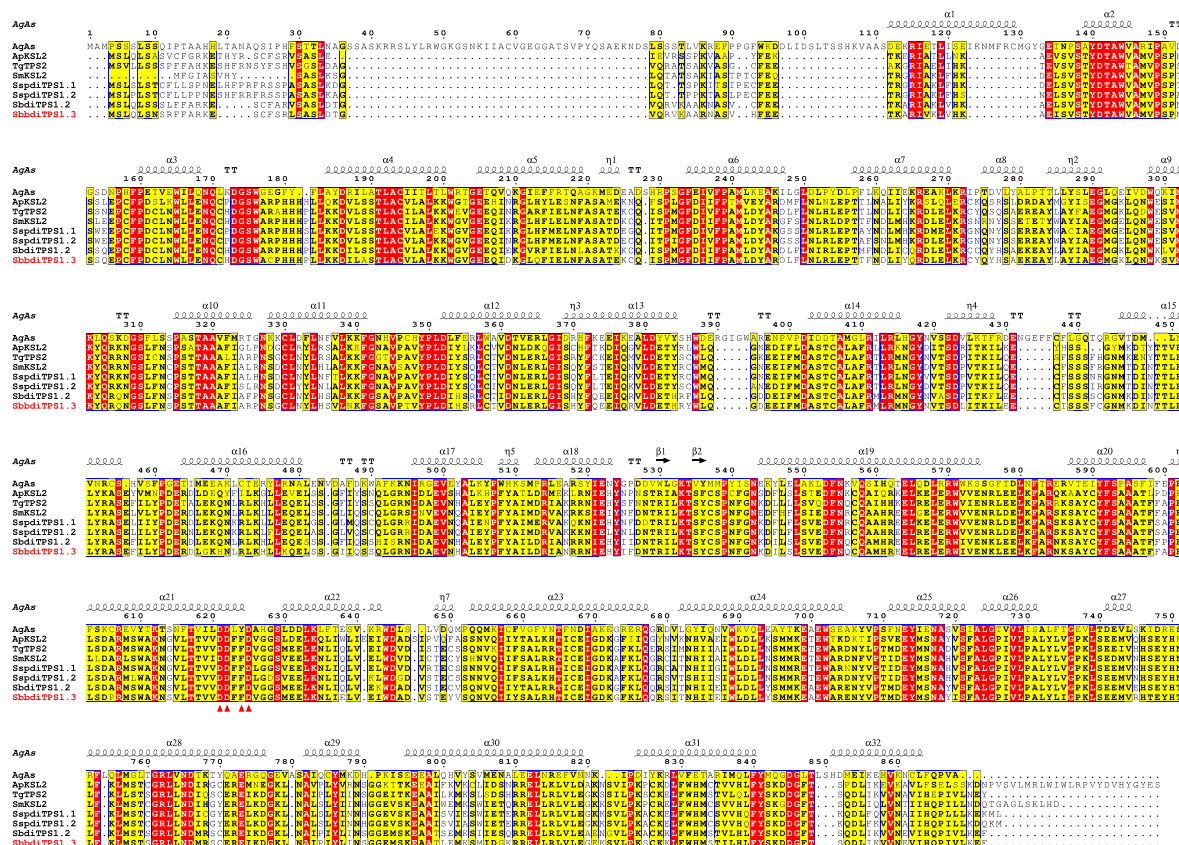

**Fig. S33. Alignment of class I diTPSs from microsynteny II (Fig. S17).** The amino acid sequences of class I diTPSs (with KS activity) were aligned with *Abies grandis* abietadiene synthase (Peters *et al.*, 2001). Amino acid residues which are conserved within a group but not conserved from one group to the other are highlighted in yellow. Amino acid residues which are strictly conserved are highlighted in red.

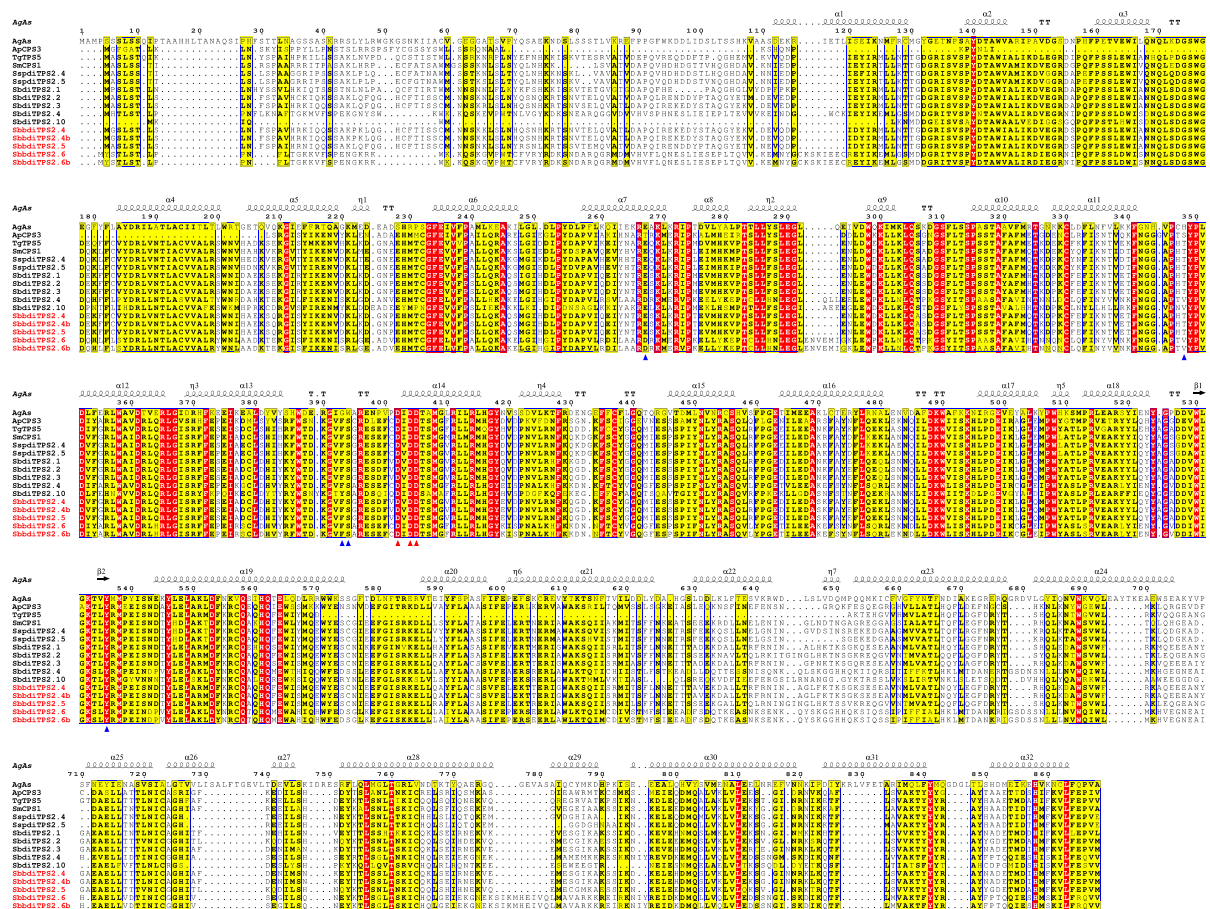

**Fig. S34. Alignment of class II diTPSs from microsynteny III (Figure 4).** The amino acid sequences of class II diTPSs (with CPS activity) were aligned with *Abies grandis* abietadiene synthase (Peters *et al.*, 2001). Amino acid residues which are conserved within a group but not conserved from one group to the other are highlighted in yellow. Amino acid residues which are strictly conserved are highlighted in red.

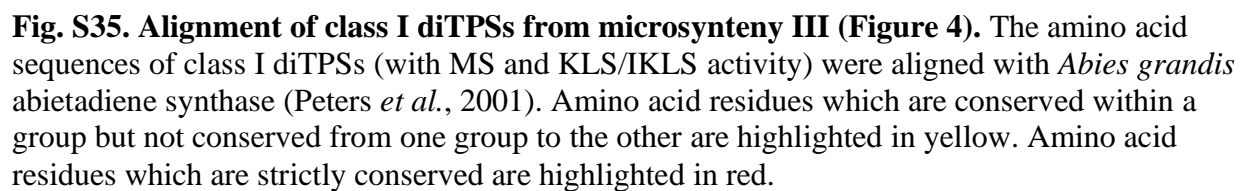

**Table S1. List of clerodane diterpenoids isolated from *Scutellaria barbata* and their reported cytotoxic activities. See the attached excel file**

**Table S2. Sequencing methods and their depth of coverage.**

| Lamiaceae | Subfamily | Genus |
| --- | --- | --- |
|  | Nepetoideae | Nepeta (Hussain et al., 2009), Salvia (Brieskorn and Stehle, 1973), Plectranthus (Ri al., 2002), Coleus (Ávila et al., 2016), Elshotzia (Hu et al., 2008) |
|  | Lamiodeae | Leonurus (Liu et al., 2014), Phlomis (Hussain et al., 2011), Ballota (Ahmad et al., 2004), Gomphostemma (Zhang et al., 2009), Otostegia (Ahmad et al., 2005) |
|  | Ajugoideae | Ajuga (SHIMOMURA et al., 1989), Glossocarya (Rasikari et al., 2005), Teucrium (Cuadrado et al., 1991), Kinostemon (Ye et al., 2002), Clerodendrum (Harada and U 1978), Caryopteris (Harada and Uda, 1978) |
|  | Peronematoideae | Peronema (Ahmad *et al.*, 2005) |
|  | Scutellariodeae | Scutellaria (Kikuchi et al., 1987), Tinnea (Borges et al., 2013) |
|  | Premnoideae | Premna (Chin et al., 2006), Cornutia (Chen et al., 1992) |
|  | Viticoideae | Vitex (Rasyid et al., 2017) |
|  | Symphorematoideae* |  |
|  | Tectonoideae | Tectona (Macías et al., 2010) |
|  | Prostantheroideae | Stachys (Fazio et al., 1992), Pityrodia (Castro and Coll, 2008), Cyanostegia (Jefferi al., 1973) |
|  | Callicarpoideae | Callicarpa (Yuan et al., 2019) |

\*There are no phytochemical data for any plant from the three genera (Congea, Sphenodesme, Symphorema) that consist of subfamily Symphorematoideae.

**Table S3. Sequencing methods and their depth of coverage.**

| Pair-end libraries | Total data(Gb) | Sequence coverage(X) |
| --- | --- | --- |
| Illumina Novaseq | 65.26 | 157.29 |
| PacBio | 135.29 | 326.09 |
| Bionano | 265.2 | 639.21 |
| Illumina Novaseq<br>(for Hi-C) | 68.93 | 166.14 |

**Table S4.**

(A) Parameters of first version reference genome assembly.

| Sample_ID | length |  | number |  |
| --- | --- | --- | --- | --- |
|  | Contig | Scaffold | Contig* | Scaffold |
| Total | 376,212,188 | **376,956,065** | 396 | 324 |
| Max | 20,085,936 | 32,118,005 | - | - |
| Number≥2000 | - | - | 396 | 324 |
| N50 | 7,463,337 | **14,970,362** | 18 | 9 |
| N60 | 6,446,489 | 13,637,937 | 23 | 12 |
| N70 | 4,376,436 | 11,827,991 | 30 | 15 |
| N80 | 2,565,505 | 9,596,772 | 41 | 19 |
| N90 | 979,745 | 7,310,276 | 63 | 23 |

\* Contig after scaffolding

(B) Assembly improvement using Hi-C

|  | Input Assembly | LACHESIS assembly |
| --- | --- | --- |
| **Total Length** | 376.96Mb | 376.97 Mb |
| **N50** | 14.97 Mb | 26.00Mb |
| **N90** | 7.31 Mb | 18.27 Mb |
| **Longest Scaffolds** | 32.12 Mb | 44.71 Mb |
| **Number of Scaffolds** | 324 | 356 |
| **Contig N50** | 7.46Mb | 7.24Mb |

**Table S5. Summary of final *S. barbata* genome assembly.**

| Assembly feature | Statistics |
| --- | --- |
| Estimated genome size (k-mer analysis) | 414.98 Mb |
| Assembly length | 376.96 Mb |
| Length loaded on pseudochromosomes | 343.99Mb |
| Number of pseudochromosomes | 13 |
| Number of super-scaffolds | 356 |
| N50 of super-scaffolds | 26.00 Mb |
| Longest super-scaffolds | 44.71 Mb |
| Assembly % of genome | 90.84% |
| Repeat region of % assembly | 54.50% |
| Predicted gene models | 25899 |
| Average coding sequence length | 1198.90 bp |
| Average exons per gene | 5.14 |

**Table S6. Scaffold number and length grouped on pseudochromosomes**

| **Group** | **Number of scaffold** | **Total Length(bp)** |
| --- | --- | --- |
| **Hic_asm_0** | 14 | 44,707,576 |
| **Hic_asm_1** | 8 | 37,561,464 |
| **Hic_asm_2** | 3 | 33,675,200 |
| **Hic_asm_3** | 12 | 32,608,499 |
| **Hic_asm_4** | 6 | 28,390,736 |
| **Hic_asm_5** | 18 | 26,007,137 |
| **Hic_asm_6** | 13 | 22,547,331 |
| **Hic_asm_7** | 12 | 21,931,902 |
| **Hic_asm_8** | 9 | 20,892,611 |
| **Hic_asm_9** | 3 | 19,619,875 |
| **Hic_asm_10** | 4 | 19,126,067 |
| **Hic_asm_11** | 3 | 18,645,315 |
| **Hic_asm_12** | 5 | 18,272,457 |
| **TOTAL** | 110 | **343,986,170 (91.25%)** |

**Table S7. Contents of each nucleotide in the draft *S. barbata* genome sequence.**

|  | Number (bp) | % of genome |
| --- | --- | --- |
| A | 124,554,272 | 33.04% |
| T | 124,364,737 | 32.99% |
| C | 63,654,526 | 16.89% |
| G | 63,638,653 | 16.88% |
| N | 743,877 | 0.20% |
| Total | 376,956,065 | - |
| GC* | 127,293,179 | 33.84% |

\*GC content of the genome without N's

**Table S8. SNP calling of the genome of *S. barbata***

|  | Number | %Percentage |
| --- | --- | --- |
| All SNP | 4,680 | 0.0132 |
| Heterozygous SNP | 4,280 | 0.0121 |
| Homologous SNP | 400 | 0.0011 |

**Table S9. Evaluation of coverage by mapping short reads**

|  |  | Percentage |
| --- | --- | --- |
| Reads | Mapping rate (%) | 96.50 |
| Genome | Average sequencing depth | 127.84 |
|  | Coverage (%) | 99.78 |
|  | Coverage at least 4X (%) | 99.71 |
|  | Coverage at least 10X (%) | 99.62 |
|  | Coverage at least 20X (%) | 99.47 |

**Table S10. Evaluation of coverage by mapping ESTs**

| Dataset | Number | Total length<br>(bp) | Sequences<br>Covered by<br>Assembly<br>(%) * | Sequences with<br>>90% coverage in<br>one scaffold ** |  | Sequences with >50%<br>coverage in one<br>scaffold |  |
| --- | --- | --- | --- | --- | --- | --- | --- |
|  |  |  |  | Number | Percent<br>(%) | Number | Percent<br>(%) |
| >0bp | 72106 | 122185731 | 99.885 | 66911 | 92.795 | 71604 | 99.304 |
| >200bp | 72106 | 122185731 | 99.885 | 66911 | 92.795 | 71604 | 99.304 |
| >500bp | 61529 | 118368282 | 99.924 | 56985 | 92.615 | 61119 | 99.334 |
| >1k | 45891 | 106939687 | 99.985 | 42321 | 92.221 | 45636 | 99.444 |
| >2k | 22965 | 73109074 | 99.996 | 20633 | 89.845 | 22812 | 99.334 |

**Table S11. CEGMA evaluation of the assembly**

| species | complete |  | Complete+partial |  |
| --- | --- | --- | --- | --- |
|  | #Core genes | % completeness | #Prots | % completeness |
| *S. barbata* | 223 | 93.95 | 237 | 95.56 |

**Table S12. BUSCO evaluation of the assembly.** C: Complete Single-Copy BUSCOs; D: Complete Duplicated BUSCOs; F: Fragmented BUSCOs; M: Missing BUSCOs; n: Total BUSCO groups searched.:

| Species | BUSCO notation assessment results |
| --- | --- |
| *S. barbata* | C: 93.1% [S: 86.3%, D: 6.8%], F: 1.7%, M: 5.2%, n: 1440 |

**Table S13. Metrics for gene annotation**

| Species | Number | Average transcript length (bp) | Average CDS length (bp) | Average exons per gene | Average exon length (bp) | Average intron length (bp) |
| --- | --- | --- | --- | --- | --- | --- |
| *S. barbata* | 25,899 | 3,065.18 | 1,198.90 | 5.14 | 233.12 | 450.49 |

**Table S14. Metrics for interspersed sequences**

|  | *De novo* + Rep base |  | TE Proteins |  | Combined TEs |  |
| --- | --- | --- | --- | --- | --- | --- |
|  | Length (bp) | % in Genome | Length (bp) | % in Genome | Length (bp) | % in Genome |
| DNA | 13,057,209 | 3.46 | 635,348 | 0.17 | 13,311,687 | **3.53** |
| LINE | 876,570 | 0.23 | 456,560 | 0.12 | 1,184,669 | 0.31 |
| SINE | 54,221 | 0.01 | 0 | 0.00 | 54,221 | 0.01 |
| LTR | 163,763,603 | 43.44 | 32,107,234 | 8.52 | 165,126,155 | **43.80** |
| Unknown | 25,653,934 | 6.81 | 0 | 0.00 | 25,653,934 | 6.81 |
| Total | 198,745,169 | 52.72 | 33,198,832 | 8.81 | 200,008,480 | **53.06** |

**Table S15. Metrics for non-coding RNA**

| Type | Copy (w*) | Average length (bp) | Total length (bp) | % of genome |
| --- | --- | --- | --- | --- |
| miRNA | **623** | 114.67 | 71,437 | 0.018951 |
| tRNA | **566** | 74.83 | 42,353 | 0.011235 |
| rRNA | **1,599** | 237.80 | 380,244 | 0.10 |
| snRNA | **513** | 114.40 | 58,688 | 0.015569 |

**Table S16. List of clerodane diterpenoids isolated from *Salvia splendens*.**

**Table S17. List of diterpene synthases from Lamiaceae family used for construction of class II diTPS (Fig. S21) and class I diTPS (Fig. S22) phylogenetic trees.**

|  | Species | Accession number or DOI |
| --- | --- | --- |
| **Class II diTPS** |  |  |
| SsTpsSa3 | *Salvia sclarea* | AET21247 |
| SsTpsSa9 | *Salvia sclarea* | AET21248 |
| PbTPS2 | *Plectranthus barbatus* | KF444507 |
| SmCPS4 | *Salvia miltiorrhiza f. alba* | KP063138 |
| ArTPS1 | *Ajuga reptans* | AZB50377 |
| IrTPS3 | *Isodon rubescens* | KX831651 |
| MvCPS3 | *Marrubium vulgare* | KJ584452 |
| NrTPS1 | *Nepeta racemosa* | AZB50382 |
| OmTPS1 | *Origanum majorana* | AZB50383 |
| PbTPS1 | *Plectranthus barbatus* | KF444506 |
| PbTPS16 | *Plectranthus barbatus* | AZB50379 |
| RoCPS1 | *Rosmarinus officinalis* | KF805857 |
| SfCPS | *Salvia fruticosa* | AJQ30184 |
| SmCPS | *Salvia miltiorrhiza* | ABV57835 |
| SyTPS1 | *Salvia yangii* | AZB50384 |
| TgTPS5 | *Tectona grandis* | 10.1093/gigascience/giz005 |
| IrCPS1 | *Isodon rubescens* | APJ36371 |
| SmCPS1 | *Salvia miltiorrhiza f. alba* | KC814639 |
| IrCPS2 | *Isodon rubescens* | APJ36372 |
| SmCPS2 | *Salvia miltiorrhiza f. alba* | KC814640 |
| SscTPS1 | *Salvia sclarea* | ADW66454 |
| PcTPS1 | *Pogostemon cablin* | AZB50385 |
| IeCPS2 | *Isodon eriocalyx* | AEP03175 |
| IrCPS5 | *Isodon rubescens* | KU180503 |
| IrTPS5 | *Isodon rubescens* | ARO38143 |
| SdCPS1 | *Salvia divinorum* | KX424876 |
| TgTPS6 | *Tectona grandis* | 10.1093/gigascience/giz005 |
| IrCPS4 | *Isodon rubescens* | APJ36374 |
| SmCPS5 | *Salvia miltiorrhiza f. alba* | KC814642 |
| SdCPS2 | *Salvia divinorum* | KX424877 |
| SdKPS | *Salvia divinorum* | AOZ15895 |
| VacTPS5 | *Vitex agnus-castus* | MG696752 |
| SsLPPS | *Salvia sclarea* | AFU61897 |
| MsTPS1 | *Mesosphaerum suaveolens* | MH626627 |
| IeCPS1 | *Isodon eriocalyx* | AEP03177 |
| ArTPS2 | *Ajuga reptans* | MH626625 |
| LlTPS1 | *Leonotis leonurus* | AZB50381 |
| MvCPS1 | *Marrubium vulgare* | AIE77090 |
| VacTPS1 | *Vitex agnus-castus* | MG696748 |
| VacTPS3 | *Vitex agnus-castus* | MG696750 |
| CamTPS1 | *Callicarpa americana* | QMW69081 |
| CamTPS2 | *Callicarpa americana* | QMW69082 |
| CamTPS3 | *Callicarpa americana* | QMW69080 |
| CamTPS6 | *Callicarpa americana* | QMW69083 |
| SspdiTPS2.1 | *Salvia splendens* | Saspl_043012 |
| SspdiTPS2.2 | *Salvia splendens* | Saspl_048790 |
| SspdiTPS2.3 | *Salvia splendens* | Saspl_027494 |
| SspdiTPS2.4 | *Salvia splendens* | Saspl_009166 |
| SspdiTPS2.5 | *Salvia splendens* | Saspl_017770 |
| SspdiTPS2.6 | *Salvia splendens* | Saspl_009980 |
| SspdiTPS2.7 | *Salvia splendens* | Saspl_013955 |
| SspdiTPS2.8 | *Salvia splendens* | Saspl_006526 |
| SbdiTPS2.1 | *Scutellaria barbata* | Sb06g19680 |
| SbdiTPS2.2 | *Scutellaria barbata* | Sb06g19670 |
| SbdiTPS2.3 | *Scutellaria barbata* | Sb06g19660 |
| SbdiTPS2.4 | *Scutellaria barbata* | Sb06g19690 |
| SbdiTPS2.5 | *Scutellaria barbata* | Sb03g27890 |
| SbdiTPS2.6 | *Scutellaria barbata* | Sb01g04220 |
| SbdiTPS2.7 | *Scutellaria barbata* | Sb06g03970 |
| SbdiTPS2.8 | *Scutellaria barbata* | Sb06g05650 |
| SbdiTPS2.9 | *Scutellaria barbata* | Sb08g08190 |
| SbdiTPS2.10 | *Scutellaria barbata* | Sb06g19710 |
| SbbdiTPS2.1 | *Scutellaria baicalensis* | Sbb02g0030951 |
| SbbdiTPS2.2 | *Scutellaria baicalensis* | Sbb06g0014700 |
| SbbdiTPS2.3 | *Scutellaria baicalensis* | Sbb02g0030960 |
| SbbdiTPS2.4 | *Scutellaria baicalensis* | Sbb02g0014540 |
| SbbdiTPS2.5 | *Scutellaria baicalensis* | Sbb02g0014580 |
| SbbdiTPS2.6 | *Scutellaria baicalensis* | Sbb02g0014541 |
| SbbdiTPS2.7 | *Scutellaria baicalensis* | Sbb02g0014610 |
| SbbdiTPS2.8 | *Scutellaria baicalensis* | Sbb02g0014630 |
| **Class I diTPS** |  |  |
| MvELS | *Marrubium vulgare* | KJ584454 |
| MsTPS1 | *Mentha spicata* | MH626616 |
| VacTPS2 | *Vitex agnus-castus* | MG696749 |
| VacTPS6 | *Vitex agnus-castus* | MG696753 |
| PbTPS4 | *Plectranthus barbatus* | KF444509 |
| IrKSL4 | *Isodon rubescens* | KX580633 |
| MvEKS | *Marrubium vulgare* | KJ584453 |
| NmTPS2 | *Nepeta mussinii* | MH626617 |
| PbTPS14 | *Plectranthus barbatus* | AGN70881 |
| SmKSL2 | *Salvia miltiorrhiza f. alba* | KC814643 |
| TgTPS2 | *Tectona grandis* | 10.1042/BJ20120654 |
| VacTPS4 | *Vitex agnus-castus* | MG696751 |
| IrKSL5 | *Isodon rubescens* | KX580634 |
| IrKSL2 | *Isodon rubescens* | KU180505 |
| IrKSL6 | *Isodon rubescens* | KX580635 |
| SdKSL1 | *Salvia divinorum* | KY057342 |
| OmTPS4 | *Origanum majorana* | MH626619 |
| PbTPS3 | *Plectranthus barbatus* | KF444508 |
| ArTPS3 | *Ajuga reptans* | MH626614 |
| IrKSL1 | *Isodon rubescens* | KU180504 |
| IrKSL3 | *Isodon rubescens* | KU180506 |
| IrTPS4 | *Isodon rubescens* | KX831652 |
| PaTPS3 | *Perovskia atriplicifolia* | MH626621 |
| PvTPS1 | *Prunella vulgaris* | MH626622 |
| RoKSL1 | *Salvia rosmarinus* | KF805858 |
| RoKSL2 | *Salvia rosmarinus* | KF805859 |
| SfKSL | *Salvia fruticosa* | KP091841 |
| SmKSL1 | *Salvia miltiorrhiza* | ABV08817 |
| SoTPS1 | *Salvia officinalis* | MH626623 |
| SpMilS | *Salvia pomifera* | KP119676 |
| TgTPS1 | *Tectona grandis* | AWW87326 |
| IrTPS2 | *Isodon rubescens* | KX831650 |
| OmTPS5 | *Origanum majorana* | MH626620 |
| LlTPS4 | *Leonotis leonurus* | MH626615 |
| SsSS | *Salvia sclarea* | AET21246 |
| SsTps1132 | *Salvia sclarea* | JN133922 |
| OmTPS3 | *Origanum majorana* | MH626618 |
| SspdiTPS1.1 | *Salvia splendens* | Saspl_006722 |
| SspdiTPS1.2 | *Salvia splendens* | Saspl_001373 |
| SspdiTPS1.3 | *Salvia splendens* | Saspl_037723 |
| SspdiTPS1.4 | *Salvia splendens* | Saspl_048791 |
| SspdiTPS1.5 | *Salvia splendens* | Saspl_037211 |
| SbbdiTPS1.1 | *Scutellaria barbata* | Sbb02g0014530 |
| SbbdiTPS1.2 | *Scutellaria barbata* | Sbb02g0014640 |
| SbbdiTPS1.3 | *Scutellaria barbata* | Sbb02g0011460 |
| SbbdiTPS1.4 | *Scutellaria barbata* | Sbb02g0014620 |
| SbdiTPS1.1 | *Scutellaria baicalensis* | Sb06g19650 |
| SbdiTPS1.2 | *Scutellaria baicalensis* | Sb03g16790 |
| SbdiTPS1.3 | *Scutellaria baicalensis* | Sb06g19700 |

**Table S18. Linearized plasmids with restriction enzyme sites and corresponding overhangs.**

| Plasmid | Restriction enzymes | Overhangs |
| --- | --- | --- |
| pESC-HIS (Gal 1) | *Bam*H I-HF/*Sal* I-HF | 5'-3' AGGAGAAAAAACCCCGGATCCG<br>3'-5' CAACTTCTGTTCCATGTCGAC |
| pESC-HIS (Gal 10) | *Spe* I-HF | 5'-3' CACTAAAGGGCGGCCGCACTAGTA<br>3'-5' CTTGTAATCCATCGATACTAGTGC |
| pESC-LEU (Gal 10) | *Spe* I-HF | 5'-3' ACCCTCACTAAAGGGCGGCCGCAACC<br>3'-5' GTCATCCTTGTAATCCATCGATAC |
| pESC-URA (Gal 1) | *Bam*H I-HF/*Sal* I-HF | 5'-3' AGGAGAAAAAACCCCGGATCCG<br>3'-5' CAACTTCTGTTCCATGTCGAC |
| pOPINF | *Hind* III-HF/*Kpn* I-HF | 5'-3' AAGTTCTGTTTCAGGGCCCG<br>3'-5' ATGGTCTAGAAAGCTTTA |
| pOPINM | *Hind* III-HF/*Kpn* I-HF | 5'-3' AAGTTCTGTTTCAGGGCCCG<br>3'-5' ATGGTCTAGAAAGCTTTA |
| pTRBO | *Bmt* I-HF /*Sap* I-HF | 5'-3' AAGTTCTGTTTCAGGGCCCG<br>3'-5' ATGGTCTAGAAAGCTTTA |

**Table. S19. Protein activities in *S. barbata*, *S. baicalensis* and *S. splendens*.**

| Protein | Gene_id | Species | Activity | NCBI<br>Accession<br>numbers |
| --- | --- | --- | --- | --- |
| **Class II diTPS** |  |  |  |  |
| SbbdiTPS2.1 | Sbb02g0030951 | *Scutellaria barbata* | kolavenyl diphosphate synthase | MT886857 |
| SbbdiTPS2.2 | Sbb06g0014700 | *Scutellaria barbata* | ent-copalyl diphosphate synthase | MT886858 |
| SbbdiTPS2.3 | Sbb02g0030960 | *Scutellaria barbata* | isokolavenyl diphosphate synthase | MT886859 |
| SbbdiTPS2.4 | Sbb02g0014540 | *Scutellaria barbata* | copalyl diphosphate synthase | MT886860 |
| SbbdiTPS2.5 | Sbb02g0014541 | *Scutellaria barbata* | copalyl diphosphate synthase | MT886861 |
| SbbdiTPS2.6 | Sbb02g0014610 | *Scutellaria barbata* | - | MT886862 |
| SbbdiTPS2.7 | Sbb02g0014650 | *Scutellaria barbata* | - | MT886863 |
| SbbdiTPS2.8 | Sbb02g0014660 | *Scutellaria barbata* | - | MT886864 |
| SbdiTPS2.1 | Sb06g19680 | *Scutellaria baicalensis* | copalyl diphosphate synthase | MT909790 |
| SbdiTPS2.2 | Sb06g19670 | *Scutellaria baicalensis* | copalyl diphosphate synthase | MT909791 |
| SbdiTPS2.3 | Sb06g19660 | *Scutellaria baicalensis* | copalyl diphosphate synthase | MT909792 |
| SbdiTPS2.4 | Sb06g19690 | *Scutellaria baicalensis* | - | MT909793 |
| SbdiTPS2.5 | Sb03g27890 | *Scutellaria baicalensis* | ent-copalyl diphosphate synthase | MT909794 |
| SbdiTPS2.6 | Sb01g04220 | *Scutellaria baicalensis* | ent-copalyl diphosphate synthase | MT909795 |
| SbdiTPS2.7 | Sb06g03970 | *Scutellaria baicalensis* | isokolavenyl diphosphate synthase | MT909796 |
| SbdiTPS2.8 | Sb06g05650 | *Scutellaria baicalensis* | isokolavenyl diphosphate synthase | MT909797 |
| SbdiTPS2.9 | Sb08g08190 | *Scutellaria baicalensis* | - | MT909798 |
| SbdiTPS2.10 | Sb06g19710 | *Scutellaria baicalensis* | - | MT909799 |
| SspdiTPS2.1 | Saspl_043012 | *Salvia splendens* | kolavenyl diphosphate synthase | MT909805 |
| SspdiTPS2.2 | Saspl_048790 | *Salvia splendens* | - | MT909806 |
| SspdiTPS2.3 | Saspl_027494 | *Salvia splendens* | ent-copalyl diphosphate synthase | MT909807 |
| SspdiTPS2.4 | Saspl_009166 | *Salvia splendens* | copalyl diphosphate synthase | MT909808 |
| SspdiTPS2.5 | Saspl_017770 | *Salvia splendens* | copalyl diphosphate synthase | MT909809 |
| SspdiTPS2.6 | Saspl_009980 | *Salvia splendens* | ent-copalyl diphosphate synthase | MT909810 |
| SspdiTPS2.7 | Saspl_013955 | *Salvia splendens* | - | MT909811 |
| SspdiTPS2.8 | Saspl_006526 | *Salvia splendens* | - | MT909812 |
| Si011084261 | Si011084261 | *Sesamum indicum* | - | MT909813 |
| **Class I diTPS** |  |  |  |  |
| SbbdiTPS1.1 | Sbb02g0014530 | *Scutellaria barbata* | miltiradiene synthase | MT886853 |
| SbbdiTPS1.2 | Sbb02g0014640 | *Scutellaria barbata* | kolavenol synthase/isokolavenol synthase | MT886854 |
| SbbdiTPS1.3 | Sbb02g0011460 | *Scutellaria barbata* | ent-kaurene synthase | MT886855 |
| SbbdiTPS1.4 | Sbb02g0014620 | *Scutellaria barbata* | kolavenol synthase/isokolavenol synthase | MT886856 |
| SbdiTPS1.1 | Sb06g19650 | *Scutellaria baicalensis* | miltiradiene synthase | MT909787 |
| SbdiTPS1.2 | Sb03g16790 | *Scutellaria baicalensis* | ent-kaurene synthase | MT909788 |
| SbdiTPS1.3 | Sb06g19700 | *Scutellaria baicalensis* | isokolavenol synthase | MT909789 |
| SspdiTPS1.1 | Saspl_006722 | *Salvia splendens* | ent-kaurene synthase/kolavenol synthase | MT909800 |
| SspdiTPS1.2 | Saspl_001373 | *Salvia splendens* | ent-kaurene synthase/kolavenol synthase | MT909801 |
| SspdiTPS1.3 | Saspl_037723 | *Salvia splendens* | miltiradiene synthase/kolavenol synthase | MT909802 |
| SspdiTPS1.4 | Saspl_048791 | *Salvia splendens* | miltiradiene synthase | MT909803 |
| SspdiTPS1.5 | Saspl_037211 | *Salvia splendens* | kolavenol synthase | MT909804 |
| **P450s** |  |  |  |  |
| SbbCYP1 | Sbb02g0014531 | *Scutellaria barbata* | - |  |

|  |  |  |  |
| --- | --- | --- | --- |
| SbbCYP2 | Sbb02g0014542 | *Scutellaria barbata* | ferruginol synthase |
| SbbCYP3 | Sbb02g0014550 | *Scutellaria barbata* | ferruginol synthase |
| SbbCYP4 | Sbb02g0014560 | *Scutellaria barbata* | ferruginol synthase |
| SbbCYP2b | Sbb02g0014570 | *Scutellaria barbata* | ferruginol synthase |
| SbbCYP1b | Sbb02g0014581 | *Scutellaria barbata* | ferruginol synthase |
| SbbCYP3b | Sbb02g0014590 | *Scutellaria barbata* | ferruginol synthase |
| SbbCYP2c | Sbb02g0014600 | *Scutellaria barbata* | ferruginol synthase |
| SbCYP1 | – | *Scutellaria baicalensis* | ferruginol synthase |
| SbCYP2 | – | *Scutellaria baicalensis* | ferruginol synthase |
| SbCYP3 | – | *Scutellaria baicalensis* | ferruginol synthase |
| SbCYP4 | – | *Scutellaria baicalensis* | ferruginol synthase |
| SspCYP1 | Saspl_009167 | *Salvia splendens* | - |
| SspCYP2 | Saspl_009168 | *Salvia splendens* | ferruginol synthase |
| SspCYP3 | Saspl_017768 | *Salvia splendens* | ferruginol synthase |
| SspCYP4 | Saspl_017769 | *Salvia splendens* | ferruginol synthase |

**Table S20. Primers used in this study.**

| Gene name | Primer name | Sequences (5' to 3') |
| --- | --- | --- |
| *Scutellaria barbata* |  |  |
| SbbdiTPS2.1 | pTRBO-F | AAGTTCTGTTTCAGGGCCCGATGGCATTTGCTTGGCAAACAGCC |
|  | pTRBO-R | ATGGTCTAGAAAGCTTTAACTACTTTCTCAAATAGCAC |
| SbbdiTPS2.2 | pTRBO-F | AAGTTCTGTTTCAGGGCCCGATGCCTTTCCTCCTCCCTTCCTC |
|  | pTRBO-R | ATGGTCTAGAAAGCTTTATGAACTCTTTCAAAGAGCAC |
| SbbdiTPS2.3 | pTRBO-F | AAGTTCGTTCAGGGCCCGATGGCATTGCTTGGCAAACAAC |
|  | pTRBO-R | ATGGTCTAGAAAGCTTTAGAGTACTTTCTCAAATAGTAC |
| SbbdiTPS2.4 | pTRBO-F | AAGTTCTGTTTCAGGGCCCGATGGGCTCTCTATCAACTCTAAGC |
|  | pTRBO-R | ATGGTCTAGAAAGCTTTACATGACTGGTTCGAAGAGTAC |
| SbbdiTPS2.4b | pTRBO-F | AAGTTCTGTTTCAGGGCCCGATGGGCTCTCTATCAACTCTAAGC |
|  | pTRBO-R | ATGGTCTAGAAAGCTTTACATGACTGGTTCGAAGAGTAC |
| SbbdiTPS2.5 | pTRBO-F | AAGTTCTGTTTCAGGGCCCGATGGCCTCTTTATCAACTCTG |
|  | pTRBO-R | ATGGTCTAGAAAGCTTTACATGACTGGTTCGAAAAGCAC |
| SbbdiTPS2.6 | pTRBO-F | AAGTTCTGTTTCAGGGCCCGATGTACAGTACTCTCTCAACTC |
|  | pTRBO-R | ATGGTCTAGAAAGCTTTACACAACTTGCCGAAATAGTATC |
| SbbdiTPS2.6b | pTRBO-F | AAGTTCTGTTTCAGGGCCCGATGTACAGTACTCTCTCAACTC |
|  | pTRBO-R | ATGGTCTAGAAAGCTTTACACAACTTGCCGAAATAGTATC |
| SbbdiTPS2.7 | pTRBO-F | AAGTTCTGTTTCAGGGCCCGATGATTTTAGTGAAGACAATCG |
|  | pTRBO-R | ATGGTCTAGAAAGCTTTAAAATACCTGTTATGCATG |
| SbbdiTPS2.8 | pTRBO-F | AAGTTCTGTTTCAGGGCCCGATGTATATTCTCTCAGCTCAAC |
|  | pTRBO-R | ATGGTCTAGAAAGCTTTACACAAGTTGCTCAAAACAGTAC |
| SbbdiTPS1.1 | pTRBO-F | AAGTTCTGTTTCAGGGCCCGATGTCGGCCGGGTTAAACCTC |
|  | pTRBO-R | ATGGTCTAGAAAGCTTTAGTTTTTGCTGCTAATATTATAATTAG |
| SbbdiTPS1.2 | pTRBO-F | AAGTTCTGTTTCAGGGCCCGATGTATTCTCTTAGAGTTTTCTC |
|  | pTRBO-R | ATGGTCTAGAAAGCTTTATGCTGAAAGAGGAAGATGAATAG |
| SbbdiTPS1.3 | pTRBO-F | AAGTTCTGTTTCAGGGCCCGATGTCTCTTCAGCTTTCCAATAG |
|  | pTRBO-R | ATGGTCTAGAAAGCTTTAAAATTCTTTGAGAACAAATAGG |
| SbbdiTPS1.4 | pTRBO-F | AAGTTCTGTTTCAGGGCCCGATGTCCGTAGGATATTCTCTCAG |
|  | pTRBO-R | ATGGTCTAGAAAGCTTTAAGTTGAAATAGGAGGAAGATG |
| SbbCYP1 | pESC-URA-g1 F | AGGAGAAAAAACCCCGGATCCGATGGATCTCTTCACTATTTTC |
|  | pESC-URA-g1 R | CAACTTCTGTTCCATGTCGACTGGTTTGATTGGAATAGCCTTAAG |
| SbbCYP2 | pESC-URA-g1 F | AGGAGAAAAAACCCCGGATCCGATGGAACTATTTACGCTTCTG |
|  | pESC-URA-g1 R | CAACTTCTGTTCCATGTCGACCTTTTTAATTGGGTAAGCC |
| SbbCYP3 | pESC-LEU-g10 F | ACCCTCACTAAAGGGCGGCCGCAACCATGGATACTTATGCAATAGTG |
|  | pESC-LEU-g10 R | GTCATCCTTGTAATCCATCGATACGAGCTTGATTGGAATGAGCTTG |
| SbbCYP4 | pESC-LEU-g10 F | ACCCTCACTAAAGGGCGGCCGCAACCATGGATACTTATGCAATAGTG |
|  | pESC-LEU-g10 R | GTCATCCTTGTAATCCATCGATACGAGCTTGATTGGAATCAGCTTG |
| **Truncated version** |  |  |
| SbbdiTPS2.1(Δ76) | pOPINF-F | AAGTTCTGTTTCAGGGCCCGTTGGAAAAGGAAAAAGGCTTGGAG |
|  | pOPINF-R | ATGGTCTAGAAAGCTTTAACTACTTTCTCAAATAGCAC |
| SbbdiTPS2.3(Δ76) | pOPINF-F | AAGTTCTGTTTCAGGGCCCGTTGGAAAAGGAAAAAGGCTTG |
|  | pOPINF-R | ATGGTCTAGAAAGCTTTAGAGTACTTTCTCAAATAGCAC |
| SbbdiTPS2.4(Δ37) | pOPINF-F | AAGTTCTGTTTCAGGGCCCGTGCATGAACAACAGTAAAAAACTG |
|  | pOPINF-R | ATGGTCTAGAAAGCTTTACATGACTGGTTCGAAGAGTAC |
| SbbdiTPS2.4b(Δ37) | pOPINF-F | AAGTTCTGTTTCAGGGCCCGTGCATGAACAACAGTAAAAAACTG |
|  | pOPINF-R | ATGGTCTAGAAAGCTTTACATGACTGGTTCGAAGAGTAC |
| SbbdiTPS2.5(Δ60) | pOPINF-F | AAGTTCTGTTTCAGGGCCCGGAAATGCAAGTTGCCACTGTGG |
|  | pOPINF-R | ATGGTCTAGAAAGCTTTACATGACTGGTTCGAAAAGCAC |
| SbbdiTPS1.1(Δ41) | pOPINF-F | AAGTTCTGTTTCAGGGCCCGTGCAGCCTCAAGACAGCTTCAAC |
|  | pOPINF-R | ATGGTCTAGAAAGCTTTAGTTTTTGCTGCTAATATTATAATTAG |
| SbbdiTPS1.2(Δ254) | pOPINF-F | AAGTTCTGTTTCAGGGCCCGGTTCTTGAAGCTTGCAATGG |
|  | pOPINF-R | ATGGTCTAGAAAGCTTTATGCTGAAAGAGGAAGATGAATAG |
| SbbdiTPS1.3(Δ49) | pOPINM-F | AAGTTCTGTTTCAGGGCCCGGTGAAGCTCGTCCATAAGG |
|  | pOPINM-R | ATGGTCTAGAAAGCTTTAAAATTCTTTGAGAACAAATAGG |
| SbbdiTPS1.4(Δ256) | pOPINF-F | AAGTTCTGTTTCAGGGCCCGGTTCTTGAAGCTTGCAATGG |
|  | pOPINF-R | ATGGTCTAGAAAGCTTTAAGTTGAAATAGGAGGAAGATG |
| **qRT-PCR** |  |  |
| SbbdiTPS2.1 | F | CTCCTCTCATCCCCCAATCA |
|  | R | CTTGTACTGCCTGAGGGACC |
| SbbdiTPS2.2 | F | TGAACATGGAAGCATCCTCAAA |
|  | R | GCTCTGCTTCGCCTTCTCCA |
| SbbdiTPS2.3 | F | CTCCTCTCATCCCCCAACCG |
|  | R | CTTGTACTGCCTAAGGGACA |
| SbbdiTPS2.4 | F | ATCATCTGCAAAGCCTAAAC |
|  | R | CCAAAGTGGCAACTTGCAGC |
| SbbdiTPS2.5 | F | CAATCATCTGCAAAACTTCAG |
|  | R | CCACAGTGGCAACTTGCATT |
| SbbdiTPS1.1 | F | GCCGGGTTAAACCTCAAAGT |
|  | R | AGCTTTCCCAATGACCTTCTCT |
| SbbdiTPS1.2 | F | GGACTGAAGCATTTCTCAAG |
|  | R | CAATTTGATCTGGTGTGCCA |
| SbbdiTPS1.3 | F | ATGGGGAGTTGGTGAAGAAC |
|  | R | TCTCTGCCGAGTGATATTGGT |
| SbbdiTPS1.4 | F | GGACTAAAGGATTTCTCACT |
|  | R | TAAATCGATCGGGTGTGACG |
| Sbb qACTIN | F | GCTCCTCTCAACCCTAAGGC |
|  | R | GGGGAGAGCATAACCCTCGT |

| *Scutellaria baicalensis* |  |  |
| --- | --- | --- |
| SbdiTPS2.1 | pTRBO-F | AAGTTCTGTTTCAGGGCCCGATGCCCTCTCTCTCCACTCTAAAC |
|  | pTRBO-R | ATGGTCTAGAAAGCTTTATAGGACTGGTTCTAAAAGTAC |
| SbdiTPS2.2 | pTRBO-F | AAGTTCTGTTTCAGGGCCCGATGGCATCTTTATCAACTCT |
|  | pTRBO-R | ATGGTCTAGAAAGCTTTAAAGACGAATTAAATGGTGTTTAG |
| SbdiTPS2.3 | pTRBO-F | AAGTTCTGTTTCAGGGCCCGATGGCCTCTCTATCAACTCTG |
|  | pTRBO-R | ATGGTCTAGAAAAGCTTTACATGACTGGTTCGAAAAGTACT |
| SbdiTPS2.4 | pTRBO-F | AAGTTCTGTTTCAGGGCCCGATGCACACTCTCTCAACTCTG |
|  | pTRBO-R | ATGGTCTAGAAAAGCTTTATACAACCTTGCTCAAATAGTATC |
| SbdiTPS2.5 | pTRBO-F | AAGTTCTGTTTCAGGGCCCCGATGCCTTTCTCTCCTCCCTTCCTC |
|  | pTRBO-R | ATGGTCTAGAAAGCTTTAAAGAACTCTTTCAAAGAGCACCT |
| SbdiTPS2.6 | pTRBO-F | AAGTTCTGTTTCAGGGCCCGATGCCTTTCCTCCTCCCTTCTTC |
|  | pTRBO-R | ATGGTCTAGAAAGCTTTAAAGAACTCTTTCAAAGAGCACCT |
| SbdiTPS2.7 | pTRBO-F | AAGTTCTGTTTCAGGGCCCGATGGCATTTGCTTGGCAAACAAC |
|  | pTRBO-R | ATGGTCTAGAAAGCTTTAGACTACTTTCTCAAATATCACTTT |
| SbdiTPS2.8 | pTRBO-F | AAGTTCTGTTTCAGGGCCCGATGGCATTTGCTTGGCAAACAACC |
|  | pTRBO-R | ATGGTCTAGAAAGCTTTAGACTACTTTCTCAAATATCAC |
| SbdiTPS2.9 | pTRBO-F | AAGTTCTGTTTCAGGGCCCGATGCACACTCTCTCAACTCTGC |
|  | pTRBO-R | ATGGTCTAGAAAGCTTTATACAACTTGCTCAAATAGTATC |
| SbdiTPS2.10 | pTRBO-F | AAGTTCTGTTTCAGGGCCCGATGAAGATCCAATGGATGTTG |
|  | pTRBO-R | ATGGTCTAGAAAGCTTTAGACCACACTTTCAAACAATACTTTG |
| SbdiTPS1.1 | pTRBO-F | AAGTTCTGTTTCAGGGCCCGATGTCGGTCGGCTTAAACCTC |
|  | pTRBO-R | ATGGTCTAGAAAGCTTTATAAACCAATGATTTTCATATCCTTC |
| SbdiTPS1.2 | pTRBO-F | AAGTTCTGTTTCAGGGCCCGATGTCTCTTCAGCTTTCCAGTAG |
|  | pTRBO-R | ATGGTCTAGAAAGCTTTAAAATTCTTTGAGAACAATAGGTTGAT |
| SbdiTPS1.3 | pTRBO-F | AAGTTCTGTTTCAGGGCCCGATGTCCGTAGGATACTCTCTC |
|  | pTRBO-R | ATGGTCTAGAAAGCTTTAATGGATATCTGCAGAATTGCCC |
| SbCYP1 | pESC-LEU-g10 F | ACCCTCACTAAAGGGCGGCCGCAACCATGGAAGTGTTTACGTTTCTGG<br>TA |
|  | pESC-LEU-g10 R | GTCATCCTTGTAATCCATCGATACGGTATTGATAGGGTAAGCCTTG |
| SbCYP2 | pESC-LEU-g10 F | ACCCTCACTAAAGGGCGGCCGCAACCATGGATACTTATGCAATAGTG |
|  | pESC-LEU-g10 R | GTCATCCTTGTAATCCATCGATACGGCCTTGATTGGGATGAGCTTG |
| SbCYP3 | pESC-LEU-g10 F | ACCCTCACTAAAGGGCGGCCGCAACCATGGATACTTATGCAATAG |
|  | pESC-LEU-g10 R | GTCATCCTTGTAATCCATCGATACGGCCTTGATTGGGATGAGCTTG |
| SbCYP4 | pESC-LEU-g10 F | ACCCTCACTAAAGGGCGGCCGCAACCATGGAAGTGTTTACGTTTCTGA<br>TA |
|  | pESC-LEU-g10 R | GTCATCCTTGTAATCCATCGATACGGTATTGATAGGGTAAGCCTTG |

| Truncated version |  |  |
| --- | --- | --- |
| SbdiTPS2.1(Δ32) | pOPINF-F | AAGTTCTGTTTCAGGGCCCCGTGTTTACCATTTCGTACATG |
|  | pOPINF-R | ATGGTCTAGAAAAGCTTTATAGGACTGGTTCTAAAAGTAC |
| SbdiTPS2.2(Δ55) | pOPINF-F | AAGTTCTGTTTCAGGGCCCGACATCGAACGTAACAGAACTG |
|  | pOPINF-R | ATGGTCTAGAAAGCTTTAAAGACGAATTAAATGGTGTTTAG |
| SbdiTPS2.3(Δ38) | pOPINF-F | AAGTTCTGTTTCAGGGCCCGATGAACAACAGTAAAAGACTGTCTTTG |
|  | pOPINF-R | ATGGTCTAGAAAAGCTTTACATGACTGGTTCGAAAAGTAC |
| SbdiTPS2.5(Δ42) | pOPINF-F | AAGTTCTGTTTCAGGGCCCCGTGCAATGCAATCTCTCGACCTC |
|  | pOPINF-R | ATGGTCTAGAAAAGCTTTAAAGAACTCTTTCAAAGAGCAC |
| SbdiTPS2.6(Δ42) | pOPINF-F | AAGTTCTGTTTCAGGGCCCGTGCAATGCAATCTCTCGACCTC |
|  | pOPINF-R | ATGGTCTAGAAAAGCTTTAAAGAACTCTTTCAAAGAGCAC |
| SbdiTPS2.7(Δ76) | pOPINF-F | AAGTTCTGTTTCAGGGCCCGTTGGAAACAGAAAAAGGCTTGG |
|  | pOPINF-R | ATGGTCTAGAAAAGCTTTAGACTACTTTCTCAAATATCAC |
| SbdiTPS2.8(Δ76) | pOPINF-F | AAGTTCTGTTTCAGGGCCCGTTGGAAACAGAAAAAGGCTTGG |
|  | pOPINF-R | ATGGTCTAGAAAAGCTTTAGACTACTTTCTCAAATATCAC |
| SbdiTPS2.9(Δ17) | pOPINF-F | AAGTTCTGTTTCAGGGCCCGGGAAAAATGGTGTTCTCGCCG |
|  | pOPINF-R | ATGGTCTAGAAAAGCTTTATACAACCTTGCTCAAATAGTATC |
| SbdiTPS1.1(Δ41) | pOPINF-F | AAGTTCTGTTTCAGGGCCCGTGCAGCCTCAAGACAGCTTCAAC |
|  | pOPINF-R | ATGGTCTAGAAAGCTTTATAAACCAATGATTTTCATATC |
| SbdiTPS1.2(Δ46) | pOPINF-F | AAGTTCTGTTTCAGGGCCCGGCAAGAATTGCGAAGCTGGTTCATAAG |
| SbdiTPS1.3(Δ43) | pOPINF-R<br>pOPINF-F<br>pOPINF-R | ATGGTCTAGAAAGCTTTAAAATTCTTTGAGAACAATAGGTTGAT<br>AGGAGAAAAACCCCGGATCCGTCACGAATAATGAGAAAGTTGATTG<br>ATGGTCTAGAAAGCTTTAATGGATATCTGCAGAATTGCC |
| **Salvia splendens** |  |  |
| SspdiTPS2.1 | pTRBO-F<br>pTRBO-R | AAGTTCTGTTTCAGGGCCCGATGAGTATTCAAGCAAACATGTCG<br>ATGGTCTAGAAAGCTTTAGACAATTTTTTCAAACAATACTTTGTTTATG<br>T |
| SspdiTPS2.2 | pTRBO-F<br>pTRBO-R | AAGTTCTGTTTCAGGGCCCGATGTACATCCTTTCAACTCCA<br>ATGGTCTAGAAAGCTTTACATAACCTGCTCGAACAGTA |
| SspdiTPS2.3 | pTRBO-F<br>pTRBO-R | AAGTTCTGTTTCAGGGCCCGATGTCCCTCGCTTCCAATC<br>ATGGTCTAGAAAGCTTTAATGTACTCTTTTGAAGAGCACTT |
| SspdiTPS2.4 | pTRBO-F<br>pTRBO-R | AAGTTCTGTTTCAGGGCCCGATGGCCTCTCTTCTCTTACA<br>ATGGTCTAGAAAGCTTTACTCGACTGGTTCGAAAAGC |
| SspdiTPS2.5 | pTRBO-F<br>pTRBO-R | AAGTTCTGTTTCAGGGCCCGATGGCCTCTCTTCTCTTACA<br>ATGGTCTAGAAAGCTTTACTCGACTGGTTCGAAAAGCA |
| SspdiTPS2.6 | pTRBO-F<br>pTRBO-R | AAGTTCTGTTTCAGGGCCCGATGGCCTCTCTTCTCTATC<br>ATGGTCTAGAAAGCTTTATTGTACTCTTTCAAAGAGTACTTTC |
| SspdiTPS2.7 | pTRBO-F<br>pTRBO-R | AAGTTCTGTTTCAGGGCCCGATGTCTACTCTAAATTTGAGCAC<br>ATGGTCTAGAAAGCTTTATACAGCCGGTTCAAACAGTACTT |
| SspdiTPS2.8 | pTRBO-F<br>pTRBO-R | AAGTTCTGTTTCAGGGCCCGATGTACATTCTTTCAACTCCAC<br>ATGGTCTAGAAAGCTTTACAGCTTATGATTTTCAAGCTTT |
| SspdiTPS1.1 | pTRBO-F<br>pTRBO-R | AAGTTCTGTTTCAGGGCCCGATGTCGCTTCTCTCTCCACTTG<br>ATGGTCTAGAAAGCTTTACAACATTTTCTCTTTGAGAAGAAGAGG |
| SspdiTPS1.2 | pTRBO-F<br>pTRBO-R | AAGTTCTGTTTCAGGGCCCGATGTCGCTTCTCTCTCCACT<br>ATGGTCTAGAAAGCTTTACATCTTTTGGTCTTGAGAAGAATAGG |
| SspdiTPS1.3 | pTRBO-F<br>pTRBO-R | AAGTTCTGTTTCAGGGCCCGATGTCGGCCACCTTCAA<br>ATGGTCTAGAAAGCTTTACATTGCTCTCAACATTATTAGG |
| SspdiTPS1.4 | pTRBO-F<br>pTRBO-R | AAGTTCTGTTTCAGGGCCCGATGTCGCTCGCCTTCAATC<br>ATGGTCTAGAAAGCTTTACTTGCCACTCACATTATTAGCTTC |
| SspdiTPS1.5 | pTRBO-F<br>pTRBO-R | AAGTTCTGTTTCAGGGCCCGATGTCGCTCGCCTTCAAC<br>ATGGTCTAGAAAGCTTTATTAACCTAGGAAGTGTAAGAGGTTTATATAT |
| SspCYP1 | pESC-LEU-g10 F<br>pESC-LEU-g10 R | ACCCTCACTAAAGGGCGGCCGCAACCATGGAGTTGTCCACTGTTGC<br>GTCATCCTTGTAATCCATCGATACATGGCTAAGGGGATAAGCC |
| SspCYP2 | pESC-LEU-g10 F<br>pESC-LEU-g10 R | ACCCTCACTAAAGGGCGGCCGCAACCATGGACTCCTTCCCTTTCC<br>GTCATCCTTGTAATCCATCGATACATGGCTTAAATGGGATGACCT |
| SspCYP3 | pESC-LEU-g10 F<br>pESC-LEU-g10 R | ACCCTCACTAAAGGGCGGCCGCAACCATGGACTCCTTCCCTTTCC<br>GTCATCCTTGTAATCCATCGATACATGGCTTAAATGGGATGACCC |
| SspCYP4 | pESC-LEU-g10 F<br>pESC-LEU-g10 R | ACCCTCACTAAAGGGCGGCCGCAACCATGGAGTTGTCCACTCTTGC<br>GTCATCCTTGTAATCCATCGATACATGGCTATGGGGATAAGC |
| **Truncated version** |  |  |
| SspdiTPS2.1Δ72 | pOPINF-F<br>pOPINF-R | AAGTTCTGTTTCAGGGCCCGTCAACAATTTTGGAGGGACAAAC<br>ATGGTCTAGAAAGCTTTAGACAATTTTTTCAAACAATACTTTGTTTATG<br>T |
| SspdiTPS2.3Δ70 | pOPINF-F<br>pOPINF-R | AAGTTCTGTTTCAGGGCCCGACAAAGATCGCCGGGC<br>ATGGTCTAGAAAGCTTTAATGTACTCTTTTGAAGAGCACTT |
| SspdiTPS2.4Δ65 | pOPINF-F<br>pOPINF-R | AAGTTCTGTTTCAGGGCCCGGCTCCACAGGTGCATGATC<br>ATGGTCTAGAAAGCTTTACTCGACTGGTTCGAAAAGC |
| SspdiTPS2.5Δ61 | pOPINF-F<br>pOPINF-R | AAGTTCTGTTTCAGGGCCCGGCTCCACAGGTGCAAGATC<br>ATGGTCTAGAAAGCTTTACTCGACTGGTTCGAAAAGCA |
| SspdiTPS1.1Δ55 | pOPINF-F<br>pOPINF-R | AAGTTCTGTTTCAGGGCCCGGCAAAGCTGTTTCATAAGGATGAAC<br>ATGGTCTAGAAAGCTTTACAACATTTTCTCTTTGAGAAGAAGAGG |
| SspdiTPS1.2Δ55 | pOPINF-F<br>pOPINF-R | AAGTTCTGTTTCAGGGCCCGGCAAAGCTGTTTCATAGCAATGAAC<br>ATGGTCTAGAAAGCTTTACATCTTTTGGTCTTGAGAAGAATAGG |
| SspdiTPS1.3Δ61 | pOPINF-F<br>pOPINF-R | AAGTTCTGTTTCAGGGCCCGGAAAAACAGTAATTTTCCGGTCACTTTT<br>ATGGTCTAGAAAGCTTTACATTGCTCTCAACATTATTAGG |
| SspdiTPS1.4Δ61 | pOPINF-F<br>pOPINF-R | AAGTTCTGTTTCAGGGCCCGGAAAAACAGTCATTTTCCGGTCACT<br>ATGGTCTAGAAAGCTTTACTTGCCACTCACATTATTAGCTTC |
| SspdiTPS1.5Δ50 | pOPINF-F<br>pOPINF-R | AAGTTCTGTTTCAGGGCCCGTTGGGGGAATTAAGGACAAAGTT<br>ATGGTCTAGAAAGCTTTATTAACCTAGGAAGTGTAAGAGGTTTATATAT |
| SspdiTPS1.3Δ61+SspdiTPS2.4 | pESC-HIS-g10 F<br>adapter F<br>adapter R<br>pESC-HIS-g10 R | CACCTAAAGGGCGGCCGCACTAGTAATGGAACAGTAATTTTCCGGTC<br>ACTTTT<br>GTTGAGAGGCAAATGGGTGGTGGTTCTATGGCC<br>GGCCATAGAACACCACCCATTTGCCTCTCAAC<br>CTTGTAATCCATCGATACTAGTGCCTCGACTGGTTCGAAAAGC |
| **Other species** |  |  |
| Salvia miltiorrhiza | SmCYP76AH1 pESC-LEU-g10 F<br>tan | ACCCTCACTAAAGGGCGGCCGCAACCATGGATTCTTTTCTCTCTCTC<br>GTCATCCTTGTAATCCATCGATACAGACTTAACTATTGGGATAATC |
| *Tripterygium wilfordii* | TwTPS14 pTRBO F | AAGTTCTGTTTCAGGGCCCGATGTTCATGTCCTCCTCCTCC |
|  | TwTPS14 pTRBO R | ATGGTCTAGAAAGCTTTATACTACTCTTTCAAAGAGTAC |
| *Ajuga reptans* | ArTPS2 pTRBO F | AAGTTCTGTTTCAGGGCCCGATGTCATTTGCTTCCAAGCC |
|  | ArTPS2 pTRBO R | ATGGTCTAGAAAGCTTTAAACCACTTTTTCGAACAGAACCTTATCGATG |
| *Hyssopus officinalis* | Ho4680 pTRBO F | AAGTTCTGTTTCAGGGCCCGTCATTTGTTACCAACACCAC |
|  | Ho4680 pTRBO R | ATGGTCTAGAAAGCTTTAAACAATTCTTCCAAACAATAC |
|  | Ho4684 pTRBO F | AAGTTCTGTTTCAGGGCCCGTCATTTGTTACCAACACCAC |
|  | Ho4684 pTRBO R | ATGGTCTAGAAAGCTTTAAACTATTCTTCCAAACAATAC |
| *Sesamum indicum* | Si011084261 pTRBO F | AAGTTCTGTTTCAGGGCCCGTCAGTAATTGCTTCCACA |
|  | Si011084261 pTRBO R | ATGGTCTAGAAAGCTTTATGTGCATATTACTCTATCAAAGAGC |

**Data S1. (separate file)**

Extension of Table S1 with list of clerodane diterpenoids isolated from *Scutellaria barbata* (with literature references) and their reported cytotoxic activities.

**Data S2. (separate file)**

Extension of Table S15 with list of clerodane diterpenoids and their structures isolated from *Salvia splendens* (with literature references).

### References

- Ahmad, V.U., Farooq, U., Abbaskhan, A., Hussain, J., Abbasi, M.A., Nawaz, S.A., and Choudhary, M.I. (2004). Four new diterpenoids from *Ballota limbata*. *Helvetica chimica acta* **87**:682-689.
- Ahmad, V.U., Khan, A., Farooq, U., Kousar, F., Khan, S.S., Nawaz, S.A., Abbasi, M.A., and Choudhary, M.I. (2005). Three new cholinesterase-inhibiting cis-clerodane diterpenoids from *Otostegia limbata*. *Chemical and pharmaceutical bulletin* **53**:378-381.
- Andersen-Ranberg, J., Kongstad, K.T., Nielsen, M.T., Jensen, N.B., Pateraki, I., Bach, S.S., Hamberger, B., Zerbe, P., Staerk, D., Bohlmann, J., et al. (2016). Expanding the Landscape of Diterpene Structural Diversity through Stereochemically Controlled Combinatorial Biosynthesis. *Angew Chem Int Ed Engl* **55**:2142-2146. 10.1002/anie.201510650.
- Ávila, F., Pinto, F., Sousa, T., Torres, M.C., Costa-Lotufo, L., Rocha, D., de Vasconcelos, M., Cardoso-Sá, N., Teixeira, E., Albuquerque, M.R., et al. (2016). Miscellaneous Diterpenes from the Aerial Parts of *Plectranthus ornatus* Codd. *Journal of the Brazilian Chemical Society* **10**.21577/0103-5053.20160255.
- Borges, C.M., de Mendonça, D.I., Pinheiro, S., Vieira, L., Mendonça, A.J., Gaspar, J.F., Martins, C., Diakanawma, C., and Rueff, J. (2013). New neo-clerodanes from *Tinnea antiscorbutica* Welw. *Journal of the Brazilian Chemical Society* **24**:1950-1956.
- Božić, D., Papaefthimiou, D., Brückner, K., de Vos, R.C., Tsoleridis, C.A., Katsarou, D., Papanikolaou, A., Pateraki, I., Chatzopoulou, F.M., Dimitriadou, E., et al. (2015). Towards Elucidating Carnosic Acid Biosynthesis in Lamiaceae: Functional Characterization of the Three First Steps of the Pathway in *Salvia fruticosa* and *Rosmarinus officinalis*. *PLoS One* **10**:e0124106. 10.1371/journal.pone.0124106.
- Brieskorn, C.H., and Stehle, T. (1973). Labiaten - Bitterstoffe: Eine neue Verbindung des Clerodantyps. *Chemische Berichte* **106**:922-928.
- Castro, A., and Coll, J. (2008). Neo-Clerodane Diterpenoids from Verbenaceae: Structural Elucidation and Biological Activity. *Natural Product Communications* **3**:1934578X0800300630.
- Chen, T.B., Galinis, D.L., and Wiemer, D.F. (1992). Cornutin A and B: novel diterpenoid repellents of leafcutter ants from *Cornutia grandifolia*. *The Journal of Organic Chemistry* **57**:862-866.
- Chin, Y.-W., Jones, W.P., Mi, Q., Rachman, I., Riswan, S., Kardono, L.B., Chai, H.-B., Farnsworth, N.R., Cordell, G.A., and Swanson, S.M. (2006). Cytotoxic clerodane diterpenoids from the leaves of *Premna tomentosa*. *Phytochemistry* **67**:1243-1248.
- Chow, S.Y., Williams, H.J., Huang, Q., Nanda, S., and Scott, A.I. (2005). Studies on taxadiene synthase: interception of the cyclization cascade at the isocembrene stage with GGPP analogues. *J Org Chem* **70**:9997-10003. 10.1021/jo0517489.
- Cuadrado, M.J.S., María, C., Rodríguez, B., Bruno, M., Piozzi, F., and Savona, G. (1991). Neo-clerodane diterpenoids from *Teucrium oxylepis* subsp. *marianum*. *Phytochemistry* **30**:4079-4082.
- Fazio, C., Passannanti, S., Paternostro, M.P., and Piozzi, F. (1992). Neo-clerodane diterpenoids from *Stachys rosea*. *Phytochemistry* **31**:3147-3149.
- Harada, N., and Uda, H. (1978). Absolute stereochemistries of 3-epicaryoptin, caryoptin, and clerodin as determined by chiroptical methods. *Journal of the American Chemical Society* **100**:8022-8024.

**Hu, H., Cao, H., Jian, Y., Zheng, X., and Liu, J.** (2008). Two new clerodane diterpenoid glucosides and other constituents from the roots of *Elsholtzia bodinieri* Van 't.

**Hussain, J., Jamila, N., Khan, F.U., Devkota, K.P., Shah, M.R., and Anwar, S.** (2009). Nepetalan and nepetanoate: a new diterpene aldehyde and a benzene derivative ester from *Nepeta juncea*. *Magn Reson Chem* **47**:625-627. 10.1002/mrc.2439.

**Hussain, J., Ullah, R., Khan, A., Khan, F.U., Muhammad, Z., and Shah, M.R.** (2011). Phlomeoic acid: a new diterpene from *Phlomis bracteosa*. *Nat Prod Commun* **6**:171-173.

**Ignea, C., Ioannou, E., Georgantea, P., Triikka, F.A., Athanasakoglou, A., Loupassaki, S., Roussis, V., Makris, A.M., and Kampranis, S.C.** (2016). Production of the forskolin precursor 11β-hydroxy-manoyl oxide in yeast using surrogate enzymatic activities. *Microb Cell Fact* **15**:46. 10.1186/s12934-016-0440-8.

**Jefferies, P., Knox, J., and Scaf, B.** (1973). Structure elucidation of some ent-clerodane diterpenes from *Dodonaea boroniaefolia* and *Cyanostegia augustifolia*. *Australian Journal of Chemistry* **26**:2199-2211.

**Jia, M., Mishra, S.K., Tufts, S., Jernigan, R.L., and Peters, R.J.** (2019). Combinatorial biosynthesis and the basis for substrate promiscuity in class I diterpene synthases. *Metab Eng* **55**:44-58. 10.1016/j.ymben.2019.06.008.

**Johnson, S.R., Bhat, W.W., Bibik, J., Turmo, A., Hamberger, B., and Hamberger, B.** (2019). A database-driven approach identifies additional diterpene synthase activities in the mint family (Lamiaceae). *J Biol Chem* **294**:1349-1362. 10.1074/jbc.RA118.006025.

**Kikuchi, T., Tsubono, K., Kadota, S., Kizu, H., Imoto, Y., and Tomimori, T.** (1987). Structures of scuterivulactone C1 and C2 by two-dimensional NMR spectroscopy. New clerodane type diterpenoids from *Scutellaria rivularis* Wall. *Chemistry Letters* **16**:987-990.

**Kumar, S., Stecher, G., Suleski, M., and Hedges, S.B.** (2017). TimeTree: A Resource for Timelines, Timetrees, and Divergence Times. *Mol Biol Evol* **34**:1812-1819. 10.1093/molbev/msx116.

**Li, H., Liu, Y., Qin, H., Lin, X., Tang, D., Wu, Z., Luo, W., Shen, Y., Dong, F., Wang, Y., et al.** (2020). A rice chloroplast-localized ABC transporter ARG1 modulates cobalt and nickel homeostasis and contributes to photosynthetic capacity. *New Phytol* **228**:163-178. 10.1111/nph.16708.

**Liu, Z.-K., Wu, D.-R., Shi, Y.-M., Zeng, T., Liu, S.-H., Du, X., Dang, Y.-J., Xiao, W.-L., and Sun, H.-D.** (2014). Three new diterpenoids from *Leonurus japonicus*. *Chinese Chemical Letters* **25**:677-679.

**Macías, F.A., Lacret, R., Varela, R.M., Nogueiras, C., and Molinillo, J.M.** (2010). Isolation and phytotoxicity of terpenes from *Tectona grandis*. *Journal of Chemical Ecology* **36**:396-404.

**Meisel, L., Fonseca, B., González, S., Baeza-Yates, R., Cambiazo, V., Campos, R., González, M., Orellana, A., Retamales, J., and Silva, H.** (2005). A rapid and efficient method for purifying high quality total RNA from peaches (*Prunus persica*) for functional genomics analyses. *Biol Res* **38**:83-88. 10.4067/s0716-97602005000100010.

**Peters, R.J., Ravn, M.M., Coates, R.M., and Croteau, R.B.** (2001). Bifunctional abietadiene synthase: free diffusive transfer of the (+)-copalyl diphosphate intermediate between two distinct active sites. *J Am Chem Soc* **123**:8974-8978. 10.1021/ja010670k.

**Rasikari, H.L., Leach, D.N., Waterman, P.G., Spooner-Hart, R.N., Basta, A.H., Banbury, L.K., Winter, K.M., and Forster, P.I.** (2005). Cytotoxic clerodane diterpenes from *Glossocarya calcicola*. *Phytochemistry* **66**:2844-2850.

**Rasyid, F.A., Fukuyoshi, S., Ando, H., Miyake, K., Atsumi, T., Fujie, T., Saito, Y., Goto, M., Shinya, T., and Mikage, M.** (2017). A novel clerodane diterpene from *Vitex cofassus*. *Chemical and Pharmaceutical Bulletin* **65**:116-120.

**Rijo, P., Gaspar-Marques, C., Simoes, M.F., Duarte, A., Del Carmen Aprea-Rojas, M., Cano, F.H., and Rodriguez, B.** (2002). Neoclerodane and labdane diterpenoids from *Plectranthus ornatus*. *J Nat Prod* **65**:1387-1390. 10.1021/np020203w.

**SHIMOMURA, H., SASHIDA, Y., and OGAWA, K.** (1989). Neo-clerodane diterpenes from *Ajuga nipponensis*. *Chemical and pharmaceutical bulletin* **37**:354-357.

**Tomlinson, M., Man, Z., Elaine, J.B., Jie, L., Haixiu, L., Juri, F., Lionel, H., Gerhard, S., Martin, R., Dongfeng, Y., et al.** (2022). Diterpenoids from *Scutellaria barbata* induce tumour-selective cytotoxicity by taking the brakes off apoptosis. *Medicinal Plant Biology* **1**:1-16. 10.48130/MPB-2022-0003.

**Ye, D., Shu - Lin, P., Qiang, Z., Xun, L., and Li - Sheng, D.** (2002). Clerodane diterpenoids from *Kinostemon alborubrum*. *Helvetica chimica acta* **85**:2547-2552.

**Yuan, W., Jing, L., Qi, W., Shang, K., De-Bing, P., Zhang, R.-H., Xiao-Li, L., Xiao-Chang, D., Zhang, X.-J., and Wei-Lie, X.** (2019). Clerodane diterpenoids with potential anti-inflammatory activity from the leaves and twigs of *Callicarpa cathayana*. *Chinese journal of natural medicines* **17**:953-962.

**Zhang, R.-T., Feng, T., Cai, X.-H., and Luo, X.-D.** (2009). Two new clerodane-type diterpenoids from *Gomphostemma microdon*. *Zeitschrift für Naturforschung B* **64**:443-446.

**Zhao, Q., Yang, J., Cui, M.Y., Liu, J., Fang, Y., Yan, M., Qiu, W., Shang, H., Xu, Z., Yidiresi, R., et al.** (2019). The Reference Genome Sequence of *Scutellaria baicalensis* Provides Insights into the Evolution of Wogonin Biosynthesis. *Mol Plant* **12**:935-950. 10.1016/j.molp.2019.04.002.
